# A Multimodal Single-Cell Epigenomic and 3D Genome Atlas of the Human Basal Ganglia

**DOI:** 10.64898/2026.02.12.705594

**Authors:** Wubin Ding, Amit Klein, Cindy Tatiana Báez-Becerra, Jonathan A. Rink, Anna Bartlett, Ruoxuan Wang, Qiurui Zeng, Rosa Gomez Castanon, Joseph R. Nery, Emma Osgood, William Owens, Alaina Petrella, Chumo Chen, Andrea Saldana Acerbo, Ariana S. Barcoma, Jiayi Liu, Kaitlyn G. Russo, Kyle W. Knutson, Carissa K. Young, Jackson K. Willier, Cesar Barragan, Jessica Arzavala, Silvia Cho, Jordan Altshul, Derek Chan, Eshaan Soma, Jammy Luo, Manya Jain, Sarah Velazquez, Guha V. Sundaram, Austin C. Manning, Yareli Sanchez, Aleksandra Bikkina, Yuanyuan Fu, Kaushik Komandur, Ethan Doan, Shuai Fu, Carolyn O’Connor, Michelle Liem, Mikayla V. Marrin, Cynthia Rose, Shane N. Alt, Jillian Berry, Joseph Colin Kern, Eric Boone, Wei Tian, Huamin Chen, Yang Xie, Yue Wu, Manoj Hariharan, Kai Li, Lei Chang, Nelson Johansen, Wenjin Zhang, Zoey Zhao, Jesus Flores, Chu-Yi Tai, Jacqueline Olness, Natalie Schenker-Ahmed, Xiao-Ping Liu, Quan Zhu, Brian E. Kalmbach, Rebecca D. Hodge, Trygve E. Bakken, Ed S. Lein, Daofeng Li, Ting Wang, Xiangmin Xu, Bing Ren, Maria Margarita Behrens, Joseph R. Ecker

**Affiliations:** Genomic Analysis Laboratory, The Salk Institute for Biological Studies, La Jolla, CA 92037, USA; Bioinformatics and Systems Biology Program, University of California, San Diego, La Jolla, CA, USA; Computational Neurobiology Laboratory, The Salk Institute for Biological Studies, La Jolla, CA 92037, USA; Division of Biological Sciences, University of California, San Diego, La Jolla, CA, USA; Allen Institute for Brain Science, Seattle, WA, USA; Flow Cytometry Core Facility, The Salk Institute for Biological Studies, La Jolla, CA 92037, USA; Department of Pathology and Laboratory Medicine, University of California, Irvine, CA, USA; Center for Epigenomics, Department of Cellular and Molecular Medicine, University of California, San Diego, La Jolla, CA 92093, USA; New York Genome Center, New York, NY, USA; Department of Cellular and Molecular Medicine, University of California, San Diego, La Jolla, CA, USA; Department of Genetics, The Edison Family Center for Genome Sciences and Systems Biology, Washington University School of Medicine, St. Louis, MO 63110, USA; Department of Neurobiology and Biophysics, University of Washington, Seattle, WA 98195, USA; Department of Anatomy and Neurobiology, School of Medicine, University of California, Irvine, Irvine, CA 92697, USA; Department of Microbiology and Molecular Genetics, School of Medicine, University of California, Irvine, Irvine, CA 92697, USA; Department of Biomedical Engineering, University of California, Irvine, Irvine, CA 92697, USA; Department of Computer Science, University of California, Irvine, Irvine, CA 92697, USA; Center for Neural Circuit Mapping, University of California, Irvine, Irvine, CA 92697, USA; Institute for Genomic Medicine, University of California, San Diego, La Jolla, CA, USA; Moores Cancer Center, University of California, San Diego, La Jolla, CA, USA; Department of Genetics and Development, Systems Biology, Biochemistry and Molecular Biophysics, Columbia University Irving Medical Center, New York, NY, USA

## Abstract

The basal ganglia (BG) underlie motor control, reward processing, and many neurological and psychiatric disorders, but a comprehensive epigenomic and 3D-genome atlas of the human BG is lacking. Here we present a multimodal single-cell atlas profiling DNA methylation and 3D chromatin conformation in 261,331 nuclei (snm3C-seq) across eight subregions, resolving 12 classes, 31 subclasses, and 59 groups. Harmonized under the HMBA basal-ganglia consensus taxonomy, this atlas integrates with matched RNA, ATAC-seq, and histone-modification data across five regulatory layers. We identify millions of cell-type- and region-specific differentially methylated regions enriched for distinct transcription factor motifs and link them to disease-associated heritability. Neuron-specific loops dominate cell-type-specific 3D contact remodeling, while most non-neuron-specific loops are constitutive. Among spiny projection neurons (SPNs), chromatin loops, rather than TAD boundaries, distinguish D1, D2, and eccentric SPN subclasses, with eccentric SPNs showing the most loop-level reorganization among the three. We characterize STR D2 SMYD2-HTR7 SPN, a newly recognized POU6F2⁺ D2-SPN subtype, and reveal region-specific methylation and contact gradients of disease-associated genes, including CADM1 and PDE8B. Integrative gene-regulatory networks reconstruct cell-type-resolved enhancer-promoter links to interpret Parkinson’s disease risk variants at SNCA. Finally, MERFISH spatial profiling combined with cross-species Patch-seq identifies non-SPN neuronal subtypes, including a MOXD1⁺ striosomal STR FS PTHLH-PVALB GABA subtype with distinct electrophysiology, partitioning across the striatal matrix-striosome boundary.

**Highlights:**

- A multimodal single-cell atlas maps DNA methylation and 3D genome architecture across human basal ganglia cell types and subregions.
- Neuron-specific loops dominate cell-type-specific 3D contact remodeling in the human BG, whereas most non-neuron-specific loops are constitutive.
- Spiny Projection Neuron (SPN) subtypes exhibit regionally organized epigenomic and 3D genome signatures that align with dorsal-ventral identities.
- Chromatin loops are the primary distinguishing feature among D1, D2, and eccentric SPN subclasses, with eccentric SPNs being the most 3D-reorganized.
- Integrated regulatory maps link cell-type-specific enhancers to disease-associated genetic risk in the human basal ganglia.
- Non-SPN interneurons differ in distribution and in transcriptional and epigenetic identity across the matrix-striosome boundary.

## Introduction

The brain structures that make up the human Basal Ganglia (BG) are integral to our ability to regulate cognitive-motor functions^1^ and to execute action-reward processing^2^. BG dysfunctions include Parkinson’s disease (PD), dyskinesias, addiction disorders, Tourette syndrome, Huntington’s disease, and forms of eating disorders such as anorexia nervosa and bulimia nervosa^3–8^. Disorders involving lesions to BG regions often manifest as cell-type-specific cell death, leading to an imbalance in the striatocortical circuitry^7^. Delineating the exact cell types susceptible to each disorder is therefore an important step in defining the molecular mechanisms underlying disease onset and in refining therapeutics to target only the affected cell types^9^. A major challenge in identifying these disease-associated cell types is the substantial heterogeneity within the BG.

The BG are a group of deep interconnected nuclei, which include the Striatum (Caudate [Ca], Putamen [Pu], and Nucleus Accumbens [NAC]), Globus Pallidus (GP) external (GPe) and internal (GPi) segments, Subthalamic nucleus (STH), and Substantia Nigra (SN) pars compacta (SNc) and pars reticulata (SNr). The predominant cell type in the Striatum (the input node of the BG^10^) is the Spiny Projection Neuron (SPN)^11^, which is synonymous with the Medium Spiny Neuron (MSN). SPNs differentially express dopamine receptors D1 and D2 depending on their involvement in the direct (striatum - GPi/SNr - thalamus) and indirect (striatum - GPe - STH - GPi/SNr - thalamus) pathways^12^, respectively^13^. SPNs can be further subdivided based on their location along the dorsal-ventral brain axis, or whether they are in a matrix or striosome (also called patch) compartment. In addition to the SPNs, various interneuron (INT) types populate the BG and help integrate afferent information, including excitatory signals from the cerebral cortex and the thalamus^14,15^. Dopaminergic outputs from the SNc innervate the BG and modulate neuronal activity and synaptic plasticity^16,17^. Neurons of the GP, including the GPi, which acts as an output node of the BG, exhibit distinct transcriptional properties and connectivity patterns^18^. GABAergic projections neurons of the SNr, another output node of the BG, utilize specific molecular machinery to tonically inhibit downstream targets in the thalamus, superior colliculus, and other midbrain structures^19^, while also inhibiting SNc dopaminergic and other SNr GABAergic neurons via axon collaterals^20^. Lastly, it has recently been shown that non-neuronal cell types, such as astrocytes, also exhibit regional specificity within the mouse striatum^21^. While previous works have attempted to characterize individual cell groups within the BG^11,14,18,22^, no comprehensive atlas of all cell types in the human BG exists to date. As part of the BRAIN Initiative Cell Atlas Network (BICAN, RRID: SCR_022794), we sought to characterize this cell-type heterogeneity and generate a harmonized cell-type taxonomy, with a focus on DNA methylation patterns and 3D genome structure across BG cells.

Cytosine Methylation (5mC) is the covalent addition of a methyl group onto the 5th Carbon of Cytosine. It is an important DNA modification involved in epigenetic regulation of cell differentiation, with widespread implications for development^23–25^, disease^24,26^, and aging^27,28^. 5mC is often classified into CpG (mCG) and non-CpG (mCH, where H is A, C, or T) due to their distinct regulatory functions. mCG is prevalent in all human tissues, whereas mCH is most abundant in post-mitotic cells such as neurons^29^. Furthermore, the 3D organization of the genome is an important regulator of transcriptional activity. The 3D genome is often viewed at 3 scales: genomic compartments divide chromatin into active (A) and inactive (B) regions; topologically associating domains (TADs) are contiguous DNA blocks enriched for internal interactions; and chromatin loops form specific contacts between regulatory elements, such as enhancer-promoter interactions^30,31^. Previous studies have shown that coupling DNA methylation and genomic 3D contact information with transcriptional cell-type atlases provides novel insights into the epigenetic control of transcriptional cell states^32,33^. Epigenetic atlases enable the use of genomic techniques to map DNA mutations onto cell-type-specific transcriptional regulatory pathways implicated in developmental and neurological disorders. Additionally, due to the transient nature of RNA transcription, covalent 5mC modifications serve as a more stable cell-type-specific marker to target^34^. Therefore, a comprehensive atlas of the epigenomes of cell types in the human BG represents an essential companion to the expression-based cell type atlas presented in the companion paper of this package^35^, together providing a multi-modal framework for studying cell-type identity in the human BG.

In this study, we utilize snm3C-seq to characterize the joint DNA methylomes and 3D interactomes of single cells across all regions of the human BG. We augment the transcriptionally defined taxonomy with methylation-specific cell types that cannot be differentiated solely by RNA expression. Then, using the integrated taxonomy, we perform hierarchical differential methylation analyses from the subclass to the group level, identifying differentially methylated genes (DMGs) and regions (DMRs) that show unique marker patterns and transcription factor (TF) enrichment. We further expand on previous work showing global differences in compartmentalization between neuronal and non-neuronal cell types, while demonstrating how 3D genome dynamics confer transcriptional dynamics in specific cell types. Across SPN subtypes, we highlight regional and cell-type-specific coupling among methylation, 3D conformation, and gene expression, and we show variation across the 3 major SPN axes. By harmonizing our DNA methylation and HiC data under the Human and Mammalian Brain Atlas (HMBA) BG taxonomy^35^ with a donor-matched histone modification dataset (Chang et al., co-submit), and with the 10X Multiome (RNA and ATAC from HMBA), we linked DMR-associated TF motifs to ABC-predicted enhancer-promoter interactions and constructed cell-type-specific gene regulatory networks. We then linked those networks to known disease-associated mutations to illustrate cell-type-specific susceptibility. Finally, by integrating our dataset with newly generated multiplexed error-robust fluorescence in situ hybridization (MERFISH) experiments covering the entire BG, we show how imputed spatial information can inform which axes of variance are biologically relevant, as in the case of matrix-striosome specific subtypes in the striatum.

## Results

### Single-cell DNA methylation and 3D chromatin interaction atlas of human basal ganglia

We applied snm3C-seq to simultaneously profile DNA methylation and 3D chromatin conformation across eight neuron-enriched dissected subregions of human BG (Figure 1A-B, S1A and STAR Methods), including head of caudate (CaH), body of caudate (CaB), tail of caudate (CaT), putamen (Pu), nucleus accumbens (NAC), globus pallidus (GP), subthalamic nucleus (STH) and gray matter of midbrain (MGM1). GP is a mixture of the external segment of the globus pallidus (GPe), the internal segment of globus pallidus (GPi) and a small fraction of the ventral pallidus (VeP). MGM1 is a mixture of the ventral tegmental region of the midbrain (VTR), the substantia nigra (SN) and the red nucleus (RN). We performed iterative clustering using 100kb mCG and mCH embedding (STAR Methods) and integrated these newly generated datasets (197,003 cells) with our previously published Human Brain Atlas (HBA)^33^ BG dataset (64,328 cells), resulting in a comprehensive joint epigenomic atlas of 261,331 high-quality cells from the BG regions of seven healthy adult human brains (see Table S1 for sample metadata).

**Figure 1.**
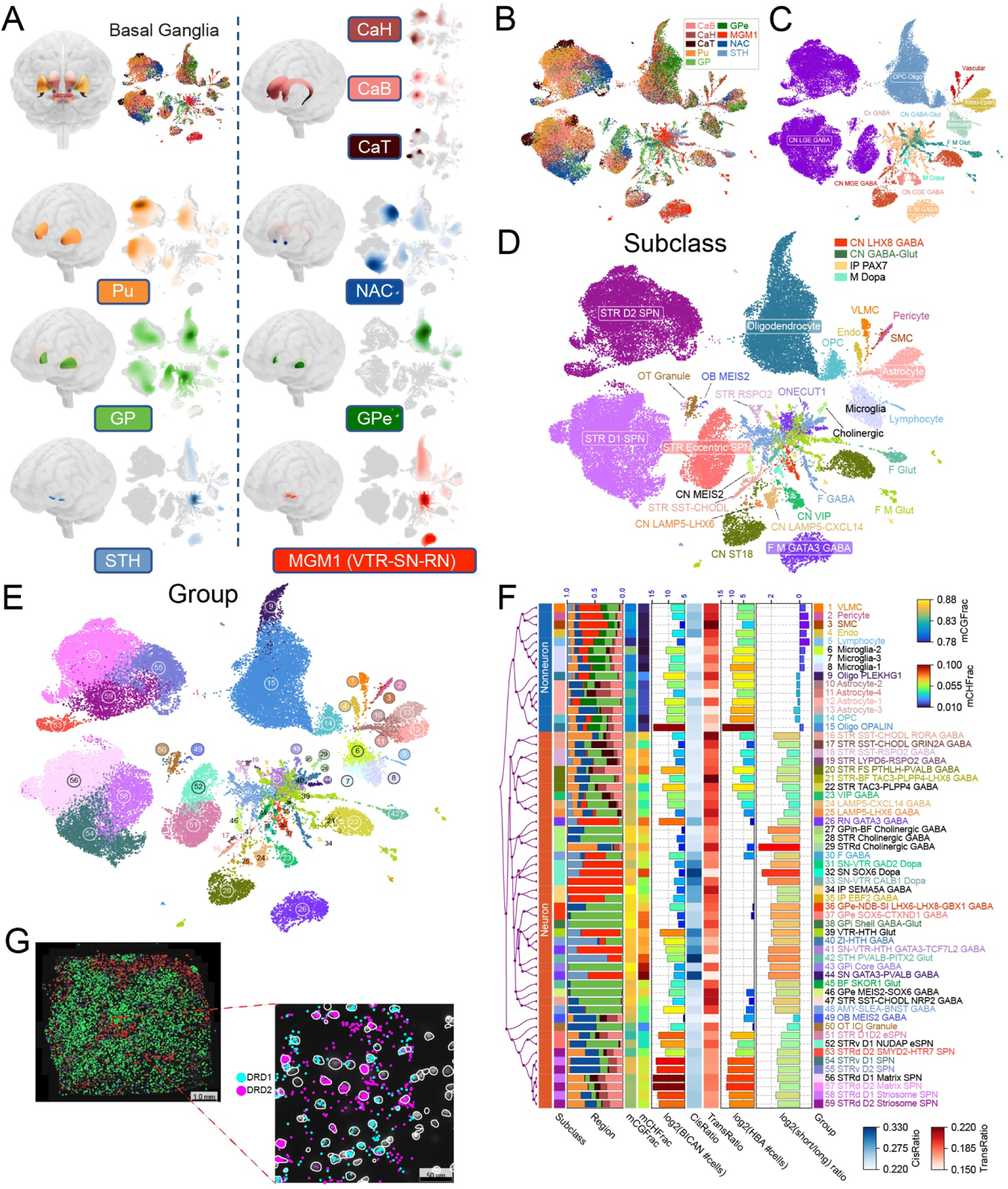
Single-nucleus DNA methylation and 3D genome architecture atlas of the human basal ganglia. (A) Basal ganglia structures and subregions profiled in this study. UMAP embedding is colored by anatomical region, with region-specific cell density distributions shown on the right. UMAP embedding of 261,331 nuclei colored by dissection region (B), class (C), subclass (D), and group (E) level annotation. (F) Epigenomic taxonomy of the human basal ganglia summarizing DNA methylation and Hi-C features. For each group, median values across BICAN cells are shown for mCG, mCH, cis contact ratio (CisRatio), trans contact ratio (TransRatio), and the log2 short-to-long Hi-C contact distance ratio. (G) An example MERFISH sample from the NAC of a single donor. In the zoomed-out image, only SPN cells are shown in color, with different colors indicating different SPN subtypes at the group level. The zoomed-in image shows *DRD1* and *DRD2* gene spots overlaid on the DAPI stain image and the cell boundaries.

To enable robust integration, we corrected for batch effects arising from sample origin, technological platforms, and donor variability (Figure S1B-C), followed by doublet removal (STAR Methods). We then categorized cells into neuron and non-neuron types based on global levels of mCG and mCH methylation (Figure S1C) and the marker-gene-based scoring method (STAR Methods). Neuronal cells were further annotated into Subpallium GABA and Splatter (Figure S1C) based on integration with the Siletti RNA dataset^36^. Subsequently, both neuronal and non-neuronal snm3C-seq cells were clustered and hierarchically aligned to the HMBA BG consensus cell-type taxonomy at the class, subclass, and group levels. These analyses resulted in an epigenomic taxonomy, including 12 classes (Figure 1C), 31 subclasses (Figure 1D) and 59 groups (Figure 1E) based on Seurat integration, regional origin, and the hypomethylation patterns of selected marker genes (See Table S2 and Table S3 for taxonomy and cell type annotations) (STAR Methods). For instance, the highly heterogeneous cluster, splatter, was further resolved into 17 subclasses (Figure S1D) through regional composition (Figure S1E), integration with HMBA RNA dataset (Figure S1F), marker genes-based scoring (Figure S1G), and hypomethylation of key marker genes such as *MEIS2*, *TH*, *LAMP5*, *CXCL14*, *LHX6*, *ONECUT1*, *VIP*, *GATA3*, *PAX7*, *RSPO2*, *SST*, *CHODL* (Figure S1H).

At the subclass level, our annotations showed strong agreement with the HBA^33^ MajorType annotations (Figure S2A), in which STR Eccentric SPN corresponded to Foxp2 and CN ST18 GABA was annotated as Chd7 in HBA, consistent with the gene-body mCH hypomethylation in the respective cell types (Figure S2B). Independent clustering of BICAN cells from this study using 100 kb methylation (mCG + mCH) and 100 kb HiC embeddings yielded consistent cell-type annotations (Figure S2C). Notably, 100kb-resolution HiC embeddings alone were sufficient to distinguish cell classes (Figure S2D) and most subclasses (Figure S2E).

Many subclass- and group-level cell types showed strong specificity to distinct regions of the BG. For example, ventral SPN (STRv D1 and D2 SPN) localized to NAC, while midbrain dopaminergic (M Dopa), F M Glut, and F M GATA3 GABA are found primarily in MGM1 and STH. Similarly, subtypes such as CN MEIS2 GABA, CN ONECUT1 GABA (GPi Core GABA), and CN GABA-Glut (GPi Shell GABA-Glut) were enriched in GP (Figure 1F, hypergeometric p-value < 0.05). The striatum (CaH, CaB, CaT and Pu of the dorsal striatum and NAC of the ventral striatum) was dominated by SPNs (Figure S2F), while STH and MGM1 were enriched for neurons expressing one of the following neurotransmitters: Glutamate, Serotonin, or Dopamine (See Table S4 for regional composition). SPNs captured in GP samples likely originated from the tissue of the laterally adjacent putamen and ventrally adjacent NAC. The GPe, derived from HBA donors (H1930001, H1930002, H1930004), was predominantly composed of non-neuronal cells, particularly oligodendrocytes, consistent with known neuronal scarcity (Figure S2F). Finally, both BICAN and HBA donors exhibited similar cell-type compositions, with most cell types reproducibly identified across donors (Figure S2G), demonstrating the robustness of our cell-type annotations. Among all groups, lymphocyte-like cells displayed the lowest global mCG fraction, while the SN GATA3-PVALB GABA group (a subtype of F M GATA3 GABA) showed the highest mCH fraction.

To compare our methylation-based cell-type taxonomy with transcriptomic classifications, we integrated the annotated snm3C-seq cells with the HMBA^35^ snRNA datasets. Overall, the majority of RNA-defined cell types were also recovered in the methylation data, showing strong cross-modality consistency (Figure S2H-I), except for a few highly heterogeneous types, such as F GABA and F M Glut. Notably, we observed a distinct group, RN GATA3 GABA (group 26 in Figure 1E), that could not be directly aligned with the HMBA BG taxonomy based on the hypomethylation of marker genes. Group 26 primarily originated from MGM1 (Figure 1F), encompassing regions including VTR, SN, and RN, which differs from the SN-VTA localized anatomical definition in the HMBA reference dataset, suggesting that this cell type population comes from the red nucleus (RN), which is supported by our MERFISH data (Figure S3).

The MERFISH dataset was generated from adjacent tissue dissections for snm3C-seq profiling, using a 920-gene panel. The panel was designed using both the HMBA snRNA and the snm3C datasets by identifying marker genes across cell types and brain regions in both the RNA and methylation space (STAR Methods). We sampled the 8 brain regions discussed above across 4 BICAN donors, in replicate (see Figure S3A-D for dissection information). The replicates were imaged from two adjacent slices at separate labs, and showed a strong correlation in transcript counts across all region-donor combinations (Figure S3E). Note that a single region-donor combination (CaT for donor UWA7648) lacked sufficient tissue for a MERFISH experiment. After data processing and QC (Figure S3F), we retained 1,482,528 cells. We annotated the MERFISH data by integrating it with snRNA and methylation data from HMBA and snm3C-seq. 1,159,607 and 1,068,878 cells had high-confidence annotations at the subclass and group level, respectively. A representative annotation for one NAC sample is shown in Figure 1G.

Taken together, we present a comprehensive single-cell atlas of DNA methylation and 3D genome architecture across the human BG, providing a foundational reference for investigating how epigenetic regulation and chromatin organization contribute to brain structure and disease.

### Cell-type-specific DNA methylation landscapes reveal regulatory signatures and disease associations

We first identified DMRs in the CG context across cell types hierarchically from the subclass to the group level. After stringent filtering (ΔmCG ≥ 0.3 and ≥ 2 differentially methylated CpGs per region), we identified 2,361,659 hypo-DMRs and 1,296,880 hyper-DMRs at the subclass level (STAR Methods), including 471,826 and 366,655 cell-type-specific hypo-DMRs that are hypomethylated exclusively in a single subclass or group, respectively (Table S5). The top 500 cell-type-specific hypo-DMRs are shown in Figure 2A. These DMRs display clear cell-type-specific methylation patterns in most populations, except for medium spiny neurons (SPNs), which exhibit high similarity across subtypes. We found that cell-type-specific DMRs tend to contain multiple differentially methylated CpGs, whereas non-cell-type-specific DMRs do not (Figure S4A). Moreover, cell-type-specific hyper-DMRs were significantly enriched in promoter and transcription start site (TSS) regions compared to hypo-DMRs (χ² test, p < 0.05, odds ratio > 2; Figure 2B), indicating that hypermethylation at promoters may contribute more directly to gene silencing and cell-type identity.

**Figure 2.**
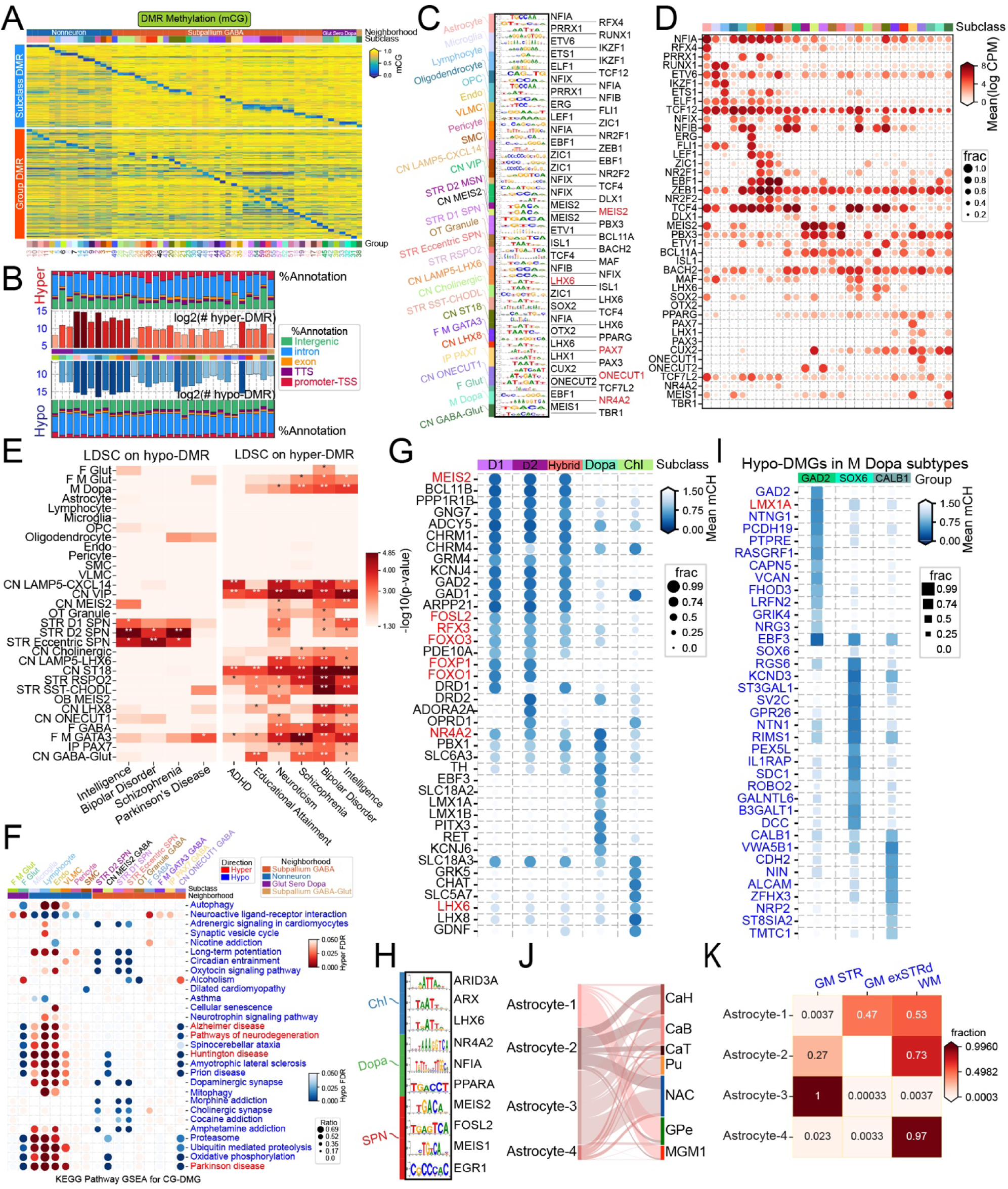
Cell-type-specific DNA methylation patterns reveal disease associations and candidate transcription factor regulators. (A) Top 500 hypomethylated DMRs, ranked by methylation difference (Δ methylation), identified at subclass and group levels. (B) Genomic annotation and counts of cell-type-specific hyper- and hypomethylated DMRs. (C) TF motif enrichment analysis in subclass DMRs, with associated candidate TFs; for visualization, the top three upregulated TFs per subclass are displayed. (D) Dot plot showing upregulated expression of the TFs highlighted in (C) in their corresponding cell types. (E) LD score regression (LDSC) results showing genetic enrichment for subclass-specific hypo- and hyper-DMRs across traits and diseases. (F) Gene set enrichment analysis (GSEA) of subclass-level DMGs against KEGG pathways. Red dots indicate significant enrichment among hypermethylated genes; blue dots indicate enrichment among hypomethylated genes. (G) Dot plot of the top hypo-DMGs distinguishing SPNs, M Dopa, and cholinergic GABAergic neurons. The TFs with binding motifs enriched in the DMRs of the corresponding cell types were highlighted in red. (H) Hypomethylated TFs with motifs enriched on hypo-DMRs in SPNs, M Dopa neurons, and cholinergic GABAergic neurons. (I) Top hypo-DMGs for the three M Dopa subtypes at the group level. The TFs with binding motifs enriched in the DMRs of the corresponding cell types were highlighted in red. (J) Sankey plot of group annotation to region flow for astrocyte subtypes. (K) Confusion matrix between the group level annotation and the reported astrocyte subgroup from Fu et.al 2025.

We next sought to identify TFs and DNA motifs associated with the identified DMRs. Motif enrichment analysis uncovered thousands of enriched motifs across subclasses and groups (STAR Methods). The top upregulated TFs, and their corresponding motifs, are shown in Figure 2C, with each TF exhibiting high expression in its respective cell type (Figure 2D). The results recovered many known cell-type-specific TFs, validating the approach: *MEIS2*, a marker of lateral ganglionic eminence (LGE) origin, exhibits motif enrichment in CN MEIS2 GABA, STR D1, and D2 SPN; *PAX3* and *PAX7* in IP PAX7 GABA; and *ONECUT1* and *ONECUT2* in CN ONECUT1 GABA. At the group level, motifs for *PITX2* were enriched in STH PVALB-PITX2 Glut, *POU6F2* in STR D2 SMYD2-HTR7 SPN (Figure S4B). Several TFs showed motif enrichment within hyper-DMRs as well, including *FOXP1*, *PBX3*, *MEIS2, and POU2F2,* among others (Figure S4C), consistent with previous reports that some TFs can also bind methylated DNA^37^.

To investigate potential links between cell-type-specific methylation and human disease risk, we performed LDSC on hypo- and hyper-DMRs at the subclass level. This analysis revealed that hypo-DMRs in SPNs and hyper-DMRs in many other neuronal types were associated with intelligence, bipolar disorder, and schizophrenia. Additionally, hypo-DMRs in F M GATA3 GABA neurons were associated with PD (Figure 2E). These results highlight the relevance of neuronal methylation landscapes to the risk of neuropsychiatric and neurodegenerative diseases.

We then identified DMGs at the subclass level (STAR Methods and Table S6). Canonical marker genes exhibited the expected gene-body hypomethylation in the relevant methylation contexts: mCH in neurons and mCG in non-neuronal cell types (Figure S4D-F). For instance, glutamatergic neurons (F M Glut and F Glut) showed hypomethylation at *SLC17A6*, while CN GABA-Glut cells, co-expressing glutamatergic and GABAergic neurotransmitters, displayed hypomethylation at both *GAD1*/*GAD2* and *SLC17A6*.

Next, we ranked all genes by the log₂ fold-change in mean normalized methylation fractions between each subclass and all remaining subclasses within the neuronal and non-neuronal categories, followed by Gene Set Enrichment Analysis (GSEA) (STAR Methods). Hypomethylated genes in SPNs were significantly enriched in KEGG pathways related to morphine and cocaine addiction, dopaminergic synapse, long-term potentiation, and circadian entrainment. Interestingly, pathways associated with neurodegenerative diseases were enriched among hypomethylated DMGs (hypo-DMGs) in CN ONECUT1 GABA, F Glut, and CN GABA-Glut, as well as hypermethylated DMGs (hyper-DMGs) in microglia, lymphocytes, and endothelial cells (Figure 2F and S5A-B). Genes contributing to these neurodegeneration pathways, particularly those involved in mitochondrial energy production, proteostasis, and intracellular transport, showed mild hypomethylation in CN ONECUT1 GABA and mild hypermethylation in microglia, relative to randomly selected background cells (Figure S5C).

Furthermore, both GSEA and over-representation analyses revealed that hypo-DMGs in CN ONECUT1 GABA are enriched for synapse organization and axonogenesis (Figure S5D and S5B). Notably, hallmark neurodegeneration-related genes, *MAPT* (tau), *APP* (Aβ), and *APOE* were hypomethylated and highly expressed in CN ONECUT1 GABA cells (Figure S5E), as well as the synaptic vesicle trafficking gene, *SV2A* (Synaptic Vesicle Glycoprotein 2A). Anatomically, CN ONECUT1 GABA neurons originate from the core of the GPi (GPi core), which provides tonic GABAergic inhibition to thalamic and brainstem motor targets. Thus, the hypomethylation and high expression of *SV2A* in these cells are consistent with the high vesicular turnover rate, long projection range, and extensive axonal arborization characteristic of projection neurons in the GPi core, as previously described in non-human primates^38^.

Next, we identified DMGs and DMRs among dopaminergic (Dopa), cholinergic (Chl), and SPNs populations, reflecting their respective roles as dopamine producers, modulators, and receptors. Several TFs showed hypomethylation and elevated expression specifically in SPNs, including *MEIS2*, *FOSL2*, and *FOXO3* (Figure 2G and S5F). In contrast, *NR4A2*, a key TF essential for dopaminergic neuron development, maintenance, and function, was hypomethylated across all three cell types, with the most pronounced hypomethylation observed in Dopa neurons. Cholinergic neurons exhibited greater hypomethylation and higher expression of *LHX6* and *LHX8* (Figure 2G and S5F), which is consistent with the origin of the medial ganglionic eminence (MGE). TF binding motif enrichment analysis of hypo- and hyper-DMRs further corroborated these cell-type-specific regulatory patterns. Enriched TFs included *MEIS2*, *FOSL2*, *RFX3*, *FOXP1*, and *FOXO3* in SPNs; *NR4A2*, *BACH1*/*BACH2* in Dopa; and *LHX6* in Chl (Figure 2H and S5G).

At the group level, we performed clustering and annotation of M Dopa, resolving three distinct subtypes (Figure S5H): SN-VTR GAD2 Dopa (GAD2), SN SOX6 Dopa (SOX6), and SN-VTR CALB1 Dopa (CALB1). Subtype-specific hypo-DMGs were characterized; for example, the GAD2 subtype showed enrichment of *LMX1A*-binding motifs in hypo-DMRs, consistent with hypomethylation and *LMX1A* expression in this population (Figure 2I and S5I). We observed clear regional specificity among astrocyte subtypes (Figure S5J). Astrocyte-2, Astrocyte-3, and Astrocyte-4 are enriched in the striatum (caudate and NAC), whereas Astrocyte-1 is enriched in the GPe and MGM1 (Figure 2J). Cross-referencing with the astrocyte subgroup annotation from the HMBA astrocyte study^39^ (co-submitted), Astrocyte-3 corresponds to the GM STR astrocyte subgroup, while Astrocyte-4 corresponds to the WM astrocyte subtypes (Figure 2K). We also examined microglial methylation heterogeneity at the group level. Microglia clustered into three subgroups (Figure S5K), with microglia-2 appearing more frequently in donors aged 60 or older than in younger donors (Figure S5L). However, the small sample size and confounding by age and sex in our dataset limited the strength of the conclusions. Nevertheless, this pattern aligns with an independent study of aged human hippocampus^40^, accompanied by DNA methylation changes that enhance responsiveness to inflammatory or injury-related signals without sustained transcriptional activation.

### Cell-type-specific compartmentalization and 3D genome architecture in the basal ganglia

Genomes can be partitioned into two distinct domains. Canonically, compartment A is enriched for open chromatin, and compartment B is enriched for closed chromatin^41^. We first identified A/B compartments at the class and subclass levels using HiC data at 100kb resolution, excluding bins with abnormal coverage (assigned as NA). Each bin was thus classified as A, B, or NA. Among all classes, Glut Sero Dopa neurons (including M Dopa and F M Glut) exhibited the weakest compartmentalization, with smaller compartment strength indicating a larger inter-(A-B) to intra-(A-A and B-B) compartment contact ratio. In contrast, non-neuronal populations, particularly OPC-Oligo and immune cells, showed the strongest compartmentalization (Figure S6A-B). Analysis of the BB/AA interaction ratio revealed elevated ratios in immune cells at the class level and in specific neuronal cell types, such as CN ONECUT1 GABA and M Dopa at the subclass level (Figure S6C-D). We examined compartment-specific methylation signatures by plotting mCG methylation saddle plots for each cell type (STAR Methods), which showed compartment-specific methylation in immune cells but less so in other cell types (Figure S6E).

Next, we examined the distribution of chromatin contact distances across different cell types. In general, non-neuronal cell types exhibit longer contact distances than neuronal cell types (Figure S6F), consistent with previous findings^42^. When chromatin interactions were stratified by compartment combinations (A-A, B-B, and A-B and NA-associated), clear differences emerged between neuronal and non-neuronal cell types (Figure S6F). For intra-compartment interactions, neuronal cells primarily exhibited short-range (<5 Mb) contacts. In contrast, non-neuronal cells showed a higher proportion of long-range (≥5 Mb) interactions, with a higher percentage of longer-range contacts in BB than in AA interaction. For comparison, inter-compartment interactions (A-B) were characterized by a substantial enrichment in long-range contacts in both neuronal and non-neuronal populations. This effect was more pronounced in non-neuronal cells, which exhibited predominantly long-range A-B interactions, while neuronal cells contained a mixture of short- and long-range contacts.

To further illustrate these differences, we randomly selected 100 cells per class and annotated interactions on chromosome 10 based on class-specific compartment assignments. Only a small fraction of long-range contacts occurred in neurons, particularly in Glut Sero Dopa (Figure 3A: 12.4% for M Dopa and 14.6% for F M Glut), and only 22.2% of the inter-compartment (A-B) contacts were long-range in M Dopa (Figure 3A). By contrast, non-neuronal cells exhibited substantially more long-range interactions, with 44.4% of long-range contacts in vascular cells at the class level and 51.7% in lymphocytes at the subclass level (Figure S6G). Approximately 50-60% of A-B contacts were long-range interactions (Figure 3A) in non-neuronal populations, such as microglia and lymphocytes at the subclass level (Figure S6G), highlighting the preferential enrichment of long-range, inter-compartment interactions in non-neuronal populations. Additionally, among all non-neuronal cell types, astrocytes exhibit the highest percentage of inter-compartment interaction (39.2%) (Figure S6G).

**Figure 3.**
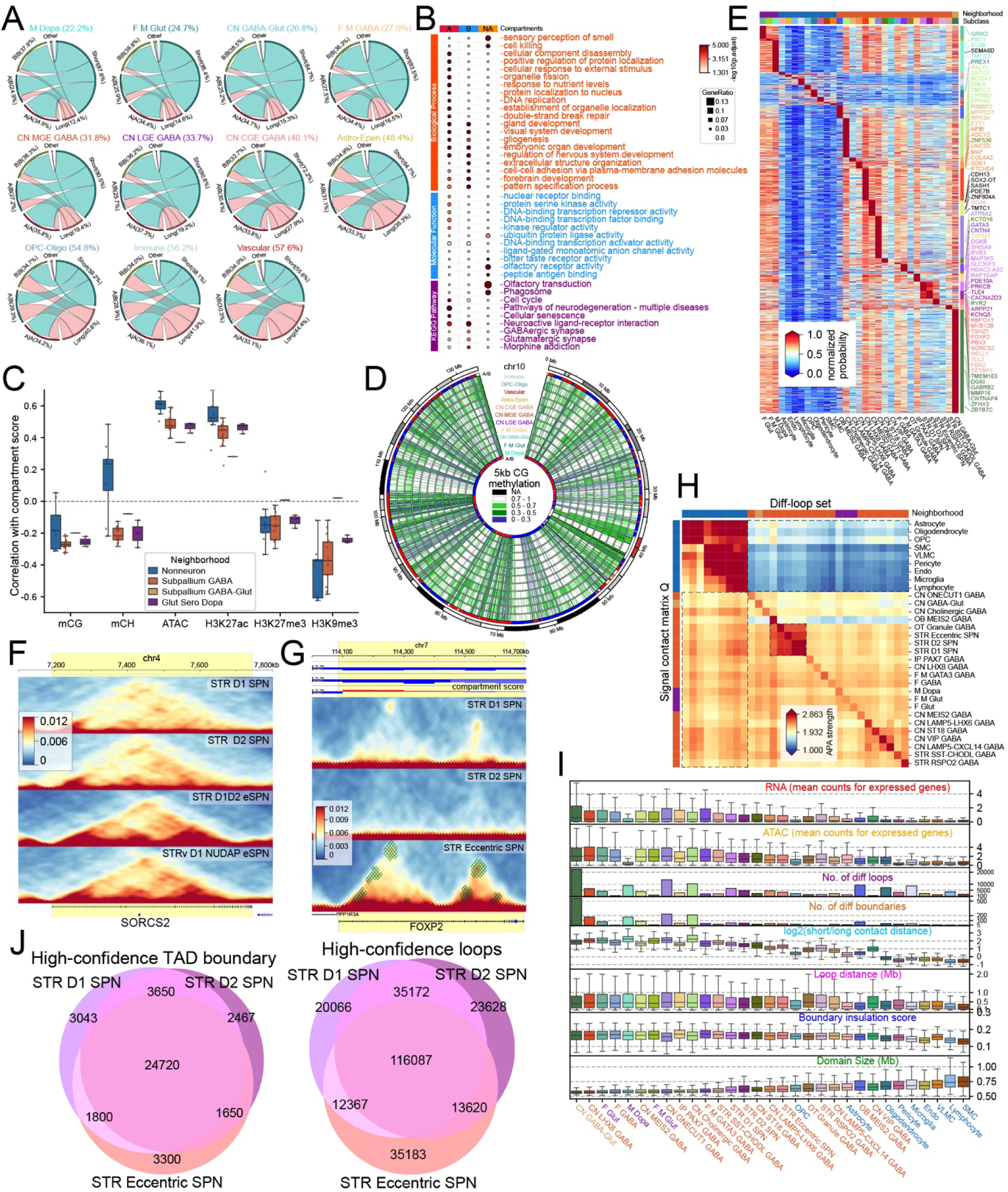
Chromatin interaction dynamics and compartmentalization in the human basal ganglia. (A) Distribution of Hi-C contact distances stratified by A/B compartment status, calculated from 1000 randomly sampled cells per subclass. (B) Gene Ontology (Biological Process, Molecular Function) and KEGG pathway enrichment analysis for genes expressed in A, B, and NA (undefined) chromatin compartments across all cell types. (C) Genome-wide Pearson correlation between subclass-level compartment score and six matched modalities, chromatin accessibility (ATAC), active (H3K27ac) and repressive (H3K27me3, H3K9me3) histone marks, and DNA methylation (mCG, mCH), computed across 100 kb bins. Each point is one subclass, grouped by modality and colored by neighborhood; boxes show median and IQR. ATAC and H3K27ac correlate positively with the A compartment, whereas H3K9me3, H3K27me3, mCG, and mCH correlate negatively, consistent with their association with inactive chromatin. (D) Circos plot depicting mean mCG methylation levels at 5-kb resolution across cell types, with A/B compartment tracks overlaid for immune cells (outer ring) and M Dopa neurons (inner ring). (E) Differential domain boundaries identified across subclass-level cell types, with overlapping genes annotated on the right. (F) Example of a differential domain boundary at the *SORCS2* locus across cell types. (G) Differential domain and chromatin loop structures at the *FOXP2* locus in STR Eccentric SPN, with compartment scores displayed on the top. (H) Cross-Subclass APA reveals lineage-shared and lineage-specific loop programs in human basal ganglia. APA strength for every pair of subclasses: the column subclass’s differential loops are piled up on the row subclass’s imputed Q matrix (21 × 21, 210 kb × 210 kb), and strength is defined as the central pixel divided by the mean of the lower-left 5 × 5 corner. Rows and columns share one hierarchical ordering (average linkage on the symmetrized matrix), so the diagonal reflects self-recovery and off-diagonal blocks reflect cross-Subclass loop sharing. Side bars indicate Neighborhood lineage. Color scale: vmin = 1.0 (no enrichment), vmax = 95th percentile. (I) Cross-cell-type comparison of 3D genome and chromatin features at the subclass level, including domain size, insulation score, loop distance, short/long-range contact ratio, counts of differential domain boundaries and differential loops, and mean ATAC-seq and RNA expression levels. (J) Shared and subclass-specific TAD boundaries and chromatin loops among striatal SPNs. Three-way Venn diagrams of high-confidence TAD-boundary bins (left) and loops (right) across STR D1, D2, and Eccentric SPN subclasses. TAD boundaries were called per 25-kb bin when the per-Subclass boundary probability (fraction of cells calling the bin from single-cell insulation scores) was ≥ 0.05 (top ∼5% genome-wide). Loops were taken from the merged loop catalogue and assigned to a subclass when both z-scored ANOVA F statistics on the imputed Q and local-background-corrected T matrices exceeded 1.036 (top 15% percentile).

To better quantify the number and sizes of A/B compartments, we merged consecutive A or B compartments into superdomains (Figure S6H). Neuronal cell types displayed a lower number of superdomains, which were significantly larger than those in non-neuronal populations. Non-neuronal cells exhibit smaller compartments enriched for long-range contacts, indicating a more finely segmented genome with smaller-scale regulatory control at 100-kb resolution compared to neurons (Figure S6H).

We then investigated the differences between genes expressed in A compartments and those in B compartments. In general, immune cells (microglia and lymphocytes) expressed the most genes in A compartments. Dopaminergic cells (M Dopa), GPi cells (CN ONECUT1 GABA and CN GABA-Glut, also known as the GPi core and GPi shell), and cholinergic GABA cells express the largest number of genes in the B compartment (Figure S6I). The genes expressed in A compartment are enriched for dynamic, regulatory, and housekeeping-like processes that support rapid cellular adaptation (responses to stimuli/nutrients), structural remodeling (component disassembly, organelle dynamics), precise protein/nuclear trafficking and genome stability (replication & repair). Additionally, genes involved in the neurodegenerative disease pathway were enriched in the A compartment. In contrast, genes expressed in the repressive B chromatin compartment showed enrichment for biological processes related to early neural and glial development, including gliogenesis, regulation of nervous system development, embryonic organogenesis, forebrain patterning, and cell adhesion (Figure 3B). This aligns with observations in neuronal systems, where B compartments often maintain repression of developmental and lineage-specification genes, thereby favoring stable neuronal identity in mature cells. The regions with abnormal coverage (annotated as compartment NA) are mainly from repeat regions, including telomeres and centromeres. They are enriched for sensory perception of smell, olfactory transduction, cell killing, and phagosome function, reflecting the evolutionary expansion of olfactory receptor gene clusters within repetitive DNA environments^43–45^. In addition, we reported some genes whose expression was strongly correlated with compartment scores across different cell types (Figure S6J). Most genes are expressed in the A compartment, but several are more likely to be expressed in the B compartment, such as *EBF3* in M Dopa (Figure S6K).

We next examined the interplay between A/B compartment scores and key epigenetic features across cell types. At 100 kb resolution, BG neurons exhibited markedly weaker correlations between compartment scores and both chromatin accessibility and histone modifications than non-neurons (Figure 3C). Specifically, the genome-wide Pearson correlation with the active mark H3K27ac was r = 0.53 in non-neuronal populations, compared to r = 0.42 in GABAergic neurons. Likewise, the repressive heterochromatin mark H3K9me3 showed a mean r of −0.44 in non-neuronal cells versus r = −0.36 in neurons.

DNA methylation analysis revealed a robust association between compartment identity and methylation status. Regions with low or intermediate methylation (5 kb mCG < 0.7) were significantly overrepresented in A compartments, with the most pronounced enrichment in non-neuronal lineages (Figure 3D). In oligodendrocyte precursor cells and mature oligodendrocytes (OPC-Oligo), hypomethylated loci were enriched 3.4-fold in A compartments relative to expectation (χ² test, odds ratio = 3.4, P = 5.08 × 10⁻³²).

Next, we called domains (25 kb) and loops (10 kb), and performed differential analysis at subclass and group levels. We identified 2,110 differential domain boundaries (Figure 3E) and 147,791 differential loops (Figure S6L) across all subclasses. 75 out of 2110 (3.55%) of these boundaries overlapped with genes that show both hypomethylation and upregulation in the corresponding cell type. This represents a 23.6-fold enrichment over the random expectation (permutation test P < 0.0001, STAR Methods), indicating that 3D domain reorganization is coupled with cell-type-specific transcriptional and epigenetic programs. For example, the gene *SORCS2* is hypomethylated and upregulated in STR Eccentric SPN, more specifically, in group STR D1D2 eSPN. It exhibits distinct domain boundaries and a higher frequency of interactions within the gene body (Figure 3F). This is consistent with its reported higher expression^11^ and hypomethylation (Figure S6M) in STR D1D2 eSPNs. Previous work showed that *SORCS2* is reduced and mislocalized in Huntington’s disease due to selective interaction with mutant huntingtin, thereby impairing *NR2A* receptor trafficking in striatal SPNs and contributing to synaptic dysfunction and motor deficits^46^.

We further examined the relationship between differential loops, chromatin accessibility, and DNA methylation. The anchors of differential loops were enriched for regions of high chromatin accessibility and low DNA methylation (Figure S6L). Notably, loop strength, ATAC signal, mCG score, and mCH score at STR D1 SPN- and STR D2 SPN-specific anchors showed the greatest similarity to each other amongst all cell-types, suggesting a very similar epigenetic state in these two cell-types. Additionally, differential looping at key marker genes such as *FOXP2* (Figure 3G) and *RELN* (Figure S6N), which define the STR Eccentric SPN and STR D1 SPN subclasses, respectively, exhibited cell-type-specific looping patterns and increased interaction frequencies within their gene bodies in the corresponding subclasses. *FOXP2* resides in the A compartment in eSPNs, whereas it is located in the B compartment in all other cell types examined.

Next, aggregate peak analysis (APA) was used to detect whether differential loops (diff-loops) in any given subclass also exist as focal contacts in other cell types. Assigning a single strength score to each APA window (central pixel/corner background) showed that diagonal entries (mean = 2.45) far exceeded off-diagonal entries (mean = 1.83; one-sided t-test P < 1 × 10⁻³⁰), confirming that diff-loop calls are subclass-specific rather than driven by global compaction differences. Hierarchical clustering recovered the expected lineage organization: non-neuronal cell types formed one block, SPN subclasses another, and the remaining GABAergic/glutamatergic/dopaminergic subclasses clustered by class (Figure 3H), supporting that 3D loop architecture, like methylation and chromatin accessibility, encodes lineage identity.

The matrix is conspicuously asymmetric, however. The upper-right block (non-neuron signal × neuronal diff-loops) is depleted, whereas the lower-left block (neuron signal × non-neuron diff-loops) is nearly as strong as the neuronal diagonal. Neuron-specific loops collapse from neurons to nonneurons with ∼40% loss in interaction frequency, while nonneuron-specific loops are essentially unchanged between lineages (<11%). Hence the cell-type-specific 3D contact landscape of the human BG is driven mainly by neuronal loop gain: neurons acquire many lineage-restricted loops that disappear in glia, while most nonneuron-specific loops are actually constitutive contacts that are flagged as glial only because their strength varies most among glial subclasses.

We observed that non-neuronal cells exhibit larger average chromatin domain sizes but lower insulation scores (Figure 3I), indicating stronger insulation^47^. This suggests that non-neuronal genomes are partitioned into fewer, larger, and more strongly insulated domains, rather than the pattern observed at the compartment level. In contrast, neuronal cells exhibit smaller domains with weaker insulation, consistent with their higher proportion of short-range contacts and the previously reported elevated TAD-scale contact density in neurons^48^. Although non-neuronal cells display more long-range contacts, these long-range interactions are unlikely to be dominated by focal looping, as the characteristic loop distances are shorter in non-neuronal cells (loop distance in Figure 3I).

Neurons exhibited more differential domain boundaries, more differential loops, increased chromatin accessibility (ATAC counts), and higher average gene expression per cell compared with non-neuronal cells (Figure 3I). Together, these results indicate a more stable, less dynamically reconfigured 3D genome architecture in non-neuronal cells, consistent with lower regulatory activity compared with neuronal cells.

To link chromatin architecture with genetic risk, we then applied linkage disequilibrium score regression (LDSC) using loop anchors as functional annotations. This analysis showed that loop anchors from different cell types were significantly enriched for GWAS heritability of brain-related traits. For example, intelligence, schizophrenia, and bipolar disorder signals were most strongly associated with loop anchors from GABA neurons in LGE, CGE, and MGE. In contrast, PD enrichment was observed in STR SST-CHODL GRIN2A GABA loops (Figure S6O), which is consistent with the association of PD and hypo-DMR of STR SST-CHODL (Figure 2E), and multiple sclerosis enrichment was detected in lymphocyte loop anchors (Figure S6O).

Finally, we assessed whether the three striatal SPN subclasses are distinguishable at the 3D genome level by intersecting their high-confidence loop and TAD-boundary catalogues (Figure 3J). At the TAD-boundary level, the picture is markedly more conserved. Of ∼32,000 high-confidence boundaries per subclass, ∼75% are shared by all three, and private boundaries are an order of magnitude smaller (D1 = 3,043; D2 = 2,467; Eccentric = 3,300). At the loop level, a pan-SPN core of 116,087 loops is shared by all three subclasses, likely reflecting striatum-identity regulation. An additional 35,172 loops are co-activated in D1 and D2 SPN but are lost in the eccentric SPN, consistent with a canonical-SPN program partially remodeled in the eccentric population. Subclass-private fractions are asymmetric, with eccentric SPN carrying the most private loops. Thus chromatin loops, not TAD boundaries, carry the cell-type-resolving signal distinguishing D1, D2 and eccentric SPNs: the TAD scaffold is mostly striatum-shared (∼75% overlap), whereas a mass of focal loops are gained or lost between subclasses, with eccentric SPN the most reorganized of the three.

### DNA methylation defines SPN subtypes and regional signatures

D1- and D2-SPNs were further annotated at the group level into matrix (D1M, D2M), striosome (D1S, D2S), and Ventral (D1V, D2V) SPN subtypes (Figure 4A). This annotation was supported by integrative analysis, mCH hypomethylation within ±2 kb of marker gene bodies (Figure S7A), and the dissection region (Figure 4B). Ventral SPNs mainly originated from the NAC, whereas dorsal SPNs were derived from the caudate and putamen (Figure 4C). The Eccentric SPN population was further annotated into STRv D1 NUDAP eSPN (NUDAP) and STR D1D2 eSPN (D1D2). Consistent with their mixed identity, D1D2 eSPNs showed gene-body hypomethylation at both DRD1 and DRD2, with the DRD2 hypomethylation being weaker, paralleling the modest DRD2 expression observed in this population. We also validated a previously unrecognized subtype within STR D2 SPNs, termed STR D2 SMYD2-HTR7 SPN (D2 SMYD2), which was independently defined by the BICAN/HMBA basal ganglia consensus taxonomy^35^, and also supported in companion studies, including cross-species MERFISH spatial transcriptomic data^49^, macaque Patch-seq electrophysiology data^50^ and Droplet Paired-Tag RNA and histone modifications datasets^51^. D2 SMYD2 cells showed hypomethylation at DRD2 and clustered closely with D2 SPNs in methylation-based uniform manifold approximation and projection (UMAP) embeddings.

**Figure 4.**
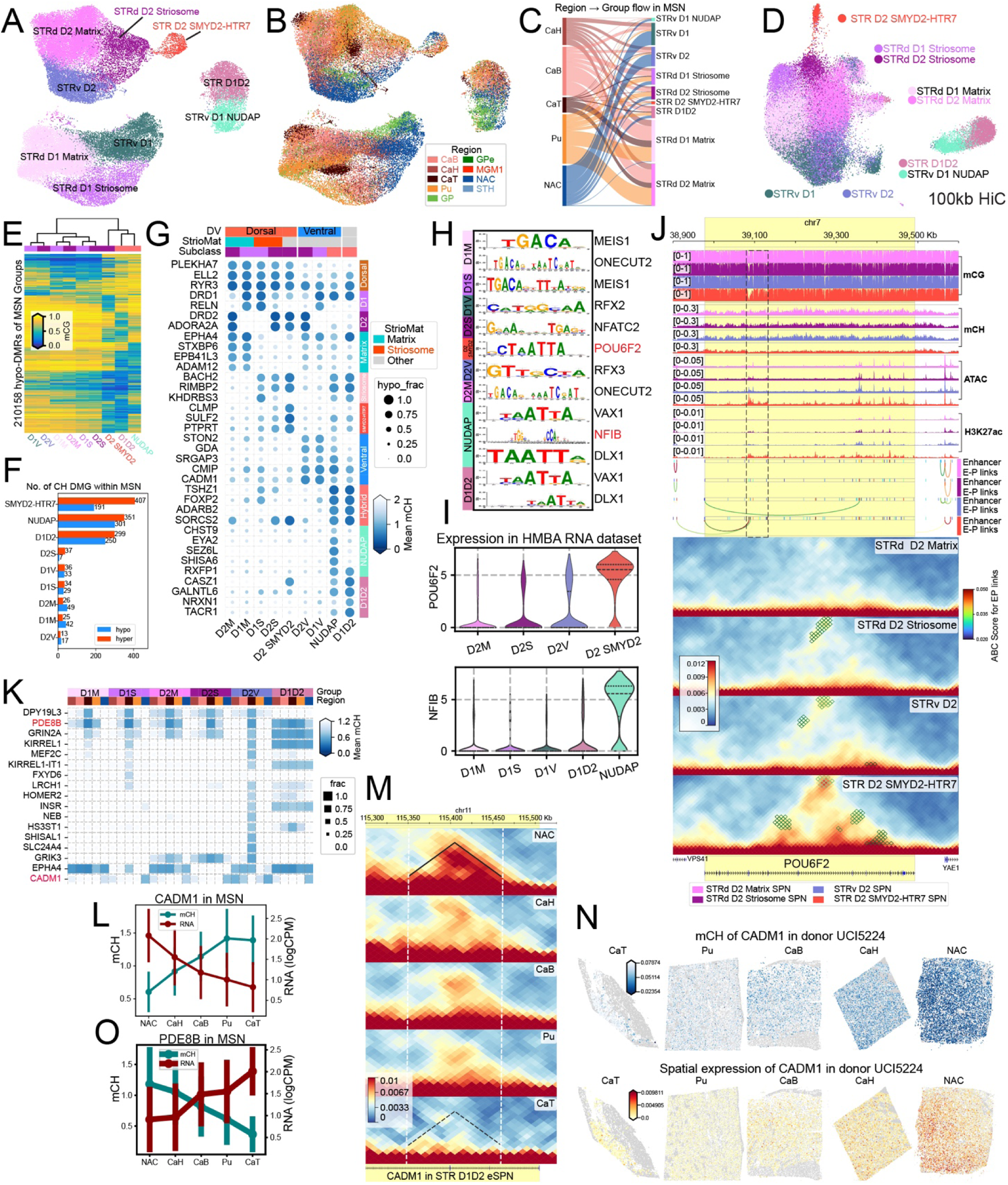
DNA methylation patterns define SPN subtypes and regional signatures in the striatum. (A) UMAP visualization of SPNs based on 100-kb binned DNA methylation features (mCG and mCH), colored by group-level SPN annotations (matrix, striosome, ventral, and eccentric subtypes). (B) The same methylation-based UMAP, colored by anatomical region of origin. (C) Sankey plot of dissection region to group level annotation for SPN subtypes. (D) UMAP embedding of SPNs using 100-kb resolution 3D genome contact profiles (Hi-C-derived features), colored by SPN group annotations. (E) Hierarchical clustering of group-specific hypomethylated differentially methylated regions (hypo-DMRs) within SPNs (one group versus all other groups), based on CG methylation levels. (F) Bar plot showing the number of differentially methylated genes (DMGs) identified for each SPN group (one group versus all others), stratified into hypermethylated and hypomethylated categories. (G) Gene-body methylation levels of canonical marker genes and the top hypo-DMGs (rank by Wilcoxon statistic) defining each SPN group, confirming subtype-specific annotations through expected patterns of marker hypomethylation. (H) Enriched TF motifs and corresponding hypomethylated TFs identified in hypo-DMRs of each SPN group (one group versus all remaining groups). (I) High expression of transcription factors POU6F2 and NFIB in D2 SMYD2 and NUDAP subtypes respectively. (J) Genomic view of *POU6F2* locus showing concordant D2 SMYD2 specific activation across all five modalities (CG/CH hypomethylation, elevated chromatin accessibility, increased Hi-C contact frequency, strong H3K27ac, and reduced H3K27me3/H3K9me3), representative of a subtype-enriched transcription factor. The black box highlights the hypomethylated and highly accessible region, representing a candidate enhancer of *POU6F2*. (K) Dot plot illustrating regional hypomethylation of DMGs across SPN groups; dot size represents the fraction of cells showing hypomethylation (normalized mCH < 1) in the corresponding anatomical region. (L) Gradient of gene expression and CH methylation levels for *CADM1*, progressing from anterior NAC to posterior CaT in the basal ganglia. (M) Chromatin interaction frequencies at the *CADM1* locus, demonstrating a gradient of change from anterior (A) to posterior (P) region of the basal ganglia in striatal D1D2 eSPN. (N) Spatial validation of the regional DMG *CADM1* across BG regions in all SPN cell types, integrating independent spatial methylation and gene expression datasets. (O) Gradient of gene expression and CH methylation levels for *PDE8B*, progressing from posterior CaT to anterior NAC in the basal ganglia.

3D genome embedding using 100-kb resolution Hi-C contact profiles across all SPN subtypes revealed weaker separation between D1 and D2 SPNs (Figure 4D and S7B) compared to RNA^51^ and methylation embeddings. However, it effectively distinguished dorsal, ventral, matrix, striosome and eccentric populations (Figure 4D and Figure S7C). Notably, clustering based on hypo-DMRs grouped SPN cells into matrix, striosome, and ventral lineages, with D2 SMYD2 positioned adjacent to eccentric SPNs (Figure 4E). This positioning is consistent with macaque Patch-seq characterization of this subclass^50^, which reported intermediate morphoelectric properties between STR D1D2 eSPN and STR D2 SPN, suggesting concordance between epigenetic and functional positioning along the SPN landscape. Differential methylation analysis identified the greatest number of hyper-DMGs in D2 SMYD2, followed by NUDAP and D1D2, whereas NUDAP exhibited the largest number of hypo-DMGs, followed by D1D2 and D2 SMYD2 (Figure 4F). Hypomethylation of established marker genes and the top hypo-DMGs for each subtype was reported (Figure 4G). Key subtype-specific hypo-DMGs included *DRD1* and *RELN* (D1 SPN), *DRD2* and *ADORA2A* (D2 SPN), *TSHZ1* and *FOXP2* (eccentric SPN), *EPHA4* and *STXBP6* (matrix), *BACH2* (striosome), *SRGAP3*, *CMIP* and *GDA* (Ventral), *EYA2* and *RXFP1* (NUDAP), and *CASZ1* (D1D2), consistent with their high expression in published RNA datasets^11,35^. Several subtype-specific hypo-DMGs have also been implicated in neurological and psychiatric disorders. In D2 SMYD2, disease-associated genes included *CLMP* (neuroinflammation^52^), *SULF2* (AD^53^), and *SMYD2* (stroke, cardiovascular disease^54^, cancer^55^). Consistent with the methylation signatures, many of these genes also showed subtype-specific differences in chromatin accessibility, histone modifications, and chromatin interactions. For example, in D2 SMYD2, *SMYD2* and *SULF2* displayed CG and CH hypomethylation, elevated chromatin accessibility, increased Hi-C interaction frequency, and strong H3K27ac, coupled with reduced H3K27me3 and H3K9me3, a combination indicative of an active regulatory state (Figure S7D). Enrichment analysis revealed that hypo-DMGs in NUDAP and D2 SMYD2 were associated with synapse organization (Figure S7E).

TF motif enrichment analysis of hypo-DMRs across SPN groups identified candidate regulators with corresponding hypomethylation patterns, including *ONECUT2* in matrix SPNs (D1M and D2M), POU6F2 in D2 SMYD2, NFIB in NUDAP (Figure 4H), *BACH2* in striosome SPNs, *RFX3* in ventral SPNs (Figure S7F), and *PBX1* on hyper-DMRs in eccentric SPNs (eSPN) (Figure S7F). Several enriched TFs, including *POU6F2* and *NFIB*, are selectively expressed in their corresponding groups (Figure 4I). The *POU6F2* motif was enriched in D2 SMYD2 hypo-DMRs in both the one-versus-rest (Figure S4B) and one-versus-other-SPN (Figure 4H) comparisons, highlighting the robustness of the *POU6F2* signal in D2 SMYD2 group. Consistent with a regulatory role, the *POU6F2* locus itself displays group-specific hypomethylation, elevated chromatin accessibility and H3K27ac, higher contact frequency within TAD, and more enhancer-promoter links (ABC model; STAR Methods) in D2 SMYD2 (Figure 4J). Interestingly, *POU6F2* has also been described as a marker of a hindbrain-derived SNr subclass that co-expresses *POU6F2*, ZFPM2 and *PAX5*^56^. Our D2 SMYD2 group similarly co-expresses *POU6F2* and *ZFPM2* but lacks PAX5 (Figure S7G), a classical r1 hindbrain lineage marker, consistent with its LGE-derived telencephalic origin.

Regional differential methylation analysis revealed several genes showing methylation differences across striatal subregions, stratified by SPN group (Figure 4K). Notably, *CADM1* displayed pronounced hypomethylation in the NAC compared with other regions in all SPN cell types, consistent with elevated *CADM1* expression in the NAC (Figure 4L and S7H). *CADM1* also exhibited a gradient of 3D chromatin interaction frequency from anterior NAC, CaH to CaB and posterior CaT in STR D1D2 eSPN (Figure 4M), corroborated by corresponding gradients in gene expression and CH methylation (Figure 4L). Spatial integration of methylation profiles with expression data confirmed a concordant gradient of hypomethylation and elevated *CADM1* expression in the NAC (STAR Methods, Figure 4N). In contrast, *PDE8B*, a gene implicated in Autosomal-Dominant Striatal Degeneration (ADSD)^57,58^ showed the opposite gradient of hypomethylation and high expression from CaT to hypermethylation and reduced expression in NAC (Figure 4O and S7G), also supported by our spatial data (Figure S7I). Analysis of 3D genome organization further revealed that NAC exhibited the smallest domain sizes and strongest insulation scores, while CaT showed larger domains and weaker insulation in SPNs (Figure S7J), suggesting elevated transcriptional activity in the NAC.

### Cross-modal comparison reveals coordinated epigenetic and transcriptional regulation across basal ganglia cell types

In addition to DNA methylation and Hi-C data, we incorporated RNA-seq, ATAC-seq, and histone modification datasets generated by other consortium groups and us. All multi-omic profiles were harmonized to the shared HMBA BG consensus cell-type taxonomy, enabling fully integrated cross-modality analyses. We performed a systematic cross-modality comparison across DNA methylation, transcription, chromatin accessibility, and histone modifications. First, we examined the relationship between gene-body methylation (mCG and mCH) and gene expression by stratifying genes in each cell type into four categories: DMG-only, DEG-only, overlapping DMG&DEG, and background genes (no differential signal in either modality). Consistent with gene-body methylation serving as an inverse proxy for transcriptional activity, gene-body methylation was significantly reduced in DMG-only and DMG&DEG genes (predominantly mCG in non-neuronal cells and mCH in neuronal cells), whereas this pattern was not evident for DEG-only genes (Figure S8A). Notably, DMG-only and DMG&DEG genes also showed reduced methylation across the gene body and promoter-proximal regions spanning TSS −1500 to Transcription End Site (TES).

Next, we assessed methylation patterns over regulatory elements defined by ATAC-seq and histone modification peaks. As expected, both mCG and mCH were depleted over ATAC and H3K27ac peaks relative to flanking regions. Interestingly, mCH exhibited a non-monotonic profile at repressive marks: H3K27me3 and H3K9me3 peaks showed overall lower mCH than surrounding genomic regions, yet within the peak itself, mCH reached a local maximum at the peak center and decreased toward the boundaries (Figure S8B); the basis for this pattern remains unclear. We also quantified ATAC and histone modification signals over cell-type-specific CG hypo- and hyper-DMRs. ATAC and H3K27ac signals were consistently elevated at hypo-DMRs across cell types, whereas signal patterns at hyper-DMRs were more variable, suggesting heterogeneous regulatory roles for hypermethylated regions (Figure S8C). H3K9me3 generally showed an inverse relationship with DNA methylation, with higher signal at hypo-DMRs and lower signal at hyper-DMRs, in SPNs and non-neuronal cell types. In contrast, H3K27me3 was broadly enriched over both hypo- and hyper-DMRs across cell types (Figure S8C). We assessed the compartmental distribution of domain boundaries, chromatin accessibility, histone modifications, and DMRs. Domain boundaries were generally depleted from both A and B compartments, whereas ATAC-seq peaks, H3K27ac, and DMRs (hyper and hypo-DMRs) were enriched in A compartments and depleted in B compartments. In contrast, the repressive marks H3K27me3 and H3K9me3 showed preferential enrichment in B compartments (Figure S8D).

### Integration of multi-omic data reveals cell-type-specific gene regulatory networks

To construct cell-type-specific gene regulatory networks, we integrated multiple genomic modalities beginning with hypo-DMRs linked to TFs through transcription factor binding site (TFBS) motif enrichment analysis (Figure 5A). We applied the Activity-by-Contact (ABC) model to infer enhancer-promoter (E-P) interactions using chromatin accessibility (ATAC-seq), Hi-C contact frequency, and RNA expression as input. E-P links exceeding the ABC score threshold of 0.021 (Figure 5B) were retained, with the requirement that target genes exhibit hypomethylation, transcriptional upregulation, or elevated chromatin accessibility in the corresponding cell type (See Table S7 for predicted enhancer-promoter links). This integrative approach identified 108,203 unique E-P linkages across cell types at the subclass level, of which 9,158 were associated with target genes that showed concurrent hypomethylation, upregulation, and high accessibility (Figure S8E). Notably, upregulated target genes were associated with a median of four distal enhancers, significantly exceeding the connectivity observed for non-upregulated genes (p-value=4.22e-31).

**Figure 5.**
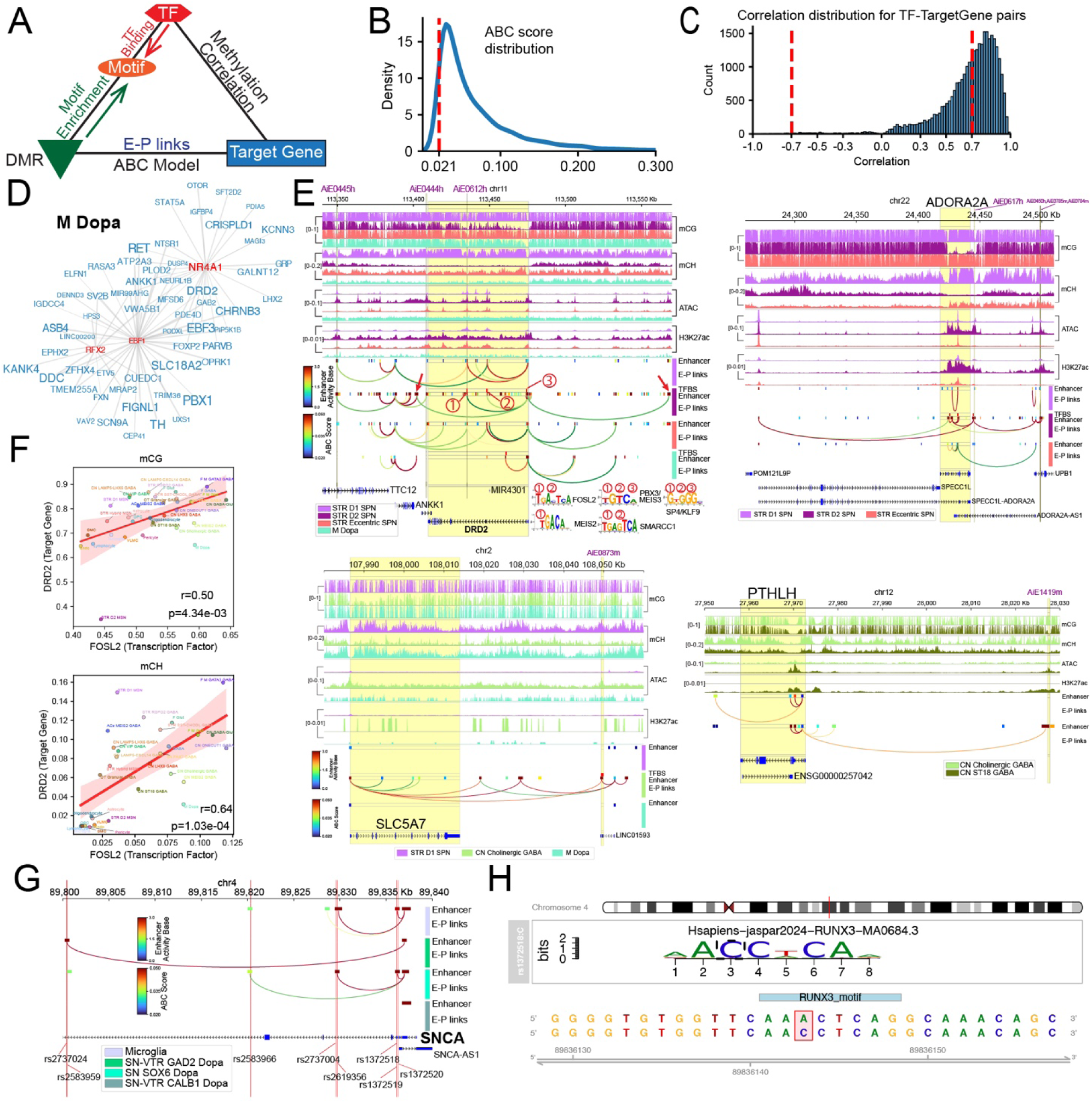
Multi-omic integration reveals cell-type-specific gene regulatory networks. (A) Schematic representation of the integrative analytical framework for identifying candidate cell-type-specific TF-target gene regulatory interactions. The workflow incorporates DMRs, TFBS motif enrichment analysis, activity-by-contact (ABC) model-predicted enhancer-promoter links, and TF-target gene methylation correlation coefficients. (B) Distribution of ABC scores across all predicted enhancer-promoter links. Significant links were defined using the ABC model recommended threshold of 0.021. (C) Distribution of Pearson correlation coefficients for gene body methylation levels between TF and target genes across cell types. TF-target gene pairs exhibiting strong correlation (|r| ≥ 0.7) were retained for downstream analysis. (D) Predicted gene regulatory network architecture in M Dopa. Red nodes represent transcription factors; blue nodes represent target genes. Edge width reflects the magnitude of TF-target methylation correlation. Node size indicates the significance of gene hypomethylation (larger nodes denote greater statistical significance). (E) Genomic view of the *DRD2, ADORA2A, SLC5A7*, and *PTHLH* illustrating ABC-predicted enhancer-promoter interactions in the corresponding cell types. Purple annotations highlight enhancers curated by the enhancer-AAV toolbox. Circled numbers (1-3) indicate hypomethylated regions (hypo-DMRs) enriched for TFBS with distinct motifs, and the red arrow indicates new candidate enhancers in STR D2 SPN. (F) Representative TF-target gene pair (*FOSL2*-*DRD2*) demonstrating correlated methylation patterns. Pearson correlation between *FOSL2* and *DRD2* calculated from pseudobulk CG and CH methylation profiles. (G) Cell type-specific enhancer landscape at the *SNCA* locus across microglia and M Dopa subtypes at the group level. Vertical lines mark six genome-wide association study (GWAS) single-nucleotide polymorphisms (SNPs) associated with Parkinson’s disease risk. (H) rs1372518 disrupts the RUNX3 binding motif. rs1372518 (chr4:89,836,140; A>C) is located within a RUNX3 binding motif (JASPAR ID MA0684.3). Top: RUNX3 position weight matrix (PWM) logo. Bottom: genomic sequence flanking the SNP, with reference (A, top) and alternative (C, bottom) alleles shown; the SNP position is highlighted in red. The C risk allele increases the predicted motif match score for RUNX3 by an alleleEffectSize of 0.158 (ratio of allele difference over maximal allele score) (strong effect, allele difference ≥ 0.7).

To further refine candidate TF-target gene pairs, we calculated Pearson correlation coefficients using pseudobulk methylation profiles between TFs and their putative target genes across all cell types. TF-target pairs were retained only if they exhibited strong methylation correlation (|r| ≥ 0.7; Figure 5C) and the target gene demonstrated hypomethylation (See Table S8 for predicted TF-Target gene pairs). This framework yielded cell-type-specific TF-target gene regulatory networks, in which a predicted regulatory relationship indicates that the TF’s cognate binding motif resides within a hypo-DMR-containing enhancer that likely regulates the target gene’s transcriptional output. For instance, *NR4A1*, previously identified as a hypomethylated and upregulated TF, was predicted to bind and potentially regulate multiple target genes in M Dopa (Figure 5D).

To benchmark our predicted cell-type-specific enhancer-promoter interactions against orthogonal experimental evidence, we compared our predictions with striatal enhancers curated by the enhancer-AAV toolbox^59^. These enhancers were identified from human and mouse ATAC-seq datasets as differentially accessible chromatin regions near striatal cell-type marker genes and were functionally validated in vivo in mouse, with select SPN enhancers additionally tested for conserved in vivo activity in rat and macaque. We successfully recapitulated some enhancers in our predictions, including AiE0445h, AiE0444h, AiE0612h at *DRD2*, and AiE0617h at *ADORA2A* in D2 SPN, AiE0873m at *SLC5A7* in CN Cholinergic GABA, AiE1419m at *PTHLH* in CN ST18 GABA (Figure 5E), AiE0603m at *POU3F1* locus in pan-SPN and AiE0779m at *SLC35D3* in D1 SPN (Figure S8F). This recapitulation represents an orthogonal computational recovery of mouse-validated enhancer-promoter linkages in their corresponding human cell types, rather than direct functional validation of these enhancers in human tissue. All of the ABC model predicted cell-type-specific enhancer-promoter links can be visualized at https://neomorph.salk.edu/SCMDAP/BasalGanglia. A further illustrative example is GATA3 in F M GATA3 GABA, where the gene displayed hypomethylation, high chromatin accessibility, elevated H3K27ac signal, robust expression, enriched E-P interactions, and strengthened chromatin contact frequencies (Figure S8G). Additionally, candidate cell-type-specific enhancers not previously reported in the corresponding cell types were identified. We also identified some hypomethylated DMRs overlapping with predicted enhancers that were enriched for TF binding motifs, including *MEIS1*, *MEIS2*, *MEIS3* and *KLF9*/*SP4* at the *POU3F1* locus, *FOSL2*, *MEIS2*, and *PBX3*/*MEIS3* at *DRD2* locus (Figure 5E). These TFs were predicted to regulate target gene expression by binding to their corresponding enhancers.

As a representative example, *FOSL2* was predicted to putatively regulate *DRD2* expression, with *FOSL2* binding sites at the *DRD2* locus supported by independent TFBS resources, including TFLink^60^ and GeneHancer^61^. Gene body methylation of *FOSL2* was strongly correlated with *DRD2* across cell types in both CG and CH contexts (Figure 5F), with *FOSL2* maintaining stable hypomethylation in D2 SPNs despite exhibiting sporadic expression patterns (Figure S8H). This observation suggests that FOSL2 expression may be transient and subject to rapid mRNA turnover, while stable DNA hypomethylation at the FOSL2 locus could serve as a persistent epigenetic signature marking the cell’s competence to express this TF.

We next examined enhancers at the *SNCA* (Synuclein Alpha) locus, a gene strongly associated with PD^62–64^, across microglia and three midbrain dopaminergic (DA) neuron subtypes (SN-VTR GAD2 Dopa, SN SOX6 Dopa, and SN-VTR CALB1 Dopa). PD-associated GWAS variants were found to localize to *SNCA* enhancer regions in microglia and in specific midbrain DA subtypes (Figure 5G). Two *SNCA* enhancer variants (rs2737024 and rs2583959), previously associated with elevated PD risk^65^, were found to reside within enhancers active specifically in the SN-VTR GAD2 Dopa subtype. Two additional variants forming a risk haplotype (rs2737004 and rs2619356) lie within the SNCA enhancer that is accessible in both microglia and SN SOX6 Dopa neurons (Figure 5G). This enhancer has been functionally characterized in microglia by Booms et al.^66^, where the risk alleles alter *CTCF/RAD21* and *CEBPB* binding and the enhancer controls *SNCA* and *MMRN1* expression via cell-type-specific enhancer-promoter looping. Our snm3C-seq data shows that the same enhancer is also active in SN SOX6 Dopa neurons, a midbrain DA population particularly vulnerable in PD^67^, raising the possibility that this regulatory module operates in both immune and dopaminergic populations relevant to disease. To extend this analysis, we applied motifbreakR^68^ to PD-associated SNPs within cell-type-specific SNCA enhancers and identified an additional candidate variant, rs1372518, which was reported to be associated with Lewy body dementia in PD^69^, significantly increasing the predicted motif match for RUNX3 (Figure 5H), with the C risk allele creating a stronger RUNX3 binding site than the A reference allele. *RUNX3* is hypermethylated and lowly expressed at baseline in both microglia and SN SOX6 Dopa, but RUNX3 protein is significantly upregulated in MPTP-induced PD models, where it transcriptionally activates downstream targets that drive dopaminergic neuron apoptosis and disrupt dopamine homeostasis^70^. These observations suggest that rs1372518 creates a latent RUNX3 binding site whose function may emerge upon disease-associated upregulation of *RUNX3*. Given the cell-type-specific enhancer-promoter contacts linking this region to *SNCA*, this mechanism may contribute to PD-associated dysregulation of *SNCA* in dopaminergic neurons, although direct functional validation will be needed. In addition, the protective variants rs1372519 and rs1372520^71^, were also identified within cell-type-specific regulatory elements at this locus.

### Integration with spatial transcriptomic datasets highlights gene expression and methylation gradients

Custom preprocessing, including cell resegmentation and QC (STAR Methods) was performed on the MERFISH dataset (Figure S3F, Table S9). Cells were annotated by integrating each sample with a region-matched HMBA reference dataset (Figure S9A and STAR Methods). Good overlap was observed at the subclass level between the MERFISH and reference data across samples; however, specific group-level cell types exhibited variable annotation scores (Figure S9B-C). Expected distributions of neuronal and non-neuronal cells were observed across brain regions (Figure S9D-F). Neuronal cells from individual samples were integrated with region-matched subsets of the snm3C dataset, as previously described^32^, using the methylation-specific marker genes included in the gene panel (Figure S9G-H; STAR Method). There was good agreement between the HMBA and snm3C annotations at the subclass level, with modest agreement at the group level (Figure S9I-J). Visual inspection of the data revealed differential levels of White Matter (WM) across samples, so a method to identify WM regions was developed (STAR Methods), noting that WM regions exhibit different global QC metrics than non-WM regions (Table S10). To address this issue, all annotated WM regions were removed from the relevant downstream analyses to control for potential bias. The WM region calling agreed well with manually annotated WM regions on a subset of the data; however, it showed variable results for the GP region, which is specifically enriched in non-neuronal cells (Figure S10A).

### Matrix vs. striosome spatial gradients correlate with known SPN expression gradients, as well as novel interneuron-specific expression and methylation gradients

We were interested in identifying spatial gradients in gene expression and methylation across the BG, as these gradients may reflect functional differences between cells. We focused on a prominent spatial feature of the Striatum: the patterning of matrix and striosome SPNs. Matrix and striosome compartments were identified using local cell neighborhood composition vectors and each cell was assigned a value (MS Score) representing how far a cell is from the compartment boundary (Figure S10A and STAR Methods). SPN cell-type proportions in each compartment largely agreed with the compartment assignment (Figure S10B). Cell-type distribution along the matrix-striosome (MS) gradient showed expected enrichments of Matrix and Ventral SPNs in the matrix and Striosome SPNs in the striosome, however some unexpected enrichments were also apparent (Figure 6A, and S10C). Significant cell-type enrichments in either compartment were calculated using t-tests across brain regions, donors, and replicates (STAR Methods), at both the subclass and group levels. Again, SPN cell types showed the expected enrichment profiles. Additionally, novel enrichments were also identified, including the MGE-derived interneuron subclass CN ST18 GABA in the matrix, the CGE-derived interneuron subclass CN LAMP5-CXCL14 GABA in the striosome, and astrocytes in the striosome (Figure S10D-E). Group-level results showed that the enrichment of CN ST18 GABA cells in the matrix was driven largely by the STR TAC3-PLPP4 GABA group (Figure 6B). In contrast, the other two groups (STR-BF TAC3-PLPP4-LHX8 GABA and STR FS PTHLH-PVALB GABA) in this subclass showed no statistically significant distribution enrichment in either compartment (Figure S10F).

**Figure 6.**
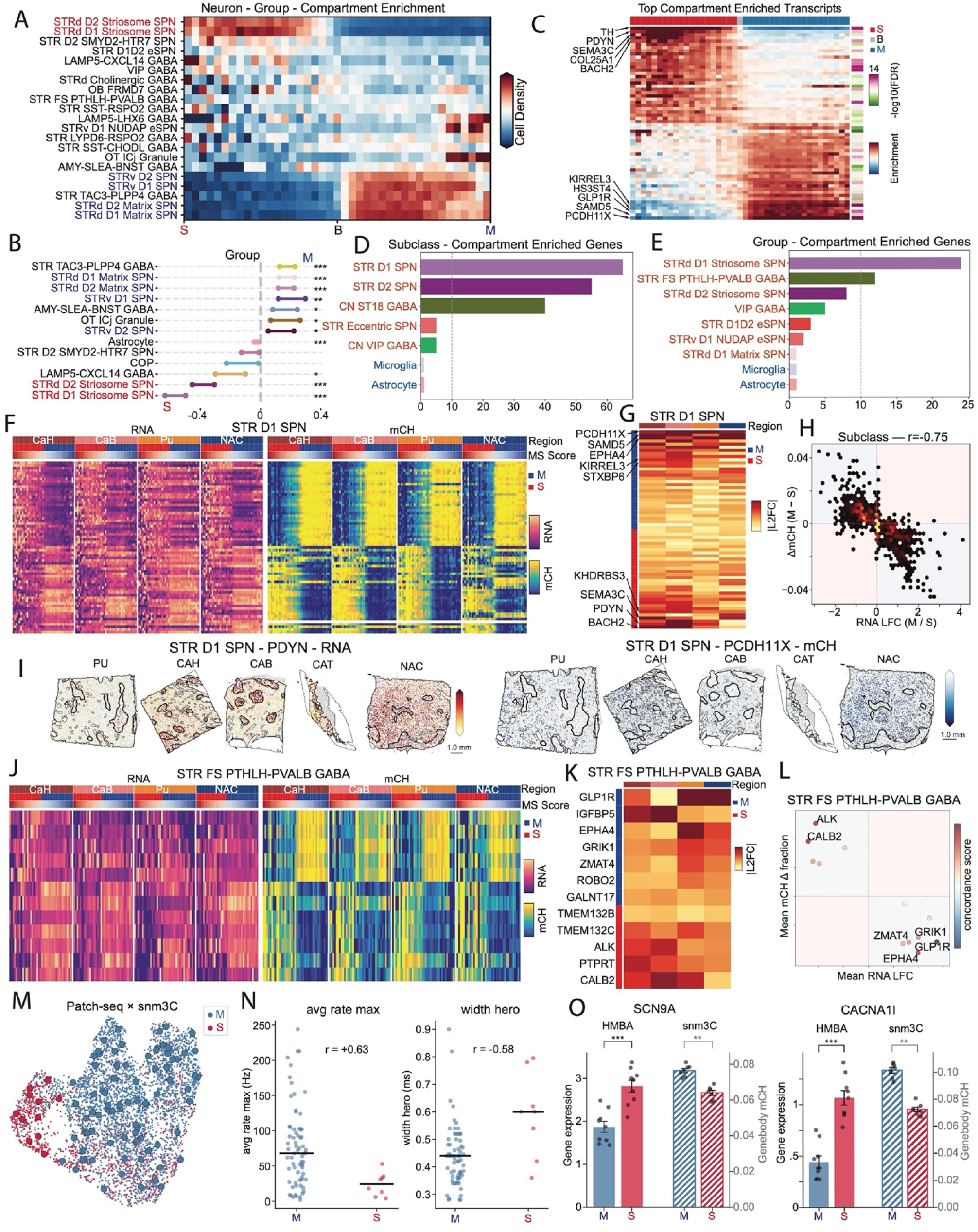
MERFISH reveals matrix-striosome compartmental differences in the striatum that extend beyond SPN identity. (A) Distribution of non-neuronal group cell-types along the matrix-striosome gradient. Each row is divided into 40 equal sized bins. Each bin is normalized by the total number of cells in the bin, and then each row is normalized to sum to 1. Bins are colored by that normalized cell density. Red signifies enrichment in those bins. Red color y-axis labels indicate canonical striosome localizing SPNs. Blue color y-axis labels indicate canonical matrix localizing SPNs. (B) Distribution of the top 30 enriched transcripts in the striosome (red) and matrix (blue) compartments. Enrichment was calculated like in (A), with red color signifying enrichment of transcript density. The right bar indicates significance of enrichment (-log10 of the FDR corrected p-value). Arrows point to the top 5 most enriched transcripts in either compartment. (C) Significant cell-type enrichments in matrix (right) or striosome (left) compartments at the group level. Y-axis labels are colored like in (A). The x-axis shows the 95% confidence intervals on the log2 fold-change of the normalized cell-type abundance in matrix and striosome compartments across regions, donors, and replicates. stars indicate FDR-adjusted significance: *<0.01, **<0.001, ***<0.0001. (D-E) Number of significant MS-DEGs within each cell type at the subclass (D) and group (E) levels. Y-axis labels are colored by cell-type neighborhood assignment. (F) Heatmap of MS-DEGs for the STR D1 SPN subclass. The left panel shows normalized RNA expression, and the right panel shows normalized mCH gene body methylation score. Rows are MS-DEGs for the subclass. The columns are 40 evenly spaced bins along the MS Score axis. Within each bin all cells are pseudobulked, and the pseudobulked value (sum of gene expression and average of mCH gene body levels) is shown. Columns are grouped into brain regions, and organized from left to right by MS Score. (G) Heatmap of the absolute log2 fold change of gene expression in the matrix versus the striosome, stratified by brain region, for the STR D1 SPN subclass. Rows are the MS-DEGs for this subclass, ordered by effect size. Top rows are matrix enriched genes, bottom rows are striosome enriched genes. Top genes are labeled via arrows. (H) Correlation between RNA log fold change (LFC) values of matrix (M) over striosome (S) expression with the difference in gene body mCH in the matrix minus the striosome. All subclass MS-DEGs are included. Color of spots indicates the number of genes that fall at that location in the graph. (I) Spatial distribution of example genes in the STR D1 SPN subclass. Only the striosome and matrix is shown. The black outlined regions are striosomes, the rest is the matrix. The gray background is all other cells in the regions. Left: measured RNA expression of *PDYN*, enriched in the striosome in all regions but the NAC. Right: imputed gene body mCH levels of *BACH2*, hypomethylated in the matrix. (J) Heatmap of MS-DEGs for the STR FS PTHLH-PVALB GABA group. The left panel shows normalized RNA expression, and the right panel shows normalized mCH gene body methylation score. Rows are MS-DEGs for the group. The columns are 40 evenly spaced bins along the MS Score axis. Within each bin all cells are pseudobulked, and the pseudobulked value (sum of gene expression and average of mCH gene body levels) is shown. Columns are grouped into brain regions, and organized from left to right by MS Score. (K) Heatmap of the absolute log2 fold change of gene expression in the matrix versus the striosome, stratified by brain region, for the STR FS PTHLH-PVALB GABA group. Rows are the MS-DEGs for this group, ordered by effect size. Top rows are matrix enriched genes, bottom rows are striosome enriched genes. (L) Mean RNA LFC values of matrix over striosome vs. mean gene body mCH differences in the matrix minus the striosome. Means taken over brain region specific values. spots are colored by concordance score, gene level correlation across donors and brain regions. (M) Integration of STR FS PTHLH-PVALB GABA neurons from snm3C dataset with the same population from the macaque Patch-seq data. Smaller cells are from snm3C, bigger outlined cells are from Patch-seq. snm3C cells are colored by predicted localization from MERFISH integration. Patch-seq cells are colored by predicted localization via label transfer from the snm3C cells. (N) Select electrophysiological differences between striosome and matrix predicted STR FS PTHLH-PVALB GABA neurons from macaque Patch-seq data. R values shown correspond to the rank-biserial effect size. Black bars are the medians. Left: average maximum firing rate. Right: Peak width of an action potential during a hero (long square pulse current with the fastest firing rate) sweep. (O) Barplots showing the increased expression (HMBA - left) and mCH hypomethylation (snm3C - right) of *SCN9A* and *CACNA1I* in striosome predicted STR FS PTHLH-PVALB GABA neurons. Matrix (M - blue). Striosome (S - red).

We next sought genes that exhibit differential expression across MS boundary. The same approach as above was applied using all detected transcripts within either the matrix or striosome boundary. An MS Score was assigned to each transcript, the enrichment test was repeated (STAR Methods), and enrichments were visualized for the top 30 enriched genes in either compartment (Figure 6C). Top enriched genes in the striosome include *TH, PDYN, COL25A1, BACH2*, and *MOXD1*. The top enriched genes in the matrix include *PCDH11X, SAMD5, GLP1R, HS3ST4*, and *KIRREL3*. GO term analysis showed no significant enrichment in either gene set (STAR Methods), however the union of the gene set was enriched in synapse and dendritic-related processes.

Next, cell-type-specific matrix-striosome differentially expressed genes (MS-DEGs) were identified using a DESeq2-based approach. Compartment-stratified, pseudobulked, and cell-type specific expression profiles were generated for each striatal brain region. PyDESeq2 was applied, with donor as a covariate, and stringent significance cutoffs were applied (STAR Methods). The highest number of MS-DEGs was observed in SPNs and CN ST18 GABA at the subclass level (Figure 6D), likely due to the differenes in compartment composition among the groups within those subclasses. At the group level, the striosome SPNs had the most MS-DEGs, likely due to the conservative striosome boundary estimates and leaky SPN annotations. Surprisingly, another CN ST18 GABA group, STR FS PTHLH-PVALB GABA, showed the second-highest number of MS-DEGs. (Figure 6E). Among the top MS-DEGs identified for D1 and D2 SPNs were known marker genes for the striosome and matrix groups, including *BACH2*, *PDYN*, *EPHA4*, *KIRREL3*, and *PCDH11X*. These detected gene expression gradients were strongly correlated with imputed methylation gradients (Figure 6F and S10G). Fold-change estimates for all MS-DEGs were compared across brain regions, revealing genes whose compartmentalization strength was regionally dependent (Figure 6G, S10H). The region-specific effects were highly concordant between RNA and gene-body mCH (Figure 6H, STAR Methods). Spatial plots of gene expression and imputed methylation for the top matrix and striosome marker genes in the SPN subclasses also showed strong compartmental patterning and agreement with our identified regional differences (Figure 6I, S10I). Reasoning that the enrichment of the STR TAC3-PLPP4 GABA group in the matrix likely accounted for most CN ST18 GABA MS-DEGs, we next focused on the STR FS PTHLH-PVALB GABA group as the only non-SPN neuronal group with MS-DEGs. A repeat of the above analysis revealed similar patterning, albeit to a lesser extent, of regionally specific compartmental genes (Figure 6J-K, S10J), and strong correlation between fold change in expression and difference in mCH gene body methylation in the MS-DEGs (Figure 6L). Reclustering the STR FS PTHLH-PVALB GABA group by itself and projecting the imputed compartmental location of the snm3C and HMBA cells onto their respective embedding spaces (Methods) revealed a striosome-enriched subtype apparent in all 3 datasets (Figure S11A-B). Integration of the snm3C and HMBA STR FS PTHLH-PVALB GABA cells showed that the striosome and matrix localization predictions were consistent across modalities (Figure S11C), further enforcing the consistent nature of these results in the MERFISH, snm3C, and HMBA datasets.

For functional validation of the identified differences in STR FS PTHLH-PVALB GABA neurons, Patch-seq data were integrated from a companion paper (Liu et al^50^). 89 STR FS PTHLH-PVALB GABA neurons from macaque brains were integrated with the HMBA and snm3C datasets independently, and the matrix or striosome spatial localization was inferred using each dataset as a bridge from the MERFISH data (Methods, Figure 6M, S11D). Of the 89 neurons, 9 were unanimously identified as striosome-localized, 72 as matrix-localized, and 8 had conflicting identities and were therefore removed from downstream analysis. Clustering of the 89 Patch-seq cells showed no enrichment of matrix or striosome cells within de novo clusters (Figure S11E).

Previously described attributes of STR FS PTHLH-PVALB GABA interneurons were used to assess the integration quality across the 4 datasets (MERFISH, HMBA, snm3C, and Patch-seq). First, enrichment of these neurons was confirmed within the caudate as compared with the putamen^14^ (Figure S11F). Next, *PVALB*+ and *PVALB*- subtypes were identified, and global *PTHLH* expression was confirmed^14,72^, noting that the split in PVALB expression did not correspond to the matrix-striosome split (Figure S11G). Lastly, *MOXD1* was identified as a strong marker gene for the striosome-specific population (Figure S11H-I). A *MOXD1* subtype of human FS *PTHLH*+ interneurons in the striatum was previously identified^14^, however that study lacked spatial data to assign anatomical meaning to the subtype.

Using the high-confidence anatomical assignments of the Patch-seq cells, clear electrophysiological differences were observed between the predicted matrix and striosome-localized cells (Figure S11J, STAR Methods). Specifically, matrix STR FS PTHLH-PVALB GABA neurons exhibited a faster firing rate, shorter action potential width, and lower excitability, suggesting a more canonical fast spiking phenotype (Figure 6N). The anatomical assignments on the HMBA dataset were then used for differential expression analysis, which was then validated in the snm3C mCH gene body levels (Methods). Several differentially expressed genes supported the identified electrophysiological differences, including higher *SCN9A* and *CACNA1I* expression in striosome cells, which are involved in modulating excitability^73^ and burst firing patterns^74^, respectively (Figure 6O). Other differentially expressed genes include *GRIK1/3*, *GALNT10/17*, and *ADRA1D* (Figure S11K).

## Discussion

Together with the BICAN community, we have generated a multi-omic atlas of the human BG by annotating cells across modalities to a shared consensus BG cell-type taxonomy. This framework enables direct cross-modality comparisons for each annotated cell type across five layers, RNA, ATAC, DNA methylation, Hi-C, and histone modifications, of which DNA methylation and Hi-C are contributed by the present study. The resulting resource provides a foundation for integrated analyses and for developing machine learning or deep learning models that leverage coordinated regulatory signals across modalities. Our analysis of the DNA methylation landscape in the human BG suggests that cells exhibit a strikingly distinct pattern of methylation at genomic loci outside of gene bodies and promoters. Of all cell types, SPNs showed the greatest similarity in their cell-type-specific hypo-DMR patterns, suggesting that although different SPN subtypes are known to be involved in distinct pathways (e.g., direct vs. indirect pathways), they are highly similar from an epigenetic perspective. Previous work has shown that differentiating regulatory domains to specific cell-types labels enables LDSC analysis to map GWAS hits to those cell types. We extend these analyses to our comprehensive BG atlas. While we did not see the expected enrichment of PD risk in the dopaminergic neurons, potentially due to insufficient sampling of the dopaminergic centers, our results suggest an indirect role for F M GATA3 GABA subclass in PD at the subclass level. A possible reason for this is SN GATA3-PVALB GABA neuronal dysfunction in PD^75^, which modulates circuit imbalance via projections to the ventral motor thalamus, motor layers of the superior colliculus, and mesencephalic locomotor regions^76^. Another potential mechanism for this enrichment is through the RN GATA3 GABA group. Previous literature has shown that compensatory mechanisms in the RN can mask PD symptoms well after neurodegeneration has taken effect^77^, and that these compensatory mechanism are hereditary in marmosets. Whether these genetic links reflect causal contributions to PD pathology or correlation with disease severity remains to be determined. These results, when combined with prior literature, suggest a compensatory mechanism in PD mediated by non-dopaminergic cell types. Additionally, the substantial overrepresentation of GWAS hits in hyper-DMRs across many cell types for psychiatric diseases such as schizophrenia and bipolar disorder merits further investigation.

We also reported associations between PD and hypo-DMRs of STR SST-CHODL GABA at the subclass level and loop anchor of STR SST-CHODL GRIN2A GABA at the group level. A recent study reported a reduction in both the number of SST+ INTs and SST mRNA levels in *PARK2*-specific induced pluripotent stem cell (iPSC)-derived GABAergic neurons from PD patients^78^. Additionally, *PARK2* mutations in *SST*+ INTs have been shown to decrease *SST* transcript levels and induce mitochondrial dysfunction, potentially leading to an excitatory/inhibitory imbalance that contributes to the motor and non-motor symptoms characteristic of PD^79^. Moreover, somatostatin (SST) has been reported to play a key role in modulating the feedback-inhibitory circuit among SPNs^80^. Our 3D genome analyses are consistent with and extend prior observations that neuronal chromosomes are relatively more compact at local scales but less compact at larger scales, reflecting increased short-range and reduced long-range interactions in neurons^48^. Previous work reported reduced intra-compartment (A-A and B-B) and increased inter-compartment (A-B) interaction frequencies in neurons^48^, but did not examine how contact-distance distributions vary within each compartment class. Although long-range contacts have been observed to be more prevalent in non-neuronal cells^33,42^, how these long-range interactions are partitioned across A/B compartment combinations remains unclear.

Here, we confirmed that non-neuronal cells exhibit longer contact distances overall, whereas neuronal cells were enriched for long-range interactions within A-B contacts, indicating that long-range connectivity in neuronal genomes is largely driven by inter-compartment mixing. Pletenev et al. reported that TAD structures are more pronounced in neurons, with higher contact density at the TAD scale^48^, consistent with the enrichment of short-range interactions in neuronal cells; our results corroborate this observation. Likewise, we also observed that chromatin loops are substantially longer in neurons than in non-neuronal populations. Notably, despite the increased fraction of long-range contacts in non-neuronal cells, loop distances are shorter, suggesting that long-range contacts in non-neuronal genomes are not primarily organized as focal loops rather long-range inter-compartment interactions. We also identified genes that undergo compartment switching between different cell types. The cases where genes showed high expression within B compartments were likely related to the resolution of our compartments; 100kb regions are larger than most genes and regulatory domains, and have recently been shown to be larger than most compartments in bulk HiC data^81^. However, the switch from an A compartment to a B compartment at 100kb resolution most likely reflects a change in the local DNA around the gene from a more poised, active state to a more repressed, inactive state.

Using both DNA methylation and HiC data, we resolved medium spiny neurons into group-level subtypes, identified a novel cell type (D2 SMYD2) with consistent gene-centric differences across all studied modalities, reported subtype-specific hypomethylated marker genes, and identified regional DMGs that were further supported by our spatial transcriptomic data. A recent study identified *POU6F2* as a marker of a hindbrain-derived (rhombomere r1) SNr projection-neuron subclass co-expressing *PAX5* and *ZFPM2*^56^; D2 SMYD2 shares *POU6F2* and *ZFPM2* with this SNr subclass but lacks *PAX5*, consistent with its LGE-derived telencephalic origin. This raises the possibility that a conserved *POU6F2/ZFPM2* module has been recruited independently in two developmentally divergent GABAergic projection-neuron lineages within basal-ganglia circuits, although lineage tracing and conditional *POU6F2* perturbation will be needed to confirm this. Our multi-modal analysis revealed that 3D genome embeddings could differentiate the dorsal-ventral axis of SPN variability, which stands in direct contrast to the results presented in the companion manuscript^35^, which showed that the dorsal-ventral SPN axis had the least variability from a transcriptomic perspective. Our analysis of regional DMGs revealed many genes exhibiting patterned methylation, 3D dynamics, and altered gene expression across the SPN subtypes in the striatum. These regional differences help identify a previously unknown differential expression gradient associated with sub-striatal specificity in disease susceptibility. In the case of ADSD with a CaT-specific expression and hypomethylation of the known causal gene *PDE8B*; in PD, NAC-specific expression and hypomethylation of CADM1, a GWAS-associated biomarker linked to PD motor progression, with proposed mechanisms involving dopaminergic signalling, synaptic cell adhesion (potentially through Necl-1:Necl-2 interactions), and receptor recycling^82^. We report associations between specific SPN subtypes and ALS. This link needs further validation due to specific caveats in the LDSC algorithm (STAR Methods) and because our data lacked cells from the motor cortex cells, which are known to be significantly associated with ALS^83^. Nonetheless, SPNs serve as the primary input nodes to the BG from the motor cortex within cortico-BG-thalamus-cortical loop, and their potential role in ALS pathology warrants further investigation. Our study of the epigenomes of SPNs provides a reference for studies of striatal cellular diversity and regional specialization in the human BG.

At the regulatory element level, millions of DMRs showed strong cell-type specificity and were accompanied by enriched TF motifs and candidate regulators. By jointly integrating DNA methylation, ATAC-seq, H3K27ac, Hi-C, and RNA profiles and linking TF motifs within DMR-overlapping enhancers to predicted target genes, we inferred enhancer-promoter interactions and constructed cell-type-specific gene regulatory networks. Some of our predicted enhancers recapitulate elements from the enhancer-AAV toolbox, which were identified from human and mouse ATAC-seq datasets as cell-type-restricted accessible regions near striatal marker genes and validated in vivo in mouse, providing cross-species computational support that these regulatory elements act in matching cell types in the human basal ganglia; others are novel candidates not previously reported. While these TF-target relationships are supported by multi-omic coherence and external TFBS resources, functional validation in BG contexts will be an important next step. We also identified enriched TF binding motifs within the predicted enhancer regions, which may regulate target gene expression in a cell-type-specific manner. The discovery of these cell-type-specific enhancer-promoter interactions enables us to determine whether PD-associated GWAS SNPs reside in enhancers active in particular M Dopa subtypes, an observation further supported by prior studies^65,66^. We point out that transient TF expression may pose an obstacle to the development of GRNs when snRNA-seq is used as the sole marker of TF activity. Persistent hypomethylation of gene bodies can serve as a secondary, and perhaps more reliable marker for TF activity in the context of GRN construction. This paper extends previous work^32^ on method development for building GRNs using extensive multimodal data in a cell-type-specific manner.

By integrating spatial data with the HMBA reference taxonomy, we showed that unique interneuronal cell types preferentially localize to either the matrix or striosome compartments in the Striatum. The TAC3-expressing, MGE-derived, interneuron type localized to the matrix, and CXCL14-expressing, CGE-derived interneurons were localized to the striosome. This was coupled with correlated differential expression and methylation of select genes within the same cell types, stratified by compartment. We identified two subtypes of STR FS PTHLH-PVALB GABA interneurons, a finding that was supported across 4 datasets and modalities. The identified, spatially restricted fast-spiking subtypes showed epigenomic, transcriptomic and electrophysiological differences, which pointed to the striosomal subtype as having less canonical fast-spiking properties. Furthermore, the identification of *MOXD1* as a subtype-specific marker aligns with previous literature^14^, and incorporates the anatomical and electrophysiological differences identified in this study. All together, these results suggest that the matrix and striosome are not differentiated solely by the identity of the SPNs within them, but also by a local microenvironment that supports their distinct identities and functions. Our analysis was limited by the size of the gene panel used; further validation with an expanded gene set is needed to determine whether other cell types also exhibit this transcriptional and epigenetic patterning.

This atlas and accompanying multi-omic analyses provide a comprehensive, layered regulatory framework for the human BG, integrating DNA methylation, chromatin accessibility, histone modifications, 3D genome architecture, transcriptional output, and spatial localization to illuminate cell-type-specific mechanisms that govern neuronal diversity. By linking these epigenetic states to gene expression programs, we predicted thousands of enhancer-promoter interactions; some of those predictions were supported by experimentally reported enhancers. Critically, our integrative approach revealed that disease-associated genetic variants, particularly those linked to basal ganglia disorders such as PD, preferentially localize to cell-type-specific enhancer regions, offering mechanistic insights into how risk variants disrupt specific neuronal subtypes and contribute to disease susceptibility. This multi-layered regulatory map thus serves as a valuable public resource for interpreting non-coding genetic variation, dissecting BG pathophysiology, and guiding targeted experimental follow-up and therapeutic strategies in neurodegenerative disorders.

### Limitations of the Study

This study has several limitations that should be considered when interpreting the results. First, the snm3C-seq dataset is derived from seven postmortem donors of different ages and sexes, which limits statistical power to disentangle donor-specific variation from true biological effects, such as age- or sex-related differences, and may affect the generalizability of our findings. Additionally, the snm3C-seq dataset is enriched for neurons, with neurons comprising 90% of the dataset, which does not reflect the true proportions of cells in these brain regions. Second, although cell-type clusters were identified de novo from methylation features and annotated by alignment to the HMBA taxonomy, the 100-kb resolution of Hi-C contact maps may miss finer-scale chromatin architectural differences between more closely related subtypes. Third, the newly identified D2 SMYD2 subtype, though consistently supported across DNA methylation, Hi-C, RNA, histone-modification, and electrophysiological modalities (including macaque Patch-seq), would benefit from further perturbation-based functional validation to directly test the lineage and functional roles inferred here. Fourth, the enhancer-promoter links and TF-target gene regulatory networks reported here are computationally predicted based on chromatin accessibility, 3D genome, motif analyses on DMRs and correlation. The enhancer-AAV toolbox elements recapitulated in our analysis were functionally validated in vivo in mice, not in primate or human tissue. Confirming their cell-type specificity in human basal ganglia will require future in vivo validation in non-human primates or post-mortem human systems. Functional validation of the predicted enhancer-promoter links, for example, through CRISPRi perturbation in iPSC-derived striatal neurons, represents an important future direction that could establish causal regulatory relationships. Fifth, disease-associated cell-type enrichments identified through LDSC are correlative, and whether disease-associated variants in the implicated cell types functionally alter gene expression or chromatin state has not been tested. LDSC enrichment estimates for cell types whose DMR annotations cover only a small fraction of the genome, for example, group-level DMRs for many cell types, can yield elevated false-positive results and should be interpreted with caution given the limited number of overlapping SNPs. Additionally, the unexpected PD enrichment in F M GATA3 GABA neurons rather than dopaminergic neurons may partly reflect the limited sampling of dopaminergic cells in our dataset. Sixth, the MERFISH spatial analysis was constrained by the size of the gene panel, which limited its ability to annotate certain cell types. Additionally, the matrix and striosome composition analysis was inconsistent in the CaT sections due to the limited number of striosome patches in those sections. Furthermore, the STR FS PTHLH-PVALB GABA Patch-seq data came from macaque, and therefore, the identified electrophysiological differences may not generalize to humans. Finally, our analyses are largely descriptive and correlative; functional experiments will be required to establish causal links between the cell-type-specific epigenomic features described here and gene regulation, cell identity, and disease susceptibility.

## STAR★Methods

### Key resources table

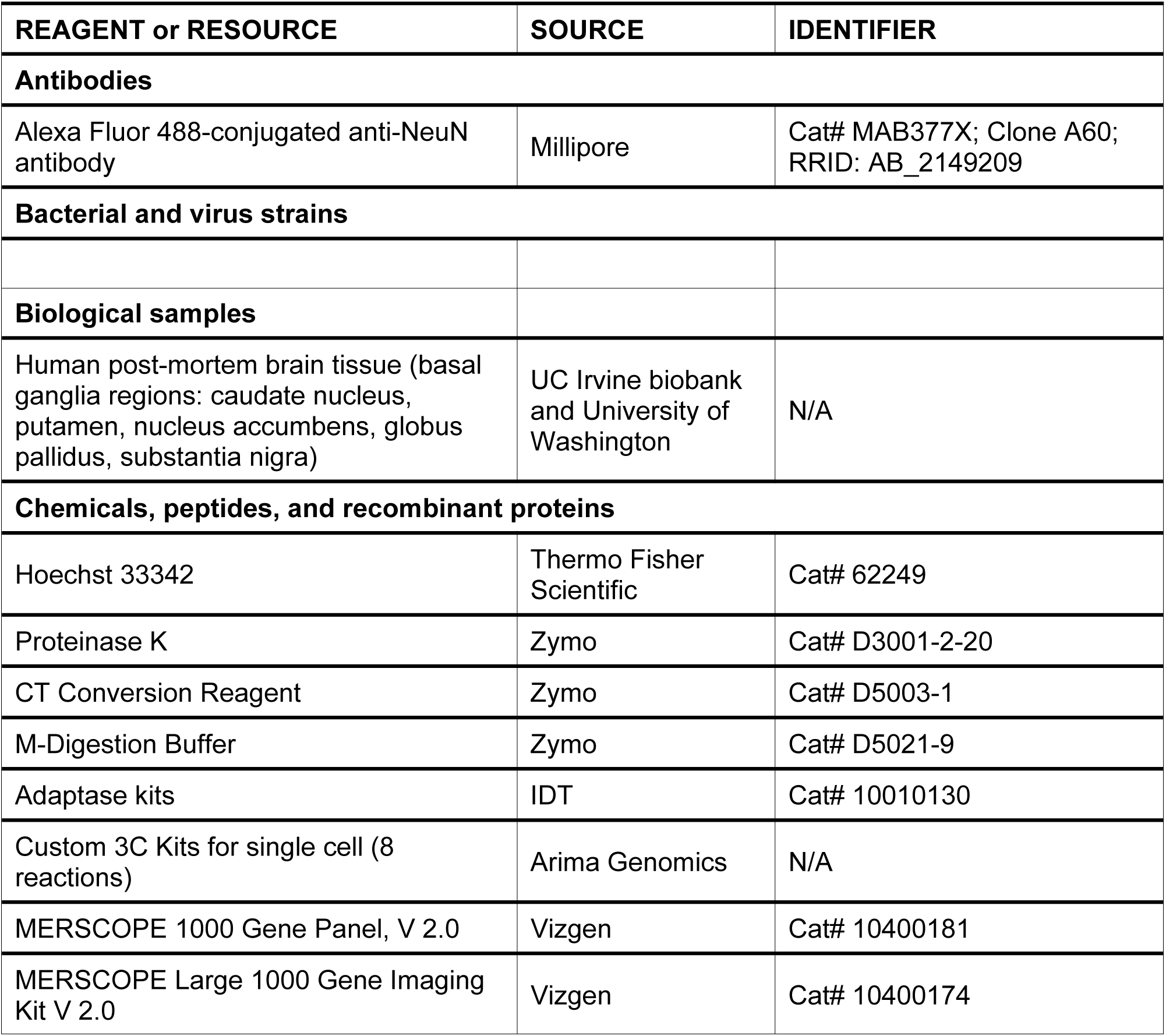
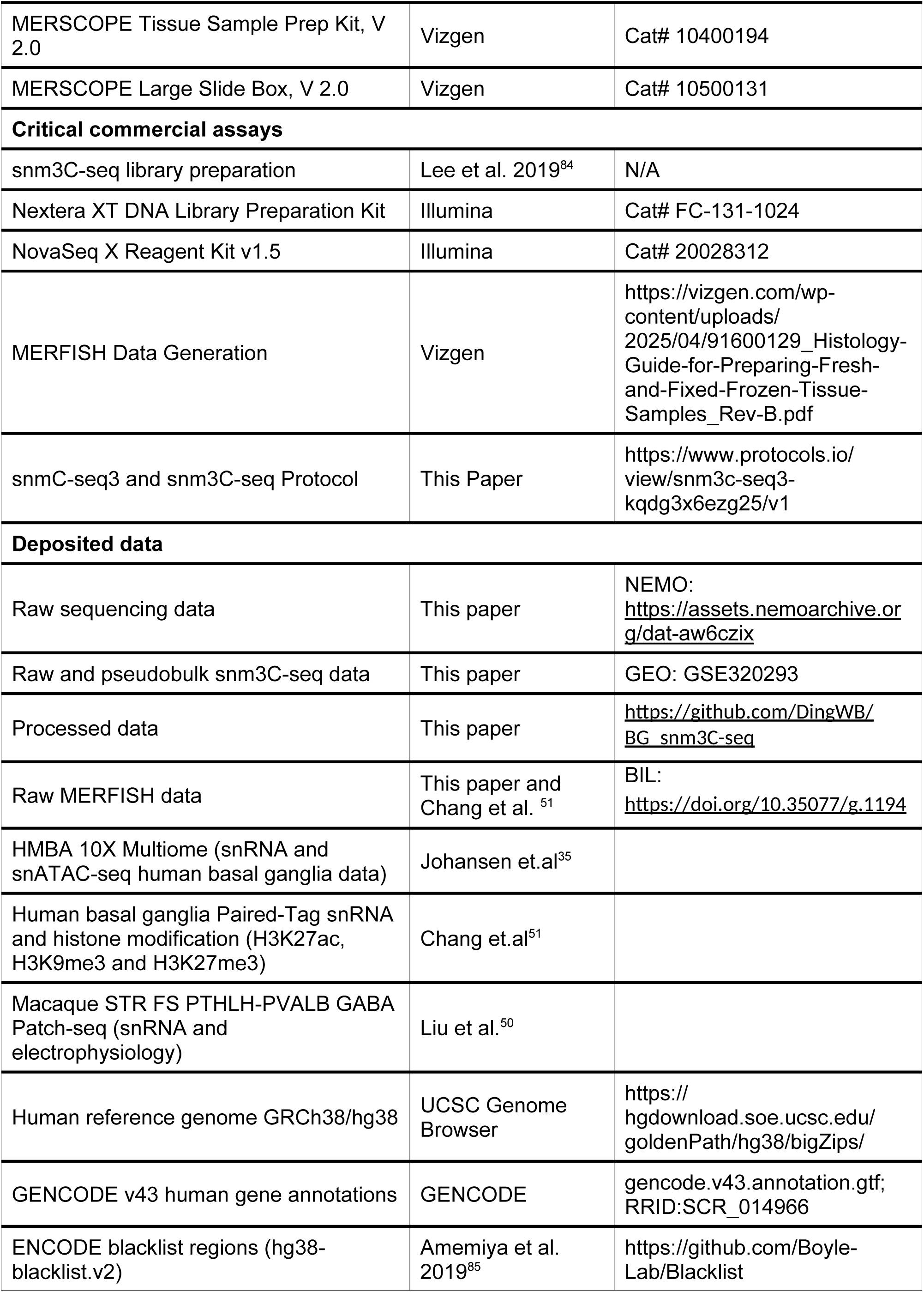
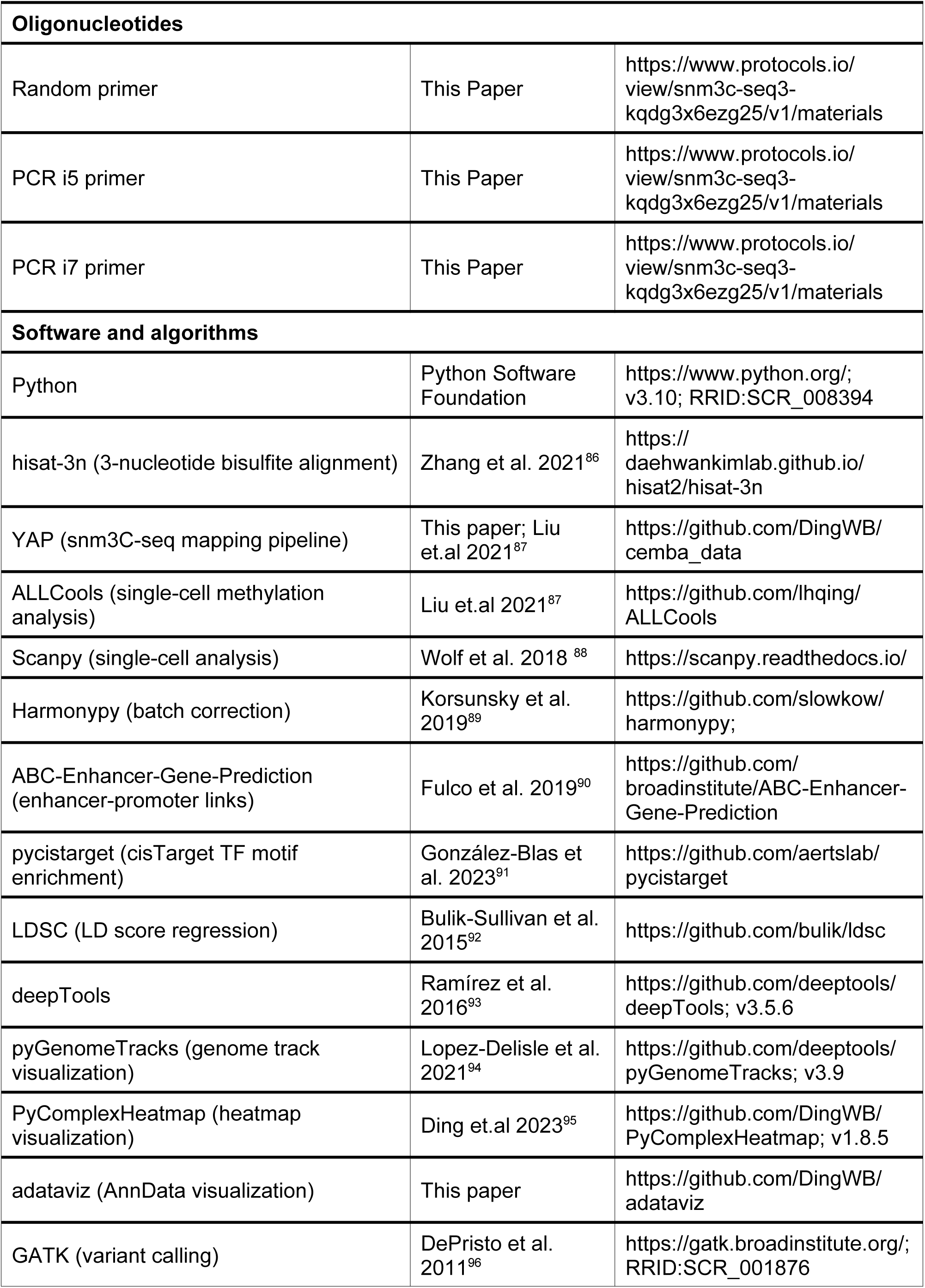
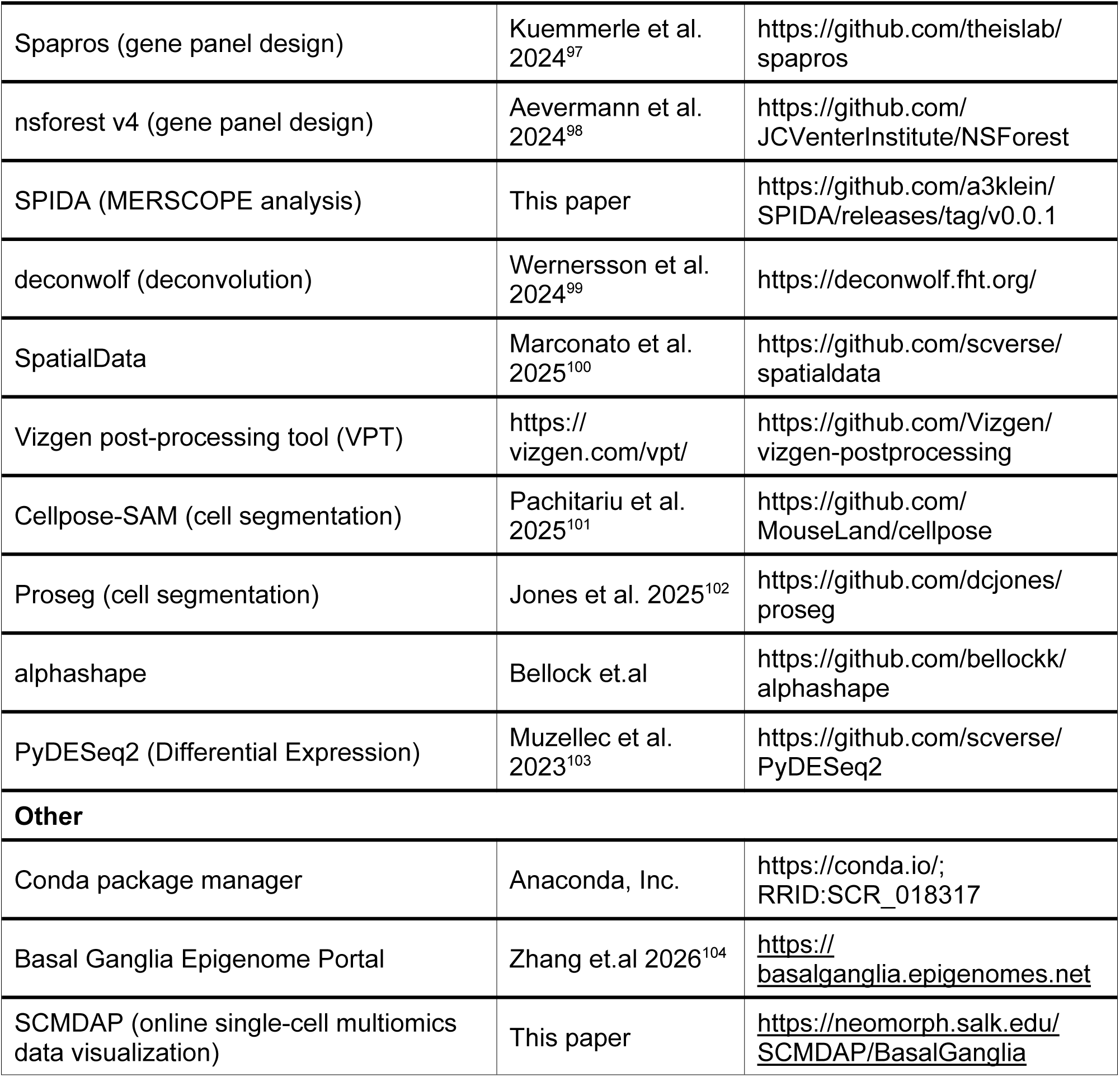

### Donors collection

Donors 29 - 68 years of age with no known history of neuropsychiatric or neurological conditions were considered for inclusion in this study. Authorization for brain donation was obtained from each donor’s legal next of kin, donor brain tissue was de-identified, and collection of postmortem brain tissue was collected in accordance with the United States Uniform Anatomical Gift Act of 2006 and all other applicable state and federal laws and regulations. Blood samples were collected from each donor for routine serological screening of infectious diseases (HIV, Hepatitis B, and Hepatitis C) and only donors testing negative for all three diseases were further considered for inclusion in the study. For each donor brain, frozen tissue samples were collected from the frontal and occipital poles, the cerebellar cortex, and the midbrain to access RNA quality. The RNA was isolated using the RNeasy Lipid Tissue Mini Kit (Qiagen 74804), and RNA Integrity Number (RIN) was determined using the Agilent 4150 Tapestation System (Agilent G2992AA). Only donor brains with RIN >6 across all assessed regions were used in the study, except for the UCI4723 cerebellar cortex (5.1) and the UCI2424 occipital cortex (5.5). Donor brain tissue used in this study was obtained from the UC Irvine biobank (donor IDs: UCI2424, UCI5224 and UCI4723) and the BioRepository and Integrated Neuropathology (BRaIN) Laboratory at the University of Washington (UWA7648) (Table S1).

This study featured tissue from four human donors (two males, 50 years, right hemisphere, and 47 years, left hemisphere; and two females, 63 and 67 years, right hemisphere) with post-mortem intervals (PMI) of 16.3, 25.5, 23.5, and 19.8 hours, and RIN scores of 9.6, 6.8, 6.3, and 8.2, respectively (Table S1). Specimens were requested through the BICAN Specimen Portal (available from https://brain-specimenportal.org/), which is part of the Neuroanatomy-anchored Information Management Platform for Collaborative BICAN Data Generation (NIMP; RRID:SCR_024684). NIMP was used to manage specimen metadata and workflow tracking from donor tissue to downstream processing and data deposition to modality-specific archives, including The NeMO Archive^105^ and NIMP^106^.

### Slabbing

Fresh human donor brains were processed according to the Allen Institute post-mortem brain processing procedure, with slight modifications (https://dx.doi.org/10.17504/protocols.io.bf4ajqse). Briefly, the cerebrum was separated from the cerebellum and brainstem, bisected into right and left hemispheres, and one cerebral hemisphere from each donor was slabbed coronally at a thickness of ∼4 mm. Fresh slabs were flash frozen and stored in vacuum-sealed bags at −80°C. All slabs were photodocumented both when fresh and after flash-freezing.

### Anatomical criteria

Frozen 4 mm coronal slabs from whole human post-mortem brains were aligned to an interactive, digitally annotated human brain atlas (Ding SL et al., 2016). For each brain, eight basal ganglia regions of interest (ROIs) were anatomically delineated and microdissected according to the following criteria (1) For head of Caudate nucleus (CaH) region: after alignment to the Ding Atlas, the full-thickness of the slab containing the ROI -encompassing plates 16 to 18 of the atlas^107^ was dissected, minimizing contamination from the adjacent regions such as Nucleus Accumbens (NAC). (2) Body of caudate nucleus (CaB) region: after alignment, the full-thickness of the slab containing the selected ROI, (Ding Atlas plates 20 to 33), minimizing contamination from the anterior CaH region. (3) Tail of caudate nucleus (CaT) region: after alignment, the full-thickness (Ding Atlas plates 49 to 69), minimizing contamination from the adjacent region Hippocampus (HIP) and white matter. (4) Putamen (Pu) region: after alignment, the full-thickness (Ding Atlas plates 17 to 27); minimizing contamination from adjacent Globus Pallidus (GP), Claustrum (Cla) and NAC regions. (5) Subthalamic nucleus (SubTH: STH) region: the full-thickness (Ding Atlas plates 37 to 49); minimizing contamination from adjacent regions from the thalamus and midbrain. (6) Globus Pallidus (GP) region: the full-thickness (Ding Atlas plates 22 to 42); minimizing contamination from the adjacent and anterior Pu region and Basal forebrain structures. (7) Nucleus accumbens (NAC) region: The full-thickness (Ding Atlas plates 18 to 22); minimizing contamination from adjacent Pu, CaH, BNST, A25 regions and anterior CaH region. (8) Gray matter of midbrain 1 (MGM1) Region: this sample was collected from the first and second slab of midbrain and used as a landmark for the presence of Substantia Nigra (SN), including regions SN, Red Nucleus (RN), and ventral tegmental region of midbrain (VTR) (FigureS1A).

### Microdissection

Tissue blocks were microdissected on an aluminum block surface inside a Leica cryostat maintained at −23°C. Guided by the Anatomical criteria and Ding atlas reference images, regions of interest were cut using a sterile razor blade (Genesee, 38-101) with slow, perpendicular see-saw motions to minimize cracking of frozen tissue. Microdissected tissue was distributed into a full-thickness 7×7 mm tissue block for spatial transcriptomics and adjacent tissue chunks for single-nucleus isolation. The required tissue mass for single-nucleus isolation was determined for each brain region based on neuronal density, with a target of isolating 1 million neuronal nuclei per region, split between the two multiomic modalities used in this study and the companion study (Chang et al. co-submit): snm3C-seq and Droplet Paired-Tag, respectively. Tissue mass required per region was determined to be as follows: CaH (270 mg), CaB (320 mg), CaT (400 mg), Pu (248 mg), STH (300 mg), GP (1,200 mg), NAC (118 mg), and MGM1 (900 mg), where CaT, STH, GP, and MGM1 samples used all available tissue mass and recovered less than the target amount of neuronal nuclei (See “Nuclei isolation” section below).

### Cryosectioning for MERFISH 2.0 and NISSL staining

Tissue blocks were embedded in Tissue-Plus™ O.C.T. Compound (Fisher Scientific, 23-730-571), sectioned at 10 μm on a Leica Cryostat (Leica CM1950) using Epredia™ MX35 Premier™ low-profile blades (VWR, 89238-778), and mounted onto MERSCOPE Large Slide V 2.0 (Vizgen; 20400118). Sections were fixed in 4% paraformaldehyde (Electron Microscopy Science, 15714-S) in 1X Phosphate-Buffered Saline (PBS, Thermo Fisher Scientific, AM9625) at 47 °C for 30 min, permeabilized overnight in 70% ethanol and photobleached for 5 hours using a MERSCOPE photobleacher (Vizgen; 10100003) (See MERFISH sample preparation section below). Additional sections were collected on Fisherbrand™ Superfrost™ Disposable Microscope Slides (Fisher Scientific, 12-550-123), fixed in 4% paraformaldehyde (Electron Microscopy Science, 15714-S) in 1X Phosphate-Buffered Saline (PBS, Thermo Fisher Scientific, AM9625) at 4 °C for 15 min and stained with 0.04% cresyl violet acetate working solution (Fisher Scientific, cat. no. 50-318-82) for 20 min, rinsed in distilled water, dehydrated through graded ethanol solutions (50%, 95%, 100%), cleared in Xylene solution (Fisher Scientific, X5-4), and cover slipped for histological analysis.

### Tissue Grinding

Frozen tissue dissections containing common brain regions were pooled across multiple slabs when necessary and ground with a porcelain mortar (VWR 470149-118) and pestle (VWR 470149-080) over dry ice. Ground tissue was distributed into aliquots of 150-250 mg in 1.5 mL microcentrifuge tubes on dry ice and stored at −80°C until further processing.

### Nuclei isolation and fluorescence-activated nuclei sorting (FANS)

Recombinant RNase inhibitor (RNH1) was purified from the plasmid pmal_c5x_RNAse_Inhib (Addgene plasmid # 153314; RRID: Addgene_153314), a gift from Drew Endy & Philippa Marrack, by the QB3 MacroLab at UC Berkeley and reconstituted at a concentration of ∼10 mg/mL.

Ground frozen tissue was homogenized in 9 mL of lysis buffer (0.32M sucrose, 0.1% Triton X-100, 5 mM MgCl_2_, 25 mM KCl, 10 mM Tris-HCl pH 8.0, 1 mM DTT, Roche COmplete Mini Protease Inhibitor, 20 U/mL SUPERaseIN, and 20 μg/mL RNH1) using glass Dounce homogenizers and sequentially filtered through 70 μm (Corning, 352350) and 30 μm (Sysmex, 04-004-2326) cell strainers.

Myelin debris was then removed by adding a sample-specific volume of myelin removal beads (Miltenyi Biotec, 130-096-433) to the homogenate, incubating on a rotating mixer at 4°C for 15 min, and passing the resulting suspension through LS columns (Miltenyi Biotec, 130-042-401) loaded onto a QuadroMACS Separator (Miltenyi Biotec, 130-091-051). The sample-specific volume of myelin removal beads was determined for each sample based on the myelin content of the tissue used. Flow through was collected and columns were washed with 2 mL of wash buffer (0.1% Triton X-100, 5 mM MgCl_2_, 25 mM KCl, 10 mM Tris-HCl pH 8.0, 1 mM DTT, Roche COmplete Mini Protease Inhibitor, 20 U/mL SUPERaseIN, and 20 μg/mL RNH1). Samples were centrifuged at 1,000g for 10 min at 4°C in a Thermo Scientific Sorvall Legend RT+ benchtop centrifuge (Thermo 75004377) with swinging bucket rotor (Sorvall 75006445). The supernatant was carefully aspirated and the pellet was resuspended in 1 mL of sort buffer (DPBS, 1% BSA, 1 mM EDTA, 1mM DTT, Roche COmplete Mini Protease Inhibitor, 20 U/mL SUPERaseIN, and 20 μg/mL RNH1). Mouse anti-NeuN conjugated to Alexa Fluor 488 (Millipore MAB377X) and Hoechst 33342 were added at a concentration of 1:1000 and nuclei suspensions were incubated on a rotating mixer at 4°C for 30 min. Nuclei suspensions were diluted to a concentration of 750K nuclei/mL using an additional sort buffer ahead of FANS.

Single nucleus sorting was carried out at 4°C on a BD Influx cell sorter (BD Biosciences 646500) using the ‘1-drop pure’ sort mode with a 100 μm nozzle and sheath pressure of 20 PSI. Gating was performed first on Hoechst-positive nuclei, followed by further exclusion of debris and nuclei aggregates using forward and side scatter pulse area and width parameters, and finally on NeuN signal. NeuN-positive and NeuN-negative nuclei were sorted into separate 2 mL DNA LoBind tubes (Eppendorf 022431048), each containing 250 μL 5X collection buffer (DPBS, 5% BSA, 5X Roche COmplete Mini Protease Inhibitor, 100 U/μL SUPERaseIN, and 100 μg/mL RNH1). All collected nuclei were then distributed between the two multiomic modalities, with 75% used for Droplet Paired-Tag in the companion study (Chang et al., co-submit) and 25% used for snm3C-seq.

### snm3C-seq

Sorted nuclei were pooled at a defined ratio of 90% NeuN-positive and 10% NeuN-negative, centrifuged at 500g for 5 min at 4°C in an Eppendorf microcentrifuge (Eppendorf 5417R) with swinging bucket rotor (Eppendorf A-8-11), and frozen in 250 μL of freezing media (DPBS, 2% BSA, 10% DMSO, 20 U/mL SUPERaseIN, and 40 U/mL RNaseOUT) at −80°C until processing for snm3C-seq.

A detailed snm3C-seq3 library preparation protocol is available at https://www.protocols.io/view/snm3c-seq3-kqdg3x6ezg25/v1. Briefly, presorted frozen nuclei were thawed to 4°C and centrifuged at 500g for 5 min at 4°C. The supernatant was then aspirated and the pellet was resuspended in DPBS. Crosslinking was then performed with 37% formaldehyde for 5 min. Crosslinking was stopped by addition of 2.5M Glycine for 5 min, followed by a wash with DPBS. Nuclei conditioning was carried out with Arima’s Conditioning Solution and Stop Solution 2. Digestion with Enzyme H1 and H2 was then performed for 1h at 37°C, followed by ligation with Enzyme C for 15 min at 22°C, resuspension in 1.5 mL DPBS+BSA, and staining with Hoechst 33342 at a concentration of 1:1000 for 5 min at 4°C. Processed nuclei were brought on ice for sorting into 384-well plates using a Bigfoot Spectral Cell Sorter (Thermo Fisher Scientific, cat. no. PL00299) with a 100 μm nozzle, sheath pressure of 30 PSI, and using 1× PBS sheath fluid. High-efficiency plate sorting was conducted at 4°C using Bigfoot’s “multi-stream” sort deposition, “single” sorting precision, and “Infini-sort” option. Hoechst-positive single nuclei were deposited into individual wells of 16 × 384-well plates (Applied Biosciences ref# 4483285) containing 1ul Digestion Buffer per well.

The snm3C-seq library preparation was done as we previously described^33^. Briefly, library construction was automated on Beckman Biomek i7 platforms for high-throughput production. Nuclei were bisulfite-converted, barcoded with random primers, pooled via SPRI cleanups (compressing 16×384-well plates to 1×96-well), adapter-ligated and PCR-amplified, then cleaned again with SPRI beads. Libraries were quantified by Qubit and normalized. Libraries were initially quality-controlled on an Illumina MiSeq to assess the cis and trans contact ratio and total read counts per cell. QC sequencing was performed using a MiSeq Reagent Micro Kit v2 flow cell with 150-bp paired-end reads. Libraries were subsequently sequenced at greater depth on an Illumina NovaSeq X platform with 156-bp paired-end reads, using either 10B flow cells for 16-plate library pools or 25B flow cells for 32-plate library pools.

### Whole genome sequencing

Genomic DNA was extracted from pulverized frozen tissue using the DNeasy Blood & Tissue Kit (Qiagen). 50ng of DNA was used to prepare the library using NEBNext Ultra II DNA Library Prep Kit for Illumina, fragmenting DNA to 150-350bp for 20min at 37°C. Final libraries were purified with AMPure XP beads and sequenced on an Illumina NovaSeqX 150-bp paired-end reads.

### snm3C-seq data processing

Raw FASTQ files were demultiplexed and aligned to donor-specific reference genomes using the snM3C pipeline (RRID:SCR_025041)^87,108^ with hisat-3n^86^. Donor-specific hg38 reference genomes were constructed by mapping whole-genome sequencing (WGS) FASTQ files to the UCSC hg38 reference, performing variant calling with the GATK pipeline (RRID:SCR_001876) ^96,109^, and incorporating homozygous SNPs into the reference sequence as previously described^33^. The snm3C-seq data for BG regions from the HBA study were downloaded from GEO (accession GSE215353)^33^ and reprocessed using the same workflow.

Following mapping, initial quality control (QC) filtering retained cells satisfying: mCCC fraction < 0.05, mCH fraction < 0.2, mCG fraction > 0.5, and final uniquely aligned reads between 500,000 and 10,000,000. Single-base-resolution methylated cytosine counts and coverage from ALLC files were then aggregated for CG and non-CG (CH) contexts into 100-kb genomic bins for clustering and into gene bodies ±2 kb (gencode v43) for cell-type annotation using ALLCools^87^ (https://github.com/lhqing/ALLCools). For clustering, chrM, chrX, chrY, and blacklist regions^85^ were excluded during the generation of 100kb genomics bins features. To remove the effect of global methylation differences between neuronal and non-neuronal cells and across different cell types, we normalized the raw methylation fraction by dividing by the prior mean methylation level (normalize_per_cell=True in ALLCools), which was estimated by fitting a beta-binomial distribution in each cell^87^.

### Single-cell RNA-seq data processing

Single-cell RNA-seq data generated by 10x Multiome and Paired-Tag^110,111^ were obtained from companion studies^35^ (Chang et al., co-submit). 10x Multiome profiles RNA and ATAC from the same nuclei, whereas Paired-Tag profiles RNA together with histone marks (H3K27ac, H3K9me3, and H3K27me3) in the same cells. Processed and cell-type-annotated scRNA-seq outputs from both platforms were analyzed using a common workflow. To balance cell-type representation and reduce computational burden, we randomly downsampled 1,500 cells per cell type at group levels and performed all downstream analyses on the downsampled dataset. Raw gene counts were aggregated into pseudobulk profiles per cell type, normalized to counts per million (CPM), and log-transformed. As a summary measure of transcriptomic activity for each subclass, we computed the mean gene expression across all cells (sum of gene expression divided by the number of cells).

### ATAC-seq and histone modification data processing

We called narrow peaks for ATAC-seq and H3K27ac and broad peaks for H3K9me3 and H3K27me3 using SnapATAC2^112^. Narrow peaks were merged across samples/cell types to generate a consensus peak set, which was used as input for the ABC model. For pseudobulk signal estimation, we aggregated raw counts per peak (or per fixed genomic bins at 5 kb, 25 kb, and 100 kb) by cell type inferred from the paired RNA profiles, then normalized to CPM and log-transformed (logCPM).

### Integration

To integrate snm3C-seq (methylation) and scRNA-seq (expression) datasets, we first identified the top 200 DMGs and top 200 differentially expressed genes (DEGs) for each cell type or cluster using Scanpy^88^. The union of these gene sets was retained as the shared feature space for integration. We performed integration separately for neuronal and non-neuronal populations. For neuronal cell types, mCH profiles were used for integration, while mCG profiles were used for non-neuronal cell types^32^. Methylation and RNA AnnData objects were log-transformed and scaled independently. The scaled matrices were then concatenated along the feature (gene) axis using the common gene set. To enable label transfer from scRNA-seq to snm3C-seq, a PCA model was trained on the scRNA-seq data and subsequently applied to project the snm3C-seq data into the same PCA space. Integration and cell-type label transfer were performed using the Python implementation of Seurat’s anchor-based integration workflow, as implemented in ALLCools^87^. This method identifies pairwise correspondences (anchors) between cells across modalities and uses these to harmonize the datasets and impute scRNA-seq-derived cell-type labels onto the snm3C-seq cells. The overlap fraction between a methylation-derived cluster and an RNA-derived cell type was calculated as the percentage of cells in the methylation cluster assigned the corresponding RNA cell-type label via the integration-based prediction. The detailed Jupyter notebook for integration is available in the “Data and Code availability” section.

### Clustering of snm3C-seq cells

To cluster snm3C-seq cells using both CG and non-CG (CH) methylation profiles, normalized mCG and mCH fraction matrices (100-kb bins) were separately log-transformed and scaled to unit variance. Principal component analysis (PCA) was then applied to each matrix using Scanpy^88^. Biologically significant principal components were selected using the significant_pc_test function in ALLCools^87^ with a Kolmogorov-Smirnov p-value cutoff of 0.1 between two adjacent PCs.

Preliminary clustering was performed independently on the significant CG and CH PCA embeddings using the Leiden algorithm to define initial cell groupings. Within each preliminary cluster and methylation context, the top 200 cluster-enriched 100-kb bins were selected using the cluster_enriched_features function in ALLCools^87^, which identifies features that are significantly hypomethylated in the target cluster relative to all other clusters. These cluster-enriched features were used to recompute PCA embeddings for CG and CH across the full dataset, followed again by selection of significant PCs via significant_pc_test (p_cutoff=0.1).

The resulting significant PC coordinate matrices from CG and CH were concatenated along the principal component axis to create a unified, multi-context embedding. To mitigate batch effects due to study origin (BICAN vs. HBA), sequencing technology (snm3C-seq vs. snmC-seq), and donor variability, Harmony^89^ was applied to harmonize the joint embedding while preserving biological signal.

Following batch correction, a neighborhood graph was constructed, and final clustering was performed using the Leiden algorithm. Two-dimensional visualization of the integrated clusters was achieved via uniform manifold approximation and projection. Finally, consensus clustering was performed using ALLCools ConsensusClustering, which integrates multiple Leiden clustering iterations to assess cluster stability and applies a supervised random forest-based merging strategy to obtain reproducible, well-separated cell clusters. This approach ensures robust cluster identification by combining unsupervised consensus analysis with supervised refinement based on classification accuracy. Clusters with more than 70% of doublet candidates were considered low-quality cells and excluded. Doublet candidates are the cells with plate-normalized cell coverage (PNCC) >1.2 or <0.8^33^. PNCC was defined as the final mC reads of each cell divided by the average final reads of cells from the same 384-well plate.

To generate a robust group-level dendrogram, we concatenated all normalized 5-kb pseudobulk mCG and mCH profiles from BICAN cells, z-score-scaled the combined matrix, and performed hierarchical clustering in SciPy (RRID:SCR_008058) using average linkage with a correlation-based distance metric^113^.

### Cell type annotation

To enable cross-modality integration and comparison, we aligned the snm3C-seq cells to the HMBA BG consensus cell type taxonomy^35^ using both DEG-based scoring and data integration. We first extracted the top 20 upregulated marker genes for each HMBA cell type at the class, subclass, and group levels from the RNA dataset. For each selected gene set, we calculated a methylation score per cell using the *score_genes* function in Scanpy^88^, which quantifies the relative methylation status of marker gene bodies across cells. Because gene body methylation, particularly mCH in neurons, is generally negatively correlated with gene expression^87,114,115^, lower methylation scores are likely to indicate cells with higher expression of the marker gene set. In parallel, we integrated snm3C-seq and HMBA RNA datasets to transfer cell-type labels from RNA to methylation profiles. Final annotations were determined by jointly considering DEG-based scoring, integration results, canonical cell-type marker and the regional composition of each HMBA cell type. Low-quality cells and cortical glutamatergic neurons (Glut), likely representing contamination, were excluded from all downstream analyses. Detailed clustering and annotation procedures are provided in the Data and Code Availability section.

### Differentially methylated genes and enrichment analysis

To identify DMG, we applied the Mann-Whitney U test to normalized methylation values, followed by Benjamini-Hochberg correction (fdr_bh) for multiple testing. We then computed the median delta methylation difference between the median normalized methylation level in a given cell type and the median normalized methylation level across all other cell types. Additionally, we calculated the area under the receiver operating characteristic (ROC) curve (AUC) to assess each gene’s discriminatory power in distinguishing between cell types. For subclass-level DMGs, we identified DMGs by comparing each subclass with all others, stratified by neuronal and non-neuronal cells. For group-level DMGs, we performed differential methylation analysis between each group and the remaining groups within the same subclass.

We applied filtering criteria to select DMGs for Gene Ontology (GO) and KEGG pathway (RRID:SCR_012773) enrichment analyses, retaining only those with an AUC ≥ 0.8, an adjusted p-value (fdr_bh) ≤ 0.05, and a median delta difference < 0 (for hypomethylated genes) or > 0 (for hypermethylated genes). GO and KEGG enrichment analyses were conducted using the clusterProfiler R package (RRID:SCR_016884)^116^. For Gene Set Enrichment Analysis (GSEA), genes were ranked by the log₂ fold change between the mean normalized methylation in one cell type and the mean across all other cell types, and the ranked list was analyzed using GSEApy^117^.

### Differentially expressed genes

DEGs were identified using the same one-vs-rest framework as for DMGs. High-confidence DEGs were defined by Benjamini-Hochberg FDR ≤ 0.05, fold change ≥ 2, and AUC ≥ 0.8. To select upregulated transcription factors to display alongside TFBS motif enrichments, we used a more permissive definition of upregulated DEGs (fold change ≥ 2 and AUC ≥ 0.7).

### Differentially methylated regions

DMRs were identified using ALLCools^87^ with parameters: max_dist=250, p_value_cutoff=0.001, corr_cutoff=0.3, residual_quantile=0.7, dms_ratio=0.8. DMRs were further filtered to retain only those with an absolute difference in mean methylation fraction (Δβ) ≥ 0.3 and containing ≥ 2 differentially methylated sites (DMSs). Cell type DMRs were defined as regions that show methylation differences (hypo- or hypermethylation) in the corresponding cell type (which may also show hypo- or hypermethylation in other cell types). In contrast, cell-type-specific DMRs were restricted to regions showing significant methylation differences in one cell type, with no significant differences in any other cell type.

### Transcription factor binding sites and motif enrichment

We first extracted DNA sequences from −250 to +250 bp relative to the center of hypo or hyper-DMRs from all cell types, then we identified transcription factor binding motif clusters using Cluster-Buster^118^ (https://github.com/weng-lab/cluster-buster), which scans DMR sequences and detects densely co-occurring motif sites to create the regions_vs_motifs rankings database. Motif enrichment analysis was then performed with pycistarget v1.1^91^ (https://github.com/aertslab/pycistarget), which integrates Cluster-Buster-based motif scores into precomputed cisTarget databases to rank DMRs and calculate normalized enrichment scores (NES) for each motif. Significantly enriched motifs (NES > 3 and Motif_hits >= 10) and annotated TFs were extracted from pycistarget results to infer the putative regulatory network. For visualization, top three enriched motifs and the associated TF (rank by AUC) were displayed for each cell type in Figure 2C.

### 3D Chromatin interaction analysis

The compartment, contact distance, domain, and loop analyses were performed using scHiCluster^119^ (https://github.com/zhoujt1994/scHiCluster). Before analysis, single-cell raw HiC chromatin interaction contacts generated by the YAP pipeline were filtered to exclude blacklisted genomic regions. Following this quality control step, we applied contact imputation at resolutions of 100 kb, 25 kb, and 10 kb using the default scHiCluster parameters. Because single-cell Hi-C data are intrinsically sparse, each cell captures only a small fraction of possible genomic contacts, imputations at higher resolutions were used to denoise and infer missing interactions, thereby improving the robustness and resolution of higher-order chromatin feature detection, including compartments, TAD domains, and chromatin loops.

#### HiC embedding at the single-cell level

To obtain a low-dimensional representation of 3D genome interactions, we performed scHiCluster embedding on the imputed 100-kb contact matrices. The upper triangle of intra-chromosomal contacts within 1 Mb was flattened into a 1D vector for each chromosome. Singular value decomposition (SVD) was applied to each chromosome, retaining the top 50 components, followed by a second round of SVD on the concatenated matrices across chromosomes. The final number of PC components was determined by the significant_pc_test (p_cutoff=0.1) function in ALLCools.

#### Compartments analysis

To identify 3D genomic compartments, we first averaged the imputed contacts at 100kb resolution for each subclass and grouped them into a pseudobulk-level contact matrix. Then, 100kb bins with abnormal coverage were excluded to eliminate low-quality regions. Pearson’s correlation was calculated on the distance-normalized pseudobulk contact matrix, followed by PCA transformation. The first principal component (PC1) was used as the raw compartment score, with its sign oriented so that CpG-rich regions correspond to A compartments. Differential compartments were identified using dcHiC^120^. To enable comparison across cell types, dcHiC normalizes PC1 values using quantiles to standardize score distributions and mitigate batch or scaling effects. The quantile-normalized compartment scores were used in the downstream analysis for cross-cell-type comparisons. To calculate compartment strengths, all 100-kb bins were ranked from smallest to largest by compartment score and divided into 50 equal-sized bins. Distance-normalized interaction frequencies were averaged across all bin-pair combinations of those 50 bins to produce saddle plots, with AA and BB interactions positioned along the diagonal (BB interaction is in the upper left corner). For each cell type, compartment strength was quantified as the ratio of intra-compartment (AA + BB) to inter-compartment (AB + BA) interactions:

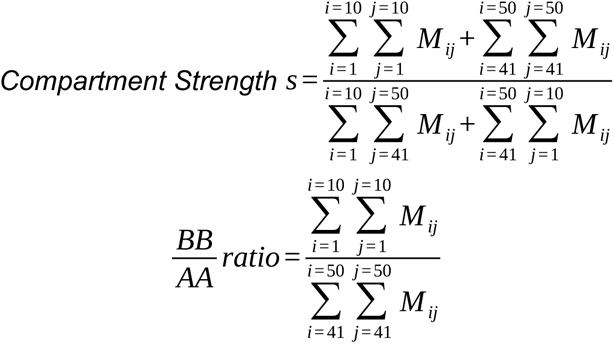

in which *M* is the average interaction frequencies matrix across the ranked 50 *×* 50 equal-size bins, *i* and *j* are the rows and columns coordinates of the matrix *M*.

##### Chromatin contact distance analysis

For each cis contact, both anchors were annotated as A, B, or NA (blacklisted) by intersecting anchor coordinates with A/B compartment using pybedtools^121^. Contact distances were computed with the HiCluster contact-distance module and summarized separately for each compartment-pair category (A-A, B-B, A-B, and NA-associated). For each cell and category, the percentage of contacts in each distance bin was calculated as the number of contacts in that bin divided by the total number of cis contacts in that category. We defined long-range contacts as ≥5 Mb and short-range contacts as <5 Mb. For heatmap visualization of contact-distance distributions, we randomly downsampled 1,000 cells per subclass.

##### Domain

For each genomic bin (at the given resolution), we computed the insulation score by looking at contacts within a window of size on either side, and measuring how ‘insulated’ that bin is from its neighbours. A low insulation score indicates a strong boundary (few contacts cross the bin), while a higher score indicates a weaker boundary (many contacts cross the bin, domain merging, or a less constrained architecture).

##### Differential domain boundaries

Per-cell domains and 25-kb insulation profiles were computed with scHiCluster’s domain module on imputed contact matrices, and per-Subclass boundary occupancy was estimated as the fraction of cells with a called boundary at each bin (bound_count_ct / n_cells). At each bin, between-Subclass differences in occupancy were tested by a chi-squared test on the (boundary, non-boundary) × subclass contingency table and corrected by Benjamini-Hochberg FDR at 0.01. A bin was retained as a differential boundary if it (i) was outside the 10 kb blacklist and ≥ 250 kb from chromosome ends, (ii) was a local minimum of at least one subclass’s insulation track (scipy.signal.find_peaks on −insulation, distance/width = 5; prominence used as boundary strength), (iii) was a local maximum of the chi² statistic (height = chi² at FDR < 10⁻³, distance = 5) with a normalized chi² z-score > norm.isf(0.025), and (iv) showed a between-Subclass boundary-probability range > 0.05. Each retained boundary was assigned to the subclass with the highest occupancy, intersected with gene bodies (TSS ± 2 kb) using bedtools, and cross-referenced with subclass DMGs and DEGs (HMBA RNA, Paired-Tag RNA).

##### Enrichment of convergent multi-omics genes at differential boundaries

For each of the 2,110 differential boundaries, we asked whether any overlapping gene (TSS ± 2 kb) was simultaneously a hypo-DMG, an HMBA snRNA-seq up-DEG and a Paired-Tag RNA-seq up-DEG (both: FC ≥ 1.5, auc ≥ 0.7, FDR ≤ 0.05) in the same subclass. Significance was assessed by Monte Carlo permutation (10,000 iterations, numpy.random, seed 42) under a hypergeometric null: each boundary in subclass G with n genes was modeled as an independent Bernoulli trial with success probability 1 − hypergeom.pmf(0, N, k_G, n) (N = 37,950 DMG-tested genes; k_G = convergent target genes in G), preserving boundary count and subclass identity. Enrichment was reported as fold change over the null mean, with a z-score and a one-sided permutation p-value.

##### Loop

Loops and differential loops were identified as we previously described^33,122^. Briefly, chromatin loops were called with scHiCluster from imputed single-cell contact maps, restricting candidate interactions to 50 kb-5 Mb. At a fixed bin resolution (10-kb), each loop corresponds to an enriched bin-bin interaction (a “pixel”) in the contact matrix. For each cell, the imputed matrix (Q) was log-transformed and diagonal-wise Z-scored to obtain a global-normalized matrix (E), and a local-background-corrected matrix (T) was computed by subtracting the 30-50 kb neighborhood background. Within each cell group, pixel-wise enrichment against global (E) and local (T) backgrounds was quantified using pseudo-bulk t statistics, with an empirical FDR estimated from diagonal-shuffled controls and additional local fold-change filters to define significant loop pixels.

##### Differential loops

For every subclass we accumulated cell-level first (Σ E) and second (Σ E²) moments of the Q and T matrices into pseudo-bulk *.Q.cool / *.Q2.cool and *.T.cool / *.T2.cool files, then computed a pixel-wise ANOVA F-statistic across subclasses (between- vs within-Subclass variance) after masking blacklisted 10 kb bins and restricting to 60 kb-5 Mb separations. Loop pixels detected in at least one subclass were retained as differential when both the Q- and T-derived F z-scores exceeded the 85th percentile (z > norm.isf(0.15) ≈ 1.04) and both anchors lay 100 kb-4.95 Mb apart and ≥ 100 kb from chromosome ends. A subclass-specific loop set was then defined per subclass c as the differential loops whose mean Q in c exceeded 1.2× the mean Q in all other subclasses and whose mean T exceeded 1.5× the mean T in the rest.

##### Aggregate peak analysis (APA)

For every subclass c the imputed Q matrix was converted to an observed-over-expected map by dividing each off-diagonal pixel by the diagonal mean (decay[k] = mean(diag(Q, k)), k = 1, …, 504). For every subclass-specific loop set s, each loop centre was extracted as a 21 × 21 ±100 kb sub-window, min-max-normalized to [0, 1] and averaged across loops, yielding an APA map result[c, s]; iterating over all (c, s) pairs produced the (N, N, 21, 21) APA tensor.

##### Cross-Subclass APA strength matrix

Each result[c, s] panel was summarized by an APA strength score = centre pixel / mean of the lower-left 5 × 5 corner, giving an N × N matrix (rows = signal subclass providing the Q matrix; columns = loop-set subclass providing the diff-loop centres). To align lineage blocks along the diagonal, the matrix was symmetrised and a shared row/column ordering was obtained by average-linkage clustering of the squareform distance (scipy.cluster.hierarchy.linkage(…, method=’average’)); the reordered matrix was rendered with PyComplexHeatmap.

### Cross-modality comparison

To relate DNA methylation to transcription, we performed one-vs-rest comparisons for each subclass (within neuronal and non-neuronal lineages) and stratified genes into four sets: DMG-only (CG or CH hypomethylation; mean normalized methylation < 1, median Δmethylation ≤ −0.2, AUC ≥ 0.8; not upregulated), DEG-only (upregulated; BH FDR ≤ 0.05, fold change ≥ 2, AUC ≥ 0.8; not hypomethylated), DEG&DMG (meeting both criteria), and a background set sampled from genes lacking evidence for differential methylation (p-value ≤ 0.05 and mean normalized methylation < 1) and differential expression (p-value ≤ 0.05 and fold change > 1.5). Those differential genes were between one subclass versus the rest of the subclasses in neuronal and non-neuronal cell types. The background set was downsampled to match the largest of the DMG-only, DEG-only, and DEG&DMG groups, ensuring comparable gene-set sizes. Metagene methylation profiles over peaks, DMRs, and TSS/gene bodies were computed with deepTools (RRID:SCR_016366, computeMatrix)^93^. For TSS/gene-body profiles, results were additionally aggregated into neuronal and non-neuronal summaries and visualized using custom Python scripts. The enrichment of histone modifications, domain boundaries, ATAC and DMR on A/B compartments was calculated using bedtools (RRID:SCR_006646, module *fisher*)^123^.

### Activity-by-Contact model

We applied the Activity-by-Contact (ABC) model^90,124^ (https://github.com/broadinstitute/ABC-Enhancer-Gene-Prediction) to predict cell-type-specific enhancer-promoter interactions using pseudobulk profiles of chromatin accessibility peaks and 3D contact frequency as primary inputs. Briefly, the pipeline first identified candidate regulatory regions by retaining the top 150,000 strongest ATAC-seq peaks (centered and resized to 500 bp, excluding blacklisted regions). Hi-C contact matrices were normalized using power-law scaling to adjust for distance-dependent decay. For each gene, the model computed an ABC score for candidate enhancers within a 5 Mb window, combining enhancer activity and scaled contact frequency with the gene promoter. The ABC score threshold of 0.021 was selected in accordance with the official ABC guidance. Predicted enhancer-promoter links were retained only if the target gene showed evidence of cell-type-specific regulation in at least one of the following criteria (all at FDR ≤ 0.05): (1) Upregulation (fold change ≥ 1.5, and auc_score ≥ 0.7); (2) Hypomethylation (mean normalized methylation fraction < 1, auc_score ≥ 0.7, and median delta difference <= −0.2); and (3) Increased chromatin accessibility (fold change ≥ 1.5, and auc_score ≥ 0.7).

### LDSC

We obtained GWAS summary statistics for Alzheimer’s Disease and Dementia^125^, Parkinson’s Disease^126^, Schizophrenia^127^, Major Depressive Disorder^127^, Attention Deficit Hyperactivity Disorder^128^, Bipolar Disorder^129^, Autism Spectrum Disease^130^, Insomnia^131^, Obsessive Compulsive Disorder^132^, tiredness^133^, Multiple Sclerosis^134^, Neuroticism^135^, educational attainment^136^, problematic opioid use^137^, intelligence^138^, Amyotrophic Lateral Sclerosis^139^, age at first birth^140^, Type 1 Diabetes^141^, and Anorexia Nervosa^142^. We converted the summary statistics into the standard format for linkage disequilibrium score regression and annotated the input regions with the 1000 Genomes Project Phase 3 SNPs in hg38 (https://doi.org/10.5281/zenodo.7768714). We ran cell-type-specific analyses to perform regression for each trait (https://github.com/bulik/ldsc). The union of regions across all cell types for a given annotation type was used as the background. In the case of a small annotation size (i.e., coverage < 1% of the genome, approximately 90,000 SNPs), LDSC can yield elevated false-positive results and should be interpreted with caution^143^.

### MERFISH

#### Gene panel design

A 920 gene panel (Table S11) was designed. The initial release of the HMBA BG taxonomy was used as the reference taxonomy, as well as the clustered snm3C-seq data. For the HMBA snRNA-seq dataset, nsforest^98^ was used to generate cell-type marker genes. On the snm3C-seq data Spapros^97^ was applied at the subclass, group, and brain region level. The gene lists were combined, and top 3 DEGs and top 3 DMGs for each cell-type were added. Additional canonical cell-type markers were chosen based on literature. 20 internal control genes, chosen by looking at similar expression levels across all cell-types in the Siletti et al.^36^ whole brain dataset, were also included in the panel for quality assurance purposes. Genes flagged by the Vizgen gene panel design tool as either being too short, too highly expressed, or not having enough probe targets were removed. The gene panel was evaluated on the initial HMBA release for its ability to recapitulate the identified cell-type groups.

#### MERFISH sample preparation

Samples were prepared for imaging in accordance with Vizgen’s MERFISH 2.0 Sample Preparation User Guide for Sectioned Tissue Samples, available on the Vizgen website at https://vizgen.com/resources/user-guides/. This protocol was performed according to MERFISH 2.0 chemistry following the guidelines and volumes for large slides. The reagents used in this protocol were from the MERSCOPE Tissue Sample Prep Kit, V 2.0, Box 1 (Vizgen, 10400194) and Box 2 (Vizgen, 10400194). Briefly, the fixed tissue sections did not undergo cell-boundary staining; instead, they were pretreated with anchoring. After permeabilization and photobleaching, 70% EtOH was aspirated from the samples and a conditioning buffer was added. Next, a pre-anchoring activator was added directly to the tissue sections and covered with a 2 cm^2^ section of parafilm, and kept at room temperature for a minimum of 16 hours. On day two, an anchoring buffer was added to the samples and incubated at 37°C for 2 hours in a humidified incubator. The tissue samples were then embedded in gel and kept at room temperature for 1.5 hours to set. Once the sections were gel-embedded, samples were cleared using a 1:100 mix of Proteinase K (NEB, P8111) and clearing solution, and incubated for 24-48h at 37 °C. To optimize the protocol for human tissue, the samples were incubated at 37°C rather than 47°C, as specified for this incubation step. The clearing was validated by ensuring the tissue was no longer visible in the gel after incubation. For the encoding probe hybridization on day three, a 1:20 mix of MERSCOPE 1000 Gene Panel, V 2.0 (specific to the basal ganglia) (gene panel ID: D2P2238) and Probe Dilution Buffer (Vizgen, 10400181) was added to hybridize directly to the tissue and incubated at 47°C for 20-24 hours. After incubation, Enhancer Probe Mix was added to the tissue and placed in the 37°C incubator for 18-24 hours, then subsequently washed before proceeding to imaging. Extra care was taken to ensure samples were prepared in an RNase-free environment, with the work area and tools cleaned with 70% EtOH and RNase Zap (Thermo Fisher, AM9782). For each incubation exceeding 30 minutes, the 60 mm petri dishes containing the MERSCOPE slides were sealed with parafilm and sealed inside a 150 mm dish with 1-2 Kimwipes and 5-10 mL of nuclease-free water (Thermo Fisher, SH3053803), creating a humidified chamber.

#### MERSCOPE imaging

The MERSCOPE imaging was performed using MERSCOPE Ultra (VMSC31910 and VMSC31810) with all imaging done in accordance with Vizgen’s MERSCOPE UltraTM Instrument User Guide, which can be found at https://vizgen.com/resources/user-guides/. In brief, the MERSCOPE Large 1000 Gene Imaging Kit V 2.0 (Vizgen, 10400174) was thawed in a 37°C water bath for 1 hour, activated with the imaging buffer activator and murine RNase inhibitor (NEB, M0314L), and then placed into the MERSCOPE. Sections were stained with DAPI and PolyT and incubated at room temperature for 15 minutes, covered from light. Samples were then fixed with formamide, washed with a wash buffer, and assembled into the large MERSCOPE flow chambers. The fluidics were primed, and the flow chambers were filled with liquid, ensuring no bubbles. The MERSCOPE instruments generated low-resolution mosaics at x10 magnification (NA=0.25). Four separate regions of interest (ROI’s) were manually defined, one for each donor, with a maximum area of 3.0 cm^2^. The instruments switched to the high-magnification objective lens, imaging the defined ROI’s with the smallest XY field of view (FOV) size of 280 µm^2^ at x40 magnification (NA=1.40). For the basal ganglia, six different markers were imaged: MBP, ACTB, GFAP, UBC, GAPDH, and MAP2. Imaging of the readout probes was performed at 750 nm, 650 nm, and 560 nm, and images were acquired in the 405 nm and 488 nm channels for DAPI and PolyT. To image the tissue with a 10 µm imaging depth, 7 z-stacks of 1.5 µm thickness were acquired. MERSCOPE imaging of the large slides took on average 24-48 hours. After imaging, image analysis was performed using Vigen’s analysis tool software version 234b.241217.1593.

#### Image preprocessing

Image processing was performed using the in-house SPIDA toolbox and mimicked the analysis from a previously published paper^144^, with adjustments to better fit our dataset. The MERSCOPE Ultra returned single Z-slice images for DAPI and PolyT; these were duplicated to 7 Z-slices and then deconvolved individually using Deconwolf^99^ with a generated PSF matching the imaging wavelength (DAPI = 405nm, PolyT = 488nm). The outermost Z-slices were then discarded, and the remaining 5 Z-slices were averaged to create a 2D high-contrast image. The images were downsampled to 4x resolution to fit in memory, then segmented using the CellposeSAM^101^ model with permissive filtering parameters. The cellpose segmentations were regarded as the ‘ground truth’ cell nuclei segmentation. After the initial cellpose segmentation, Proseg^102^ was applied to the detected transcripts using the cellpose seeds, resulting in expanded geometries. The resulting Proseg segmentations were then aligned with the initial cellpose segmentations, with Proseg cells that did not correlate with any cellpose cells being discarded, and all cellpose cells that did not match Proseg cells being kept for further analysis. Finally, the joined cellpose and proseg geometries were used to generate cell-by-gene and cell-metadata tables for downstream analysis using vizgen’s postprocessing toolkit (https://github.com/Vizgen/vizgen-postprocessing).

#### Spatial data preprocessing

The data was stored and manipulated within the SpatialData ecosystem^100^. The segmented cells were QC’d and filtered based on the following values. Total RNA count > 20 genes, number of unique genes per cell > 5, volume > 100 um^2^, and blank count < 5. Following the first round of QC, the bottom 2 percentiles of cells by gene density (RNA count/volume) were removed, as these were considered lower-quality cells. SOLO doublet detection was performed at this stage; upon manual inspection, all cells we checked which were identified as doublets, clearly corresponded to a single nucleus DAPI stain. Therefore, no doublet detection was utilized. The data was first volume-normalized by dividing all gene counts by the cell’s volume to control for artifacts relating to cell size. Then, each cell’s total number of volume-normalized genes was again normalized to the median library count size (per sample). Finally, log1p was applied to the volume-median-normalized counts for numerical stability. These counts were the ones used for all downstream analyses unless otherwise stated.

#### RNA integration and annotation

To annotate the MERFISH data, a procedure similar to that above, using the ALLCools Python implementation of CCA, was employed with several modifications. The reference HMBA scRNA-seq-derived taxonomy was regarded as the ground truth. Each individual MERFISH sample (brain region × donor × replicate) was matched to a subset of the reference dataset for the same brain region. Both datasets were independently normalized, and the MERFISH dataset was projected onto the same PCA space as the reference dataset, and CCA was applied. The k-nearest neighbors (kNN) graph was computed in the joint PCA space, and anchors were identified across datasets. Co-clustering of the same dataset was performed. Additionally, each MERFISH cell was given a label transfer score based on the weighted distance in the KNN graph between it and the 30 nearest reference cells. MERFISH cells whose co-clusters were composed of > 60% of a given reference cell-type were annotated as such, and cells with poor label transfer scores were labeled as ‘unknown’. Each sample was first annotated as “Neuron” or “Nonneuron,” and these types were later integrated separately to remove potential technical artifacts arising from gene panel design and differences in detection efficiency between white matter and gray matter. The samples were first labeled at the subclass level. Later, each individual subclass was re-integrated at the subclass level. The distribution of label transfer scores within each subclass and group was termed the ‘annotation score’, and used to assess the efficacy of the assigned annotation for each cell type.

#### Methylation integration and imputation

Each MERFISH sample was integrated with its region-matched but not donor-matched corresponding data block in the snm3C-seq dataset. The reason samples were not donor-matched is due to limitations in the amount of available tissue per donor, as MERFISH blocks were taken from adjacent dissection blocks used for sequencing, which, at times, did not directly overlap in brain substructure composition. Integration and imputation were done using the ALLCools package. First, each dataset was individually normalized, and genes were subset to those showing some variance in the MERFISH sample (std. dev. > 0.01), and then further to those enriched in a cell type in the snm3C-seq dataset (using ALLCools cluster_enriched_features function). PCA was run on the snm3C-seq data, and the MERFISH sample was projected onto that PCA space.

Similar to RNA integration, methylation integration was performed in the shared PCA space using the CCA implementation in ALLCools. Label transfer was performed using the KNN graph. The snm3C-seq data and the MERFISH sample were co-embedded post-integration to generate a joint Leiden clustering, which was used to perform annotation in a manner similar to the RNA annotation. Methylation information was imputed only for cells with matching RNA and methylation cell-type annotations. Imputation was done by assigning each methylation cell the spatial location of the nearest MERFISH cell in the shared KNN graph returned by the CCA integration. The label transfer score for the methylation type matching the MERFISH cell-type was calculated, and termed the ‘annotation score’ for the methylation cells due to its conceptual similarity to the RNA annotation score. The annotation score was used to assess the quality of the imputation of methylation cells onto the MERFISH cells.

#### White matter region calling

To avoid biases in downstream analyses, we developed a method for calling and removing white-matter tracts from analyses involving spatial localization. We reasoned that BCAS1, which is a marker of myelinating Oligodendrocytes, would exhibit a bimodal distribution of expression patterns. The first peak will correspond to gray matter, which has some myelin, and the second peak will correspond to white matter, which is enriched in myelin. We generated a hexagonal grid to lay over the spatial locations of all detected genes in a given sample; each hexagon in the grid had an internal radius of 40 um, and there was no overlap between hexagons. We then counted the number of BCAS1 genes detected per hexagon, and fit a Gaussian mixture model using sklearn’s GaussianMixture (sklearn.mixture.GaussianMixture)^145^ functionality on the hexagons. This allowed us to classify each hexagon as either coming from the lower distribution (supposedly gray matter), or from the larger distribution (supposedly white matter). This over-parametrized model worked well in cases where both white matter and gray matter were present in the same sample, but struggled in samples with only one type of region (e.g., the GP).

#### Matrix-Striosome compartment calling

To call matrix and striosome compartments, we used the SPN-specific annotations. For each SPN cell in our dataset, we generated a vector of counts of all other SPN neighbors (at the group level) within a 200um radius. We used the K-Means clustering implemented in sklearn (sklearn.cluster.KMeans) to cluster all cells based on their neighborhood vectors. K was set to 3 to account for both the dorsal-ventral and matrix-striosome axes. We manually determined which cluster corresponded to the striosome and which clusters corresponded to the Matrix, and labeled them accordingly. We constructed another nearest neighbor graph (r=150um), this time stratified by whether a cell was identified as matrix or striosome, and removed all connected components with less than 5 cells in them. We used Pysal (libpysal.weights.DistanceBand) to generate this second graph (https://pysal.org), and networkx’s connected_components function (RRID:SCR_016864) to find all connected components (https://github.com/networkx/networkx). For each connected component, we then calculated the convex hull using alphashape (https://alphashape.readthedocs.io). Since the convex hull of the matrix often included the smaller striosome region, we removed the intersection of the matrix and striosome compartments using a 50 μm buffer. Finally, to integrate the white matter with the Matrix-striosome compartments, whenever there was any intersection between the two compartment types we assigned the intersecting region to the annotation of the shape with the smaller area, reasoning that the larger area probably encompassed the smaller area due to limitations in the calculation of the convex hull to correctly capture holes.

#### Compartment enrichment analysis

To perform compartment enrichment analysis an assumption was made that each MERFISH section can be treated as an i.i.d. sample from a distribution of cell-type enrichments in either compartment. This assumption, while not strictly true, enables the use of standard statistical framework to assign significance to qualitative enrichments. Due to this assumption, care is needed when interpreting the results, and stringent p-values cutoffs are used to assign significance. For this test we define *S* as the set of all independent MERFISH sections which have matrix and striosome compartments:

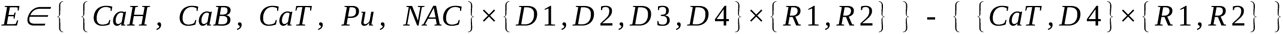

Where *D1-4* are the 4 donors, *R1-2* are the two replicates, and the brain regions are those in the striatum (one of the donors did not have a CaT sample).

Let *T* be the set of all cells, *T_i_* be the set of cells of type *i* in the entire dataset, *T_e_* be the set of cells in sample *e ∈ E*., and *T_e,i_* be the set of cells of type *i* in sample *e*. Let *M, M_i_, M_e_, M_e,i_* be the analogous sets of cells in matrix compartments, and *S, S_i_, S_e_, S_e,i_* be the analogous set of cells in the striosome compartments.

To ensure that samples with small relative proportions of a given cell-types do not bias the analysis, cell types in individual samples are filtered using two parameters. Define *R_e,i_*as the ratio of the ratio of cells of type *i* in sample *e* to the total number of cells in sample *e*, to the ratio of cells of type *i* to the total number of cells in the entire dataset.

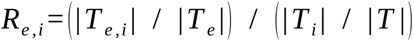

If *T_e,i_* < 5 or *R_e,i_* < 10 %, cell type *i* from sample *e* is removed.

The raw counts are normalized by the cell-type specific and compartment specific counts. Defined 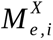 and 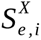 as the normalized counts of cell type *i* in sample *e* in the matrix and striosome, respectively, as follows:

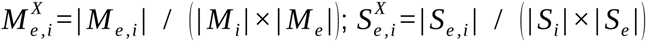

The log2 fold change is then taken between the matrix and the striosome normalized counts for each sample and cell type.

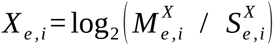

Define *X_i_* as the distribution of fold changes for each cell type { *X_e,i_* | *e ∈ E* } for which cell type *i* was not filtered from *e*. Outliers were removed from *X_i_* using a standard IQR metric (values omitted if they were more than 1.5 *× IQR* below the 25th or above the 75th quantile. The remaining values in *X_i_* were visualized and showed approximate normal distributions. At this stage cell types with less than 10 values in *X_i_* were removed to avoid small sample sizes that can significantly bias the results. A one sample t-test was performed using scipy (RRID:SCR_008058) (scipy.stats.ttest_1samp) against a population mean of 0. The resulting p-values were corrected using the Benjamini-Hochberg method implemented in statsmodels (RRID:SCR_016074) (statsmodels.stats.multitest.multipletests, method = “fdr_bh”). The FDR adjusted p-values were used for statistical analysis, the mean of *X_i_* was used to quantify effect size, and a 95th percentile confidence interval was computed using scipy for visualization.

For the enrichment analysis on genes, the same approach was used, but *i* referred to genes instead of cells.

#### Calculating compartment-enriched genes

Differential gene expression analysis between the matrix and the striosome compartments was performed using a pseudobulk DESeq2 (RRID:SCR_015687) approach implemented with PyDESeq2^103^. Matrix-striosome differentially expressed genes (MS-DEGs) were called independently for each brain region (CaH, CaB, CaT, Pu, NAC) at both the subclass and subclass levels. For each brain region - cell type - compartment combination, raw RNA counts were summed across all cells belonging to the same donor and technical replicate. This generated 8 potential pseudobulked samples in either compartment for each brain region - cell type combination for the CaH, CaB, Pu, and NAC brain regions, and 6 such pseudobulked samples for the CaT, due to the missing donor. A brain region - cell type combination was omitted from the DESeq2 analysis if one of the compartments had less than 3 pseudobulked samples.

The standard DESeq2 pipeline was run on the pseudobulked counts with the design formula: “∼ donor + group”. Donor was included to account for inter-individual variability and the group variable corresponded to the compartment (matrix vs. striosome). Cook’s distance outlier refitting was enabled, and the log2 fold-change estimates were shrunk (refit_cooks=True, lfc_shrink=True). Significance was determined on a per brain region basis using a Benjamini-Hochberg adjusted p-value cutoff of p < 0.001, and a shrunken a log2 fold change cutoff of | Log2FC| > 0.5. The stringent p-value cutoff was used due to the limited power of having only 4 donors. To further increase the reliability of the results, MS-DEGs were included in the final results only if they were significant, and directionally consistent in at least 2 brain regions.

To exclude potentially missegmented genes from being included in the downstream analysis, a filter was constructed using the separate HMBA snRNA-seq data. A gene was included as an MS-DEG in a given cell type only if it was expressed in more than 10% of cells of the same cell type in the HMBA data.

#### Imputing Compartment Status of snm3C and HMBA cells

The compartment localization of individual STR FS PTHLH-PVALB GABA snm3C cells was determined by mapping backwards from the same population of MERFISH cells, using the previously derived snm3C to MERFISH map. Since an snm3C cell can map to multiple MERFISH locations across the samples, each cell was assigned a striosome score, which was defined as the ratio of striosome labeled MERFISH cells to total MERFISH cells a given snm3C mapped to. snm3C cells were omitted if they mapped to less than 3 MERFISH cells. A cell was determined as being located in the striosome if its striosome score was greater than 20%, and it was deemed located in the matrix otherwise. The 20% cutoff was chosen qualitatively based on a split in a bimodal distribution of striosome scores. The compartment label was termed MS_compartment.

HMBA STR FS PTHLH-PVALB GABA cells were mapped in a similar fashion to the snm3C cells. An imputation graph was calculated in the same manner as the snm3C cells, through a new round of annotation that limited cell imputation to the same group level in the taxonomy. From there the same procedure was used as above, with the exception that the striosome score cutoff for HMBA cells was 25% due to a qualitative split in the bimodal distribution of striosome scores.

The snm3C STR FS PTHLH-PVALB GABA was clustered as described above, and a mapping between the de novo clusters and the striosome and matrix compartment was computed. The STR FS PTHLH-PVALB GABA HMBA dataset had precomputed Cluster labels, which were also mapped onto striosome and matrix compartments.

Integration between the snm3C and HMBA STR FS PTHLH-PVALB GABA specific datasets was done like the integration for the entire datasets. To create consistent labels across modalities, joint clusters were mapped onto matrix or striosome compartment labels via MS_compartment proportions in both datasets.

#### Integration and compartment imputation of Patch-Seq cells with snm3C and HMBA cells

Two integrations were performed with the STR FS PTHLH-PVALB GABA Patch-seq dataset. Since the gene expression matrix of the Patch-seq dataset came from macaque, first the intersection of all genes was taken between the datasets, and only those genes were kept for the integration. snm3C x Patch-seq integration was performed as above (snm3C-HMBA integration), with the snm3C cells serving as the reference. The consistent MS_compartment labels were transferred to the Patch-seq cells. HMBA x Patch-seq integration was performed as above (MERFISH-HMBA integration), with the HMBA cells serving as the reference. The consistent MS_compartment labels were transferred to the Patch-seq data. Only Patch-seq cells whose MS_compartment annotations were consistent between both integrations were kept for downstream analysis.

#### Quantifying electrophysiological differences

Electrophysiological features were generated in a companion paper using the Intrinsic Physiology Feature Extractor (RRID:SCR_028075) tool from the Allen Institute. Only features with at least 5 cells per compartment were included in the analysis. Effect size was calculated using the rank-biserial correlation:

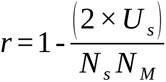

Where U_s_ is the Mann-Whitney U statistic, and N_s_, and N_M_ are the number of striosome and matrix cells, respectively. Significance testing was done using the Mann-Whitney U test as implemented in scipy (scipy.stats.mannwhitneyu), and p-values were corrected using Benjamini-Hochberg as implemented in statsmodels (statsmodels.stats.multitest.multipletests). And FDR cutoff of 0.1 was used. Low n numbers meant the test was underpowered and features did not pass strict FDR cutoffs, so differences were ranked by effect size.

### Data visualization

Heatmaps were generated with PyComplexHeatmap^95^, and multi-omic locus track visualizations were created using pyGenomeTracks^94^ and are available through SCMDAP (https://neomorph.salk.edu/SCMDAP/BasalGanglia). Circos plot was generated using pycirclize (https://github.com/moshi4/pyCirclize), and cell-type-specific gene regulatory networks were visualized using Cytoscape^146^.

## Data and code availability

Raw fastq files are available via the NEMO platform at the landing page: https://assets.nemoarchive.org/dat-aw6czix, cell-level fastqs and processed files (pseudobulk level allc and single-cell level 3C files) are available at GEO (accession GSE320293). MERFISH data are available on BIL (https://doi.org/10.35077/g.1194). All processed data and analysis code are available at https://github.com/DingWB/BG_snm3C-seq. Cell type- and region-level methylation profiles can be queried and visualized at http://neomorph.salk.edu/hbg/hbg.php, multi-omic data visualization portal of basal ganglia is accessible at https://basalganglia.epigenomes.net.

## Acknowledgments

This publication was supported by and coordinated through the Brain Initiative Cell Atlas Network (BICAN, RRID:SCR_022794). This work used Anvil^147^ HPC cluster at Purdue University through allocation MCB130189 from the Advanced Cyberinfrastructure Coordination Ecosystem: Services & Support (ACCESS) program, which is supported by National Science Foundation grants #2138259, #2138286, #2138307, #2137603, and #2138296. This work was supported by the Flow Cytometry Core Facility of the Salk Institute (RRID:SCR_014839) with funding from NIH-NCI CCSG: P30 CA014195, and Shared Instrumentation Grants S10-OD023689 (Aria Fusion cell sorter), and S10 OD034268 (Thermo Fisher Bigfoot). We gratefully acknowledge the Broad Institute and NEMO for conducting the sequencing and data storage. We also thank Wenhao Cao from UC Irvine for valuable discussions. **Funding**. National Institute of Mental Health, UM1MH130994, R.W. was supported by the T32 GM145427 training grant.

## Author contributions

J.R.E. and M.M.B. supervised the study. C.T.B.-B., J.A.R., A.B., R.G.C., J.R.N., E.O., W.O., A.P., C.C., A.S.A., A.S.B., J.L., K.G.R., K.W.K., C.K.Y., J.K.W., C.B., J.A., S.C., J.A., D.C., E.S., J.L., M.J., S.V., G.V.S., A.C.M., Y.S., A.B., C.O., M.L., M.V.M., C.R., S.N.A., J.B., M.H., Z.Z., J.F., C.T., J.O., Q.Z., X.X., E.D., Z.Z., J.F., C.T., and J.O. generated the data, Y.F., Y.X., K.L., L.C., N.J., X.L., B.K., R.D.H., T.E.B., E.S.L., B.R., shared the multiomic datasets, W.D., A.K., R.W., Q.Z., K.K., S.F., W.T., and Y.W., C.K., E.B., J.K., and E.B. analyzed the data. W.D., A.K., R.W., M.M.B., and J.R.E. interpreted the data. W.D., A.K., C.T.B.-B., J.A.R., E.O., R.W., M.M.B., and J.R.E. wrote the manuscript. N.S.A., W.Z., H.C., D.L., and T.W., coordinated and helped with the data uploading or visualization. All authors edited and approved the manuscript.

## Supplementary figures

**Figure S1.**
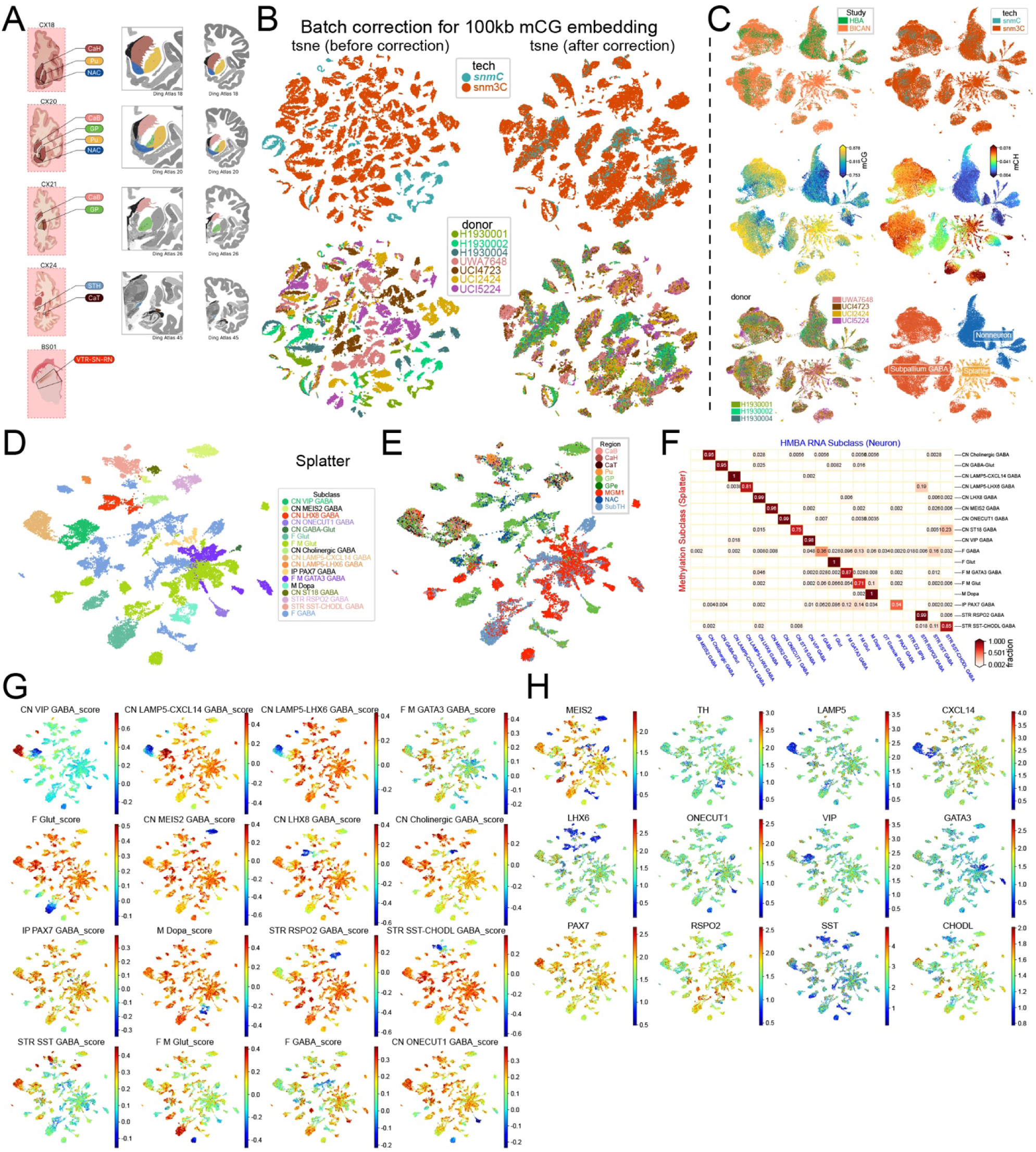
Brain dissection, UMAP embeddings, and validation of cell-type annotation transfer for splatter subtypes. (A) Anatomical criteria for brain dissection (B) UMAP embedding colored by technology and donor before and after batch correction. (C) UMAP embeddings colored by study of origin, technology, global mCG and mCH fraction, donor, and supertype. (D-H) Annotation and validation workflow for splatter. (D) UMAP of splatter cells colored by subclass. (E) UMAP colored by the regional composition of splatter cells. (F) Confusion matrix comparing RNA-derived cell-type labels with methylation cell type annotation in the shared integration space. (G) UMAP colored by hypomethylation scores computed from RNA-derived top marker genes for each cell type. (H) UMAP colored by mCH levels at canonical marker genes used for cell-type identification.

**Figure S2.**
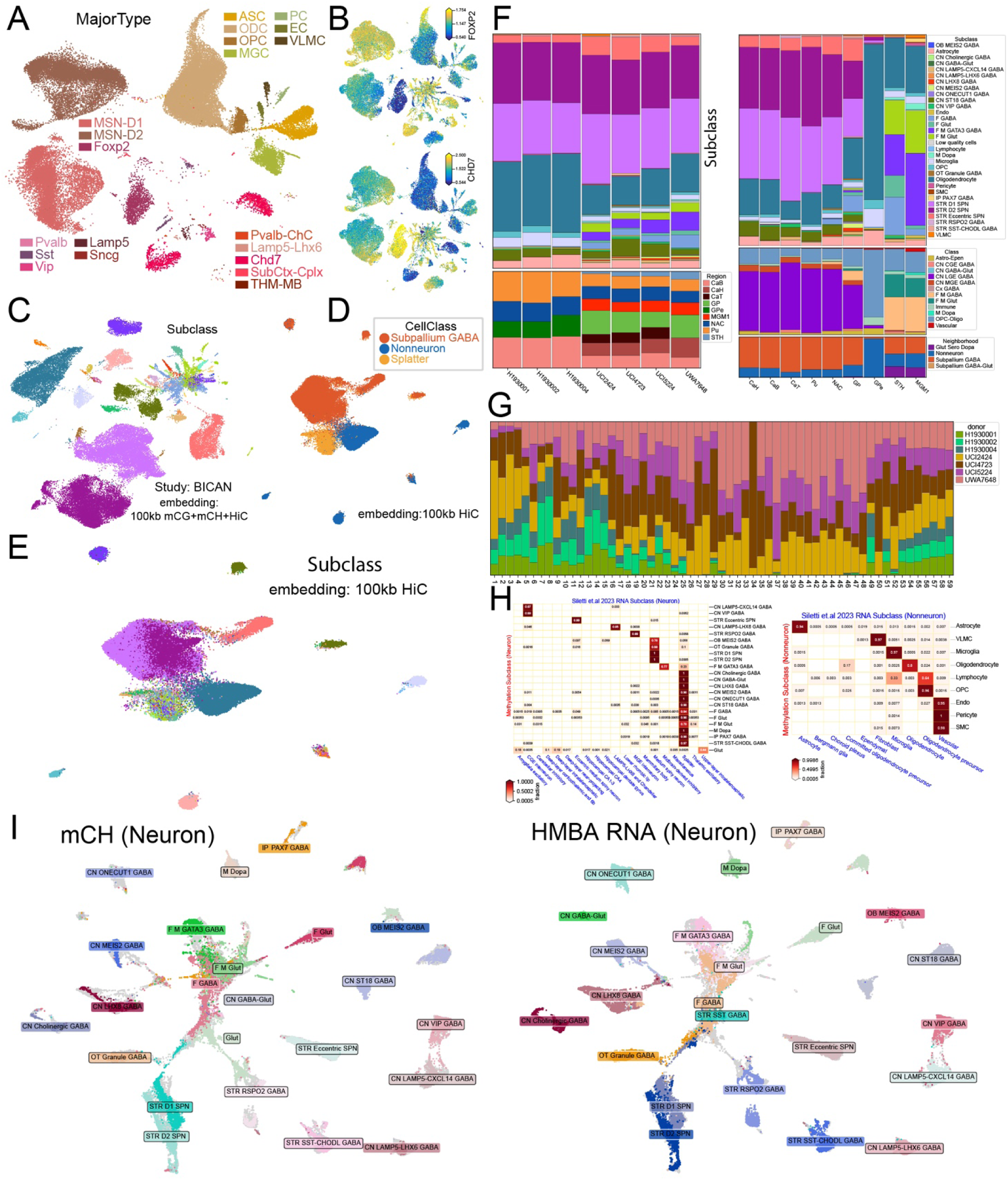
Dataset-specific embeddings, marker hypomethylation examples, and region/donor composition across integrated datasets. (A) UMAP embedding of cells from the Human Brain Atlas (HBA) study, colored by MajorType annotation. (B) UMAP showing gene-body CH hypomethylation at the *FOXP2* locus in STR eccentric SPN and CHD7 locus in striatal ST18 cells (CN ST18). (C) UMAP embeddings of BICAN cells based on 100-kb resolution mCG, mCH, and 3D chromatin interaction features, colored by brain region and subclass annotation. (D) Clustering of BICAN cells using 100-kb 3D chromatin interaction (HiC) embeddings, colored by methylation-derived CellClass labels. (E) The same embeddings as (D), but colored by subclass. (F) Cell-type composition across basal ganglia regions, and region/subclass composition across individual donors. (G) Donor composition across group-level cell types. (H) Confusion matrices comparing methylation-derived cell-type annotations with RNA-derived cell-type labels (analyzed separately for neuronal and non-neuronal cells) at the subclass level in the Siletti et al. 2023 RNA dataset. (I) UMAP embeddings that are colored by methylation cell types and HMBA RNA cell types in the same integration space.

**Figure S3.**
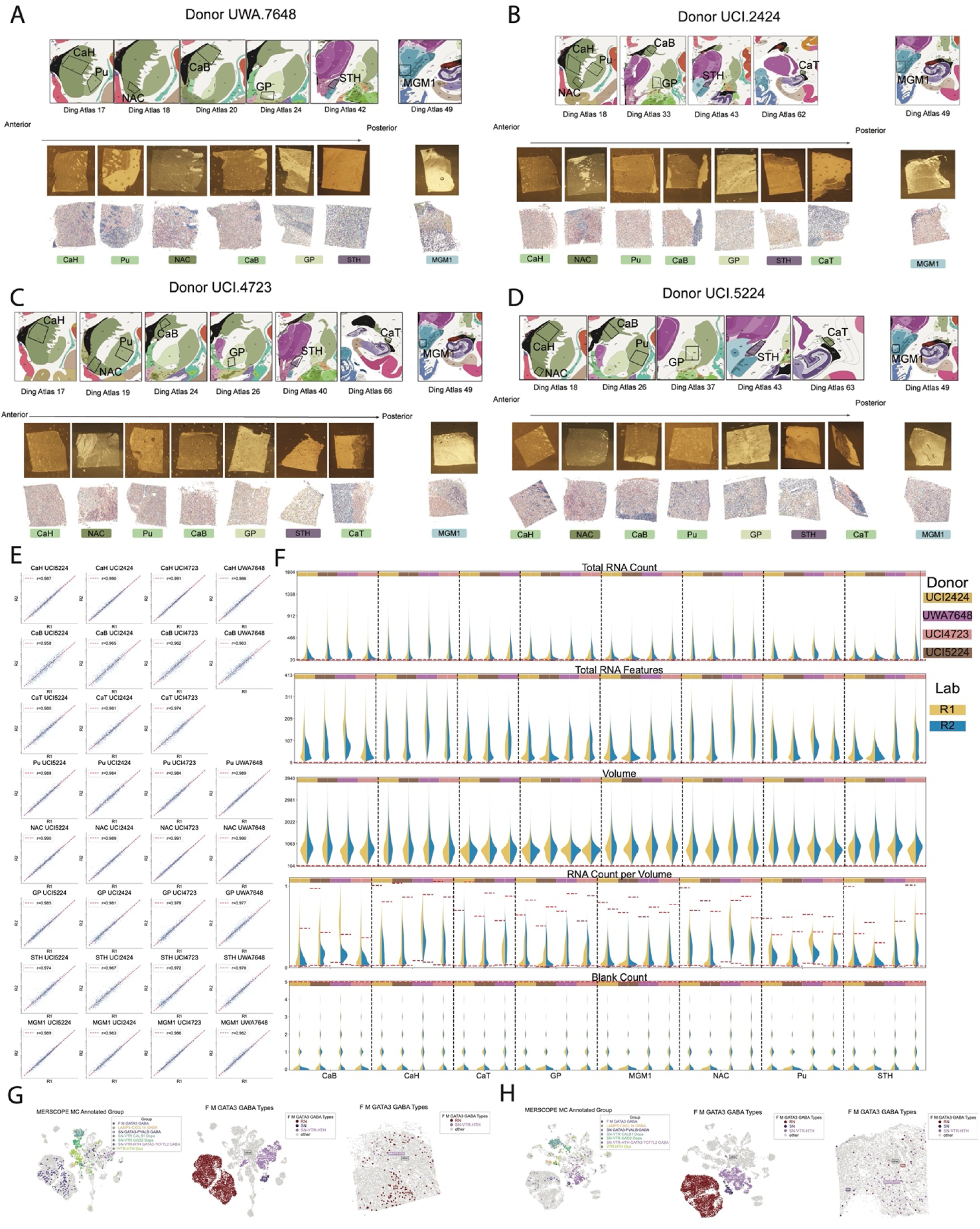
Sampling plan, quality control metrics, and RN GATA3 group in the snm3C-seq atlas corresponds to a red nucleus (RN)-derived cell type. (A-D) Sampling plan for donors UWA7648 (A), UCI2424 (B), UCI4723 (C), UCI5224 (D). The top row shows aligned dissection regions in the Ding Atlas, organized from left to right along the Anterior-to-Posterior axis. The middle row shows images of embedded tissue before MERFISH. The bottom row shows the cell type annotations on the group level of the given sample. The middle and bottom rows show the Salk-specific samples, but all the lab replicates came from adjacent 10 μm segments within the same dissection block. (E) Pearson correlation coefficients for the total number of detected transcripts per gene across all 31 replicates in the 2 labs, with columns representing donors and rows representing brain regions. (F) Violin plots for QC metrics following cell segmentation, separated out by donor, brain region, and lab. From top to bottom showing Total RNA count per cell; Total number of unique genes captured per cell; Cell volume; RNA transcript density within each cell; Number of False positives identified per cell. Horizontal red lines indicate cutoffs used for each QC metric. (G-H) Showing methylation integration results for UCI4723 - MGM1 - R1 (G) and UCI2424 - MGM1 - R1 (H). Each panel is showing (from left to right): the methylation-assigned annotations of the MERFISH cells on the integrated UMAP, the F M GATA3 GABA subclass subtypes on the integrated UMAP, the spatial cells colored by their methylation-assigned annotations, with all cells not in the F M GATA3 GABA subclass labeled as “other”. (G) is a sample whose MERFISH dissection contained the red nucleus, and (H) is a sample whose MERFISH dissection did not. The names of the subclass in the F M GATA3 GABA subclass were abbreviated for brevity in this figure, the name mapping to the taxonomy at the group level is as follows: SN corresponds to ‘SN GATA3-PVALB GABA’, SN-VTR-HTH corresponds to ‘SN-VTR-HTH GATA3-TCF7L2 GABA’, and RN corresponds to RN GATA3 GABA.

**Figure S4.**
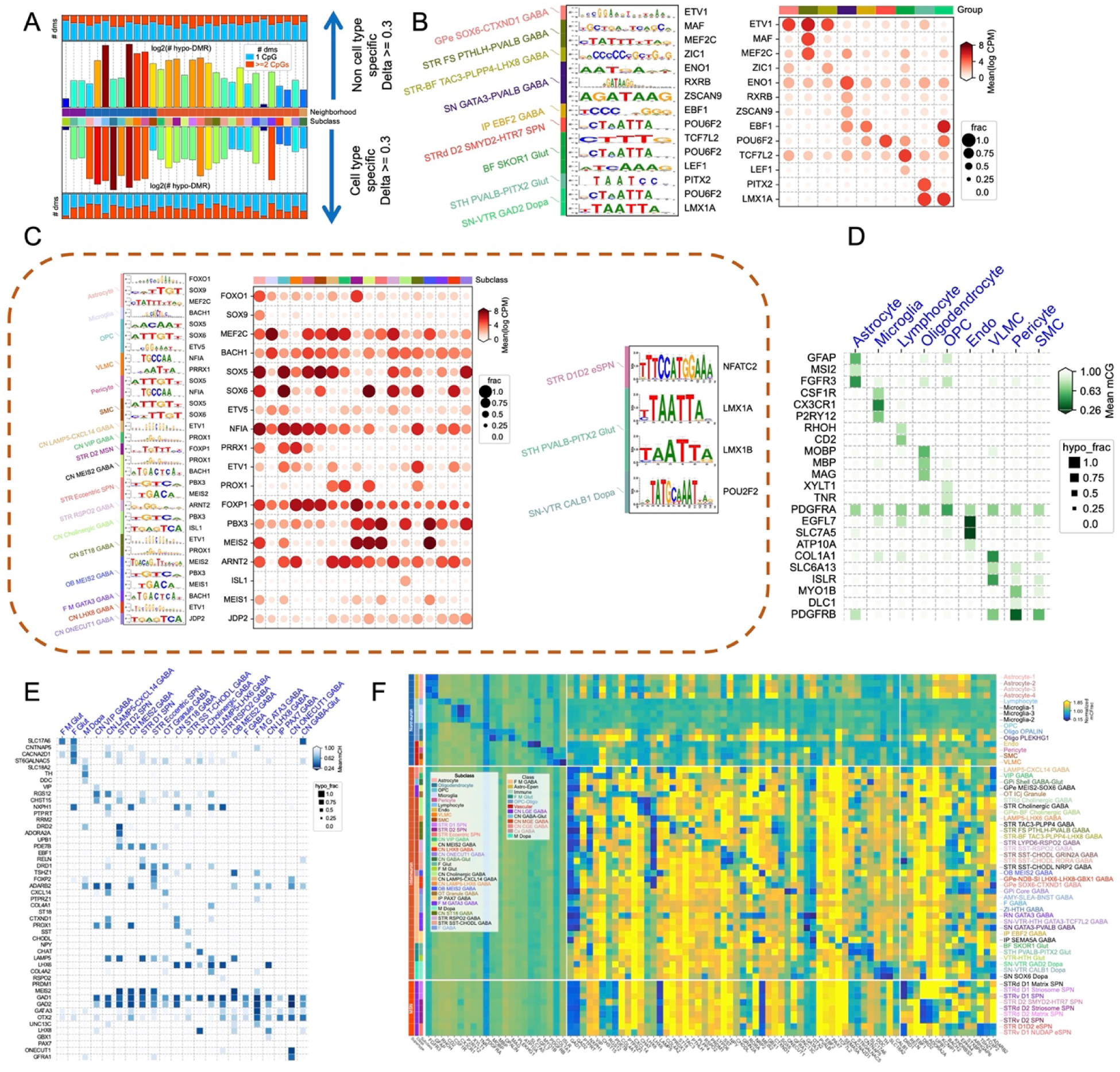
DMR characteristics, TF motif enrichment, and hypomethylation of marker genes. (A) Comparison of cell type-specific versus non-cell type-specific DMRs by the number of differentially methylated sites (DMS) per cell type. (B) Group-level TFBS motif enrichments in hypo-DMRs with corresponding transcription factor expression (upregulation) in the matched cell types. (C) TFBS motif enrichment within hyper-DMRs at subclass and group levels. (D) CG methylation of subclass-level marker genes in non-neuronal cell types. (E) CH methylation of subclass-level marker genes in neuronal cell types. (F) Gene-body methylation of group-level marker genes, shown as mCH for neuronal populations and mCG for non-neuronal populations.

**Figure S5.**
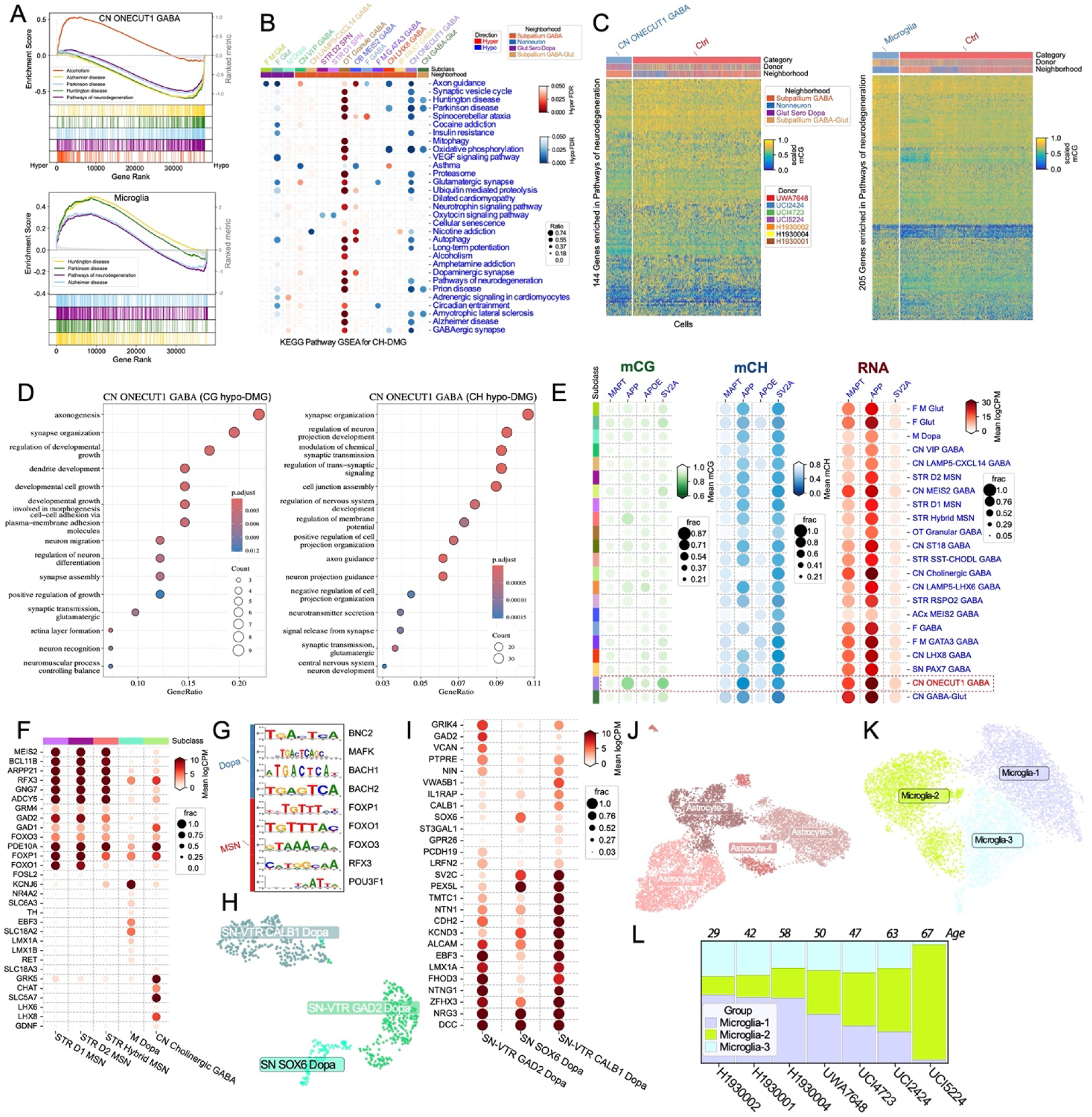
Pathway enrichment, neurodegeneration-related methylation signatures, and cell-type-specific hypomethylation patterns. (A) Representative Gene Set Enrichment Analysis (GSEA) plots showing significant enrichment of neurodegenerative disease- and alcoholism-related pathways in microglia and CN ONECUT1 GABA neurons. (B) KEGG pathway GSEA results for cell-type-specific CH DMGs across subclasses. (C) Heatmap of mCG methylation levels for genes contributing to neurodegeneration-related pathways in CN ONECUT1 GABA neurons and microglia, compared against a control set of randomly selected genes from other cell types. (D) GO Biological Process over-representation analysis of CG and CH hypo-DMGs specifically in CN ONECUT1 GABA neurons. (E) Heatmap illustrating hypomethylation (mCG and mCH) and corresponding gene expression levels of key neurodegeneration-associated genes (MAPT, APP, APOE) and the synaptic vesicle protein *SV2A* in CN ONECUT1 GABA cells. (F) Expression levels of hypo-DMGs across SPN, Dopa, and Chl populations, corresponding to Figure 2G. (G) Hypomethylated TFs with motifs enriched in hyper-DMRs of SPNs and M Dopa neurons. (H) Clustering and annotation of three M Dopa subtypes. (I) Expression levels of hypomethylated DMGs across dopaminergic subtypes, corresponding to Figure 2J. (J) Clustering and annotation of four astrocyte subtypes. (K) Group-level annotation and characterization of microglia subpopulations: UMAP embeddings colored by group label. (L) Enrichment of the microglia-2 subtype in aged donors. Top panel: donor ages; bottom panel: proportion of microglia-2 cells across donors.

**Figure S6.**
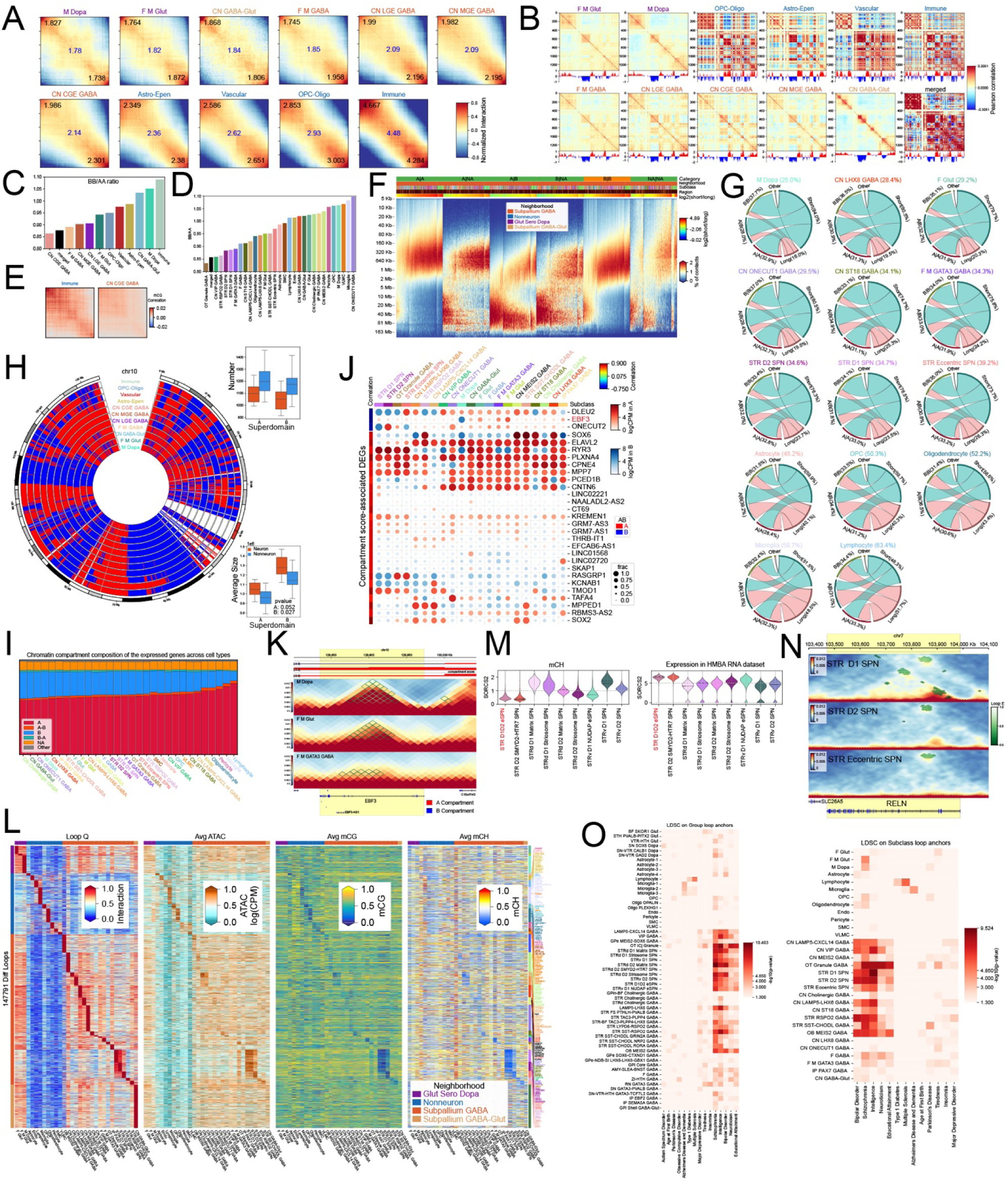
Cell type-resolved A/B compartmentalization and its relationship to contact distance, accessibility, DNA methylation, and gene features. (A) Saddle plots of normalized compartmental interaction frequencies for each subclass; mean B-B (upper left), A-A (lower right), and compartment strength (center) are indicated. (B) Pearson correlation of compartment PC1 on chromosome 10 across different cell types at the class level. (C) Ratio of B-B to A-A compartment interaction frequencies across major cell classes. (D) B-B/A-A interaction ratio (BB/AA) across subclasses. (E) Saddle plot showing mCG correlations in immune cells and CN CGE GABA neurons; the upper-left and lower-right corners correspond to B and A compartments, respectively. (F) Cell type-specific proportions of short- and long-range contacts and of intra- versus inter-compartment interactions. (G) Subclass-level summary of short- vs long-range contacts and intra- vs inter-compartment interactions. (H) Superdomains formed by merging consecutive A or B compartments across classes, with Boxplots summarizing the superdomain number and size for A and B compartments shown on the right. (I) Percentage of genes that are expressed in A, B, and NA compartments across different cell types. (J) Genes exhibiting strong correlation with compartment scores and displaying compartment switching across different cell types. (K) Example loci at *EBF3* locus illustrating compartment switching in M Dopa neurons within the B compartment. (L) Subclass-level correlations between loop interaction strength (10-kb resolution) and chromatin accessibility (ATAC) as well as DNA methylation (mCG and mCH) at anchors of differential loops. (M) CH methylation and Expression of *SORCS2* in SPN subtypes. (N) Example of a subclass-specific differential loop at the RELN locus in STR D1 SPNs. (O) LDSC heritability enrichment results for loop anchors at subclass and group levels.

**Figure S7.**
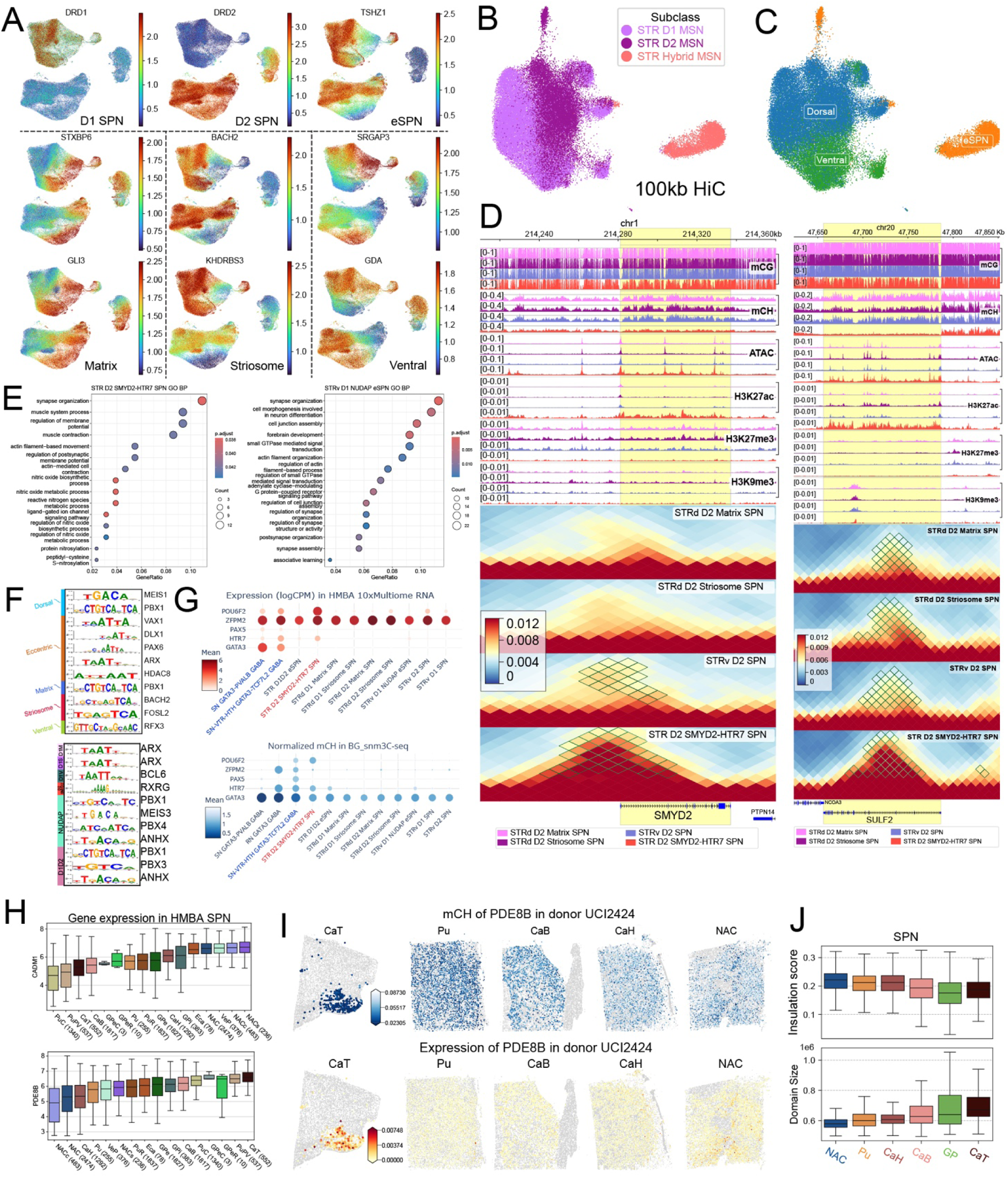
Marker Gene Methylation, Regional Distribution, and 3D Genome Analyses Supporting SPN Subtype Definitions. (A) Gene-body mCH methylation profiles of canonical marker genes utilized for group-level SPN subtype annotation. (B) UMAP embedding of SPNs based on 100-kb resolution Hi-C contact profiles, colored by subclass-level annotations. (C) The same Hi-C-based UMAP as (B), recolored by dorsal, ventral, and eccentric SPN populations, demonstrating their clear separation in 3D genome space. (D) Genome browser views of representative loci (*SMYD2* and *SULF2*) showing hypomethylation, highly accessibility and higher chromatin interaction frequency specific to D2 SMYD2. (E) Over-Representation analysis of cell type-specific CH hypo-DMGs against Gene Ontology (GO) Biological Process terms for D2 SMYD2 and NUDAP subtypes. (F) Top panel: TF motif enrichment analysis in hypo-DMRs underlying major SPN contrasts (dorsal vs. ventral and eccentric; matrix vs. striosome), with candidate regulatory TFs highlighted; Bottom panel: enriched TF motifs and corresponding hypomethylated TFs identified within hyper-DMRs of each SPN group (one group versus all remaining SPN groups). (G) Expression of genes *POU6F2*, *ZFPM2* and *PAX5* across SPN subtypes. (H) Expression of *CADM1* and *PDE8B* across anatomical regions in SPN neurons, derived from the HMBA RNA-seq dataset. (I) Spatial methylation and expression levels of *PDE8B* across anatomical regions in SPN neurons. (J) Boxplots showing the distribution of 3D genome domain sizes and insulation scores across basal ganglia regions in SPN populations.

**Figure S8.**
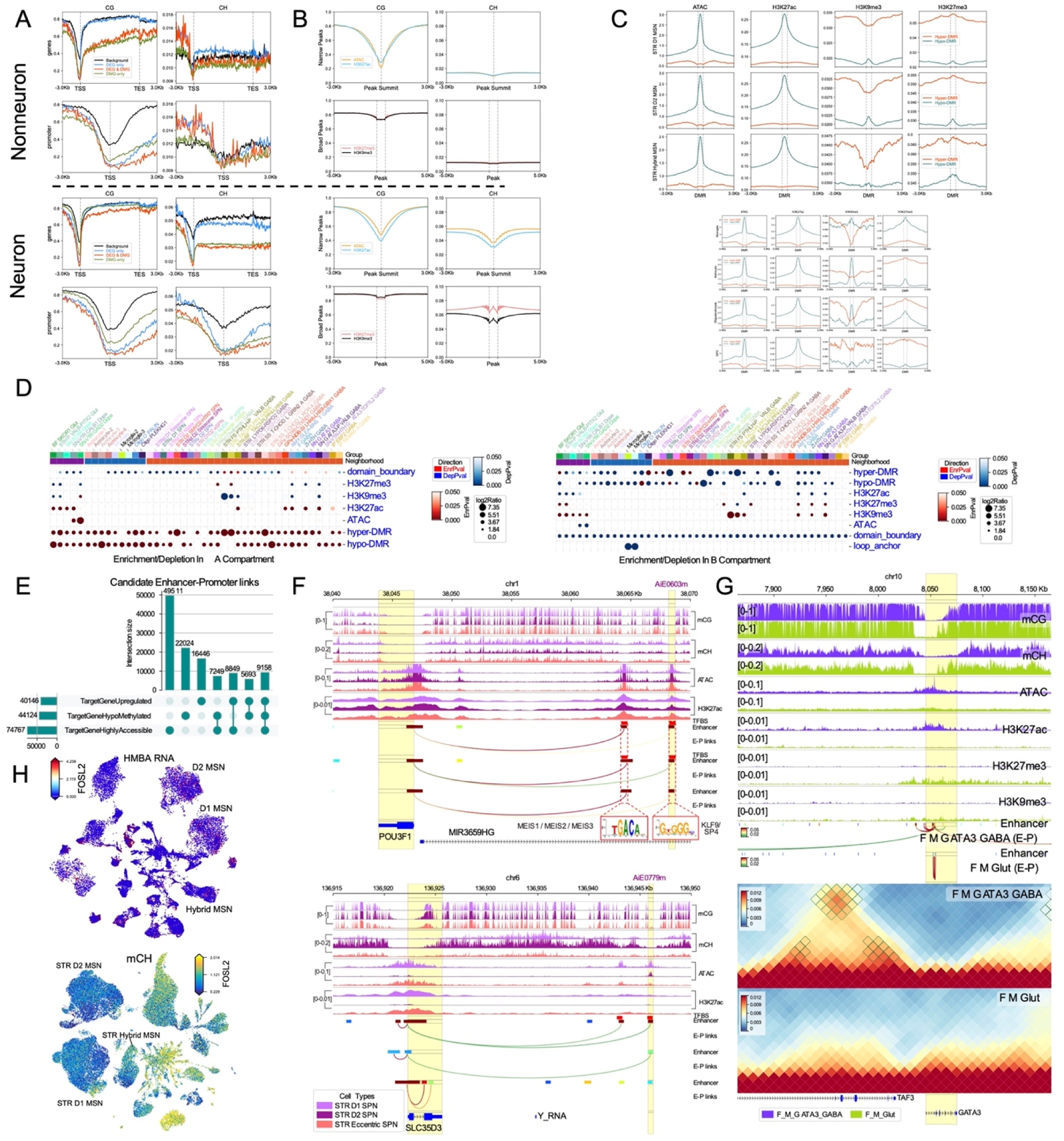
Integrated Epigenomic and 3D Chromatin Profiles with Enhancer-Promoter Interactions and Locus-Specific Genomic Views. (A) Metagene profiles showing the distribution of methylated CG (mCG) and non-CG (mCH) across transcription start sites (TSS) and gene bodies stratified by different categories of gene sets in neuronal and non-neuronal cell types. (B) Distribution of mCG and mCH levels over ATAC-seq peaks and histone modification peaks (H3K27ac, H3K27me3, and H3K9me3). (C) Enrichment of ATAC-seq and histone modification signals over CG hyper- and hypo-differentially methylated regions (hyper-/hypo-DMRs) across SPN subclasses and non-neuronal cell types. (D) Compartment-level enrichment of domain boundaries, ATAC-seq peaks, histone modification peaks, and DMRs in A versus B chromatin compartments. (E) Distribution of target genes exhibiting concurrent upregulation, DNA hypomethylation, and elevated chromatin accessibility. (F) Genomic view of the *POU3F1* and *SLC35D3* loci illustrating additional Activity-By-Contact (ABC)-predicted enhancer-promoter interactions in the corresponding cell types. Purple annotations highlight enhancers experimentally validated in these cell types using the enhancer-AAV toolbox. (G) Example genomic locus showing ABC-predicted enhancer-promoter interactions at *GATA3* in F M GATA3 GABA. (H) UMAP projections of *FOSL2* expression (HMBA RNA-seq) and CH methylation (snm3C-seq).

**Figure S9.**
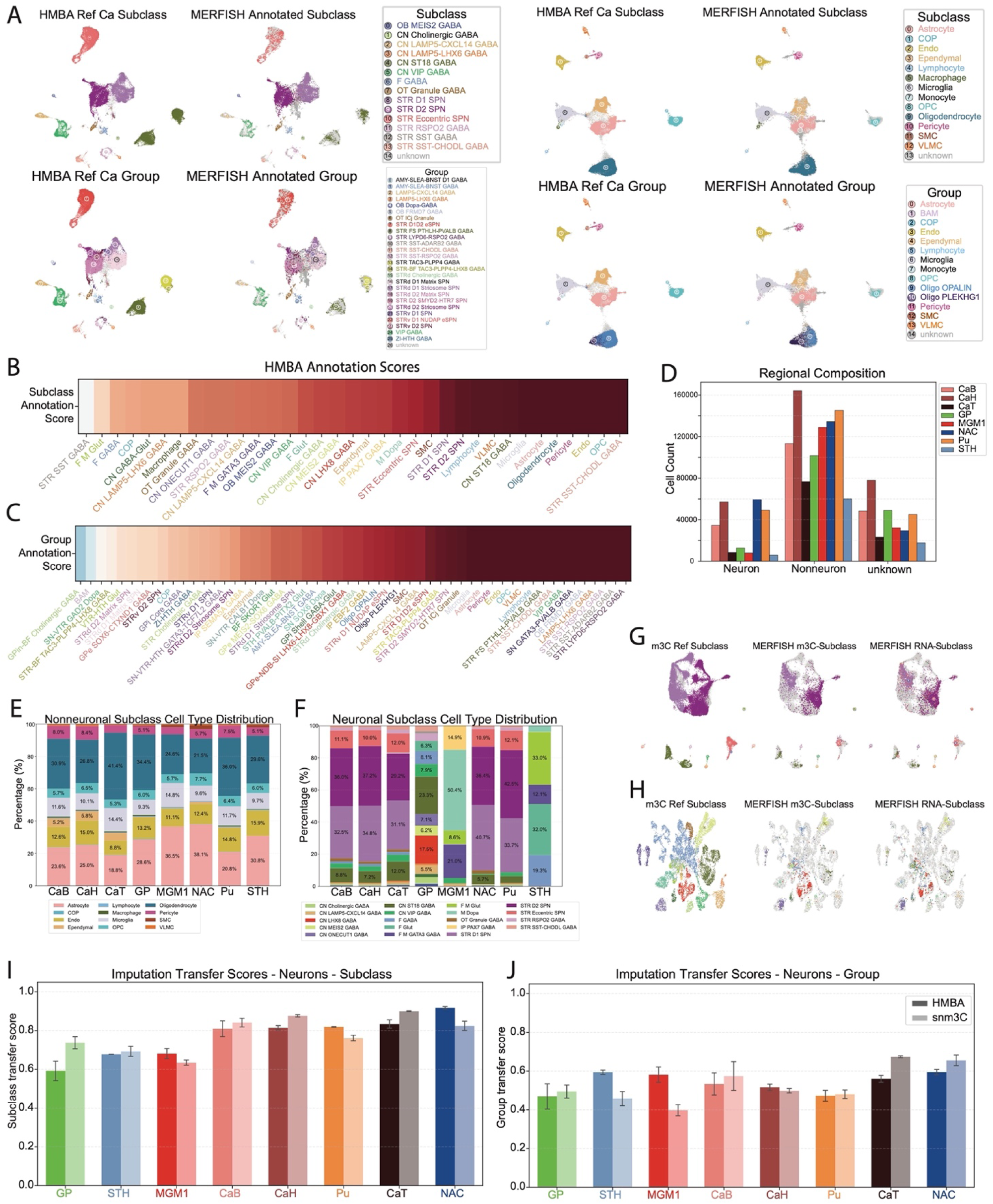
Integration with HMBA scRNA-seq data and snm3C-seq data enables annotation of the MERFISH dataset and imputation of methylation values to space. (A) Example Integration for brain region CaH, donor UCI5224, and replicate 1. 4 subplots each in the shared HMBA x MERFISH shared embedding space. In all subplots the left panel is the reference cells from the region matched HMBA dataset, and the right panel is the MERFISH cells with the assigned annotation. Top left: Neuron subclass integration. Top right: non-neuron subclass integration. Bottom left: Neuron group integration. Bottom right: non-neuron group integration. (B-C) Annotation Score for each cell type at the subclass (B) and group (C) levels. Heatmap showing median annotation scores for a given cell type. Cell-types are ordered by median annotation score. Cell-type names are colored by the cell-type neighborhoods. (D) The count of cells from each major type separated by brain region. Showing neuronal, non-neuronal, and unknown cells. Unknown cells were not reliably called as either neuronal or non-neuronal, and most likely represent low-quality cells that were removed from downstream analysis. Results are grouped together across all donors and replicates for each brain region. (E) Proportions of each non-neuronal subclass across all donors and replicates, split by brain region, colored by cell type. (F) Proportions of each Neuronal subclass across all donors and replicates, split by brain region, colored by cell type. (G-H) Example Methylation Integration for region Pu, donor UCI5224, replicate 1 (G), and region GP, donor UCI5224, replicate 1 (H). 3 subplots in the same MERFISH x snm3C-seq shared embedding space. Left: all reference cells from the region matched snm3C-seq dataset. Middle: MERFISH cells annotated from the snm3C-seq dataset. Right: MERFISH cells with the HMBA derived annotations. (I-J) Integration transfer scores for imputations, split by brain region and reference dataset. Global subclass (I) scores are higher than group (J) scores. HMBA and snm3C scores are highly comparable across regions.

**Figure S10.**
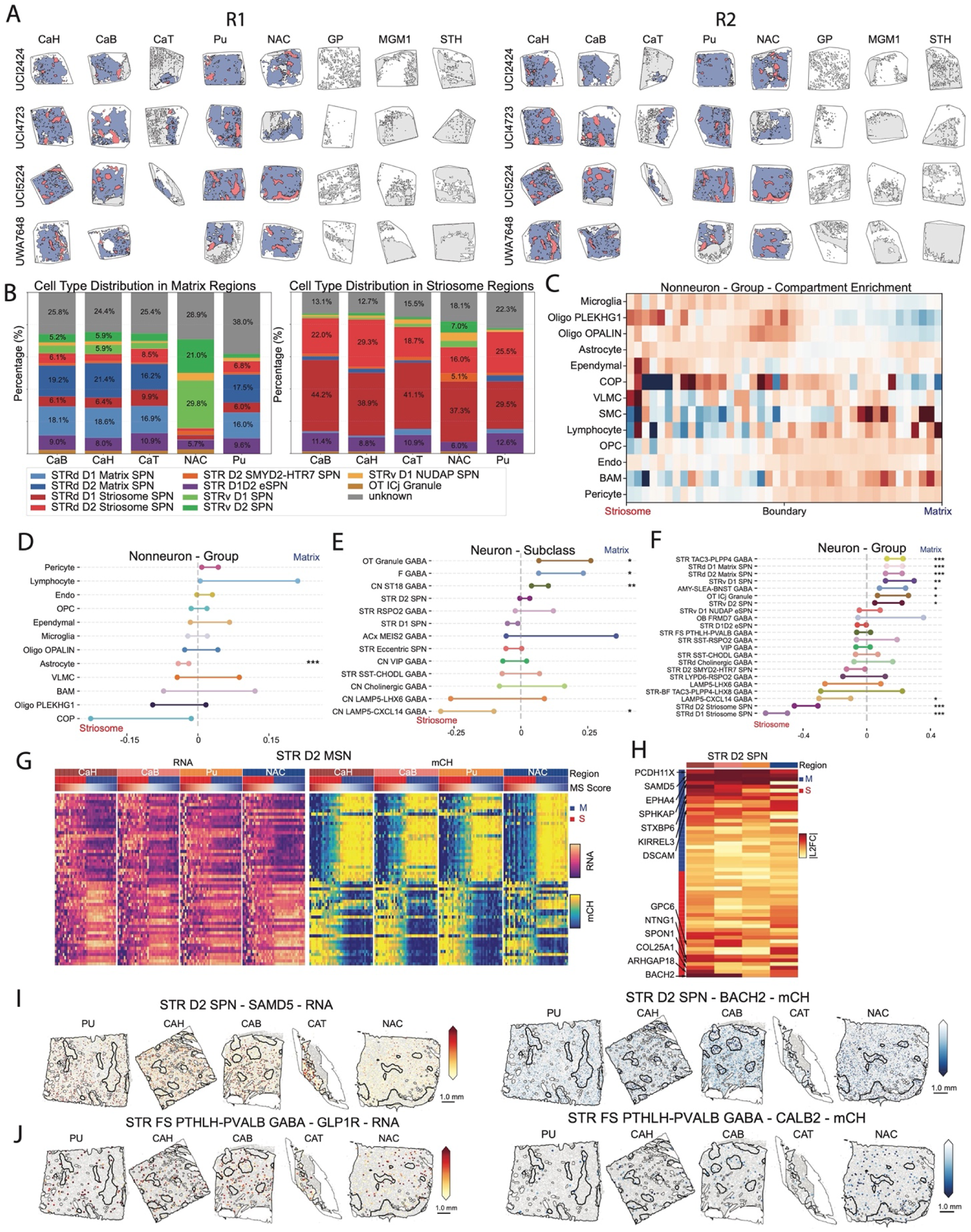
Matrix-Striosome compartment localizations show non-canonical cell type enrichments, with MS-DEGs in STR D2 SPN and STR FS PTHLH-PVALB GABA. (A) Showing all of the segmented MERFISH regions. Regions are labeled as follows: Blue regions represent matrix; Red regions represent striosome; gray regions represent White Matter. All regions not filled in were not identified as any of the above. Results shown for the two replicates. (B) SPN-specific cell-type distribution at the group level within either the called matrix (left) or striosome (right) compartments. Total bar height is normalized to the total number of cells classified as SPNs at the subclass level in the given brain region across all donors and replicates. (C) Distribution of non-neuronal group cell-types along the matrix-striosome gradient. Each row is divided into 40 equal sized bins. Each bin is normalized by the total number of cells in the bin, and then each row is normalized to sum to 1. Bins are colored by that normalized cell density. Red signifies enrichment in those bins. (D-F) Cell-type enrichments in matrix (right) or striosome (left) compartments at the non-neuron group (D), neuron subclass (E), and neuron group (F) levels. Y-axis labels are colored like in (A). The x-axis shows the 95% confidence intervals on the log2 fold-change of the normalized cell-type abundance in matrix and striosome compartments across regions, donors, and replicates. stars indicate FDR-adjusted significance: *<0.01, **<0.001, ***<0.0001. (G) Heatmap of MS-DEGs for the STR D2 SPN subclass. The left panel shows normalized RNA expression, and the right panel shows normalized mCH gene body methylation score. The rows are all MS-DEGs in the STR D2 SPN subclass (Methods). The columns are 40 evenly spaced bins of pseudobulked cells along the MS Score axis. The columns are grouped into brain regions, and within each brain region, organized from left to right by MS Score. (H) Heatmap of the absolute log2 fold change of gene expression in the matrix versus the striosome, stratified by brain region, for the STR D2 SPN subclass. Rows are the MS-DEGs for this subclass, organized by effect size, with the top rows corresponding to matrix expressed genes, and the bottom rows corresponding to striosome expressed genes. Select genes are labeled via arrows. (I) Example spatial distribution of genes in the STR D2 SPN subclass. Only the striosome and matrix is shown. The black outlined regions are striosomes, the rest is the matrix. The gray background is all other cell-types in the regions. Left: measured RNA expression of *SAMD5*, enriched in the matrix. Right: imputed gene body mCH levels of *BACH2*, hypomethylated in the striosome. (J) Same as (I) but for the STR FS PTHLH-PVALB GABA group. Left: measured expression of GLP1R, enriched in the matrix. Right: imputed gene body mCH levels of CALB2, hypomethylated in the striosome in all regions except the NAC.

**Figure S11.**
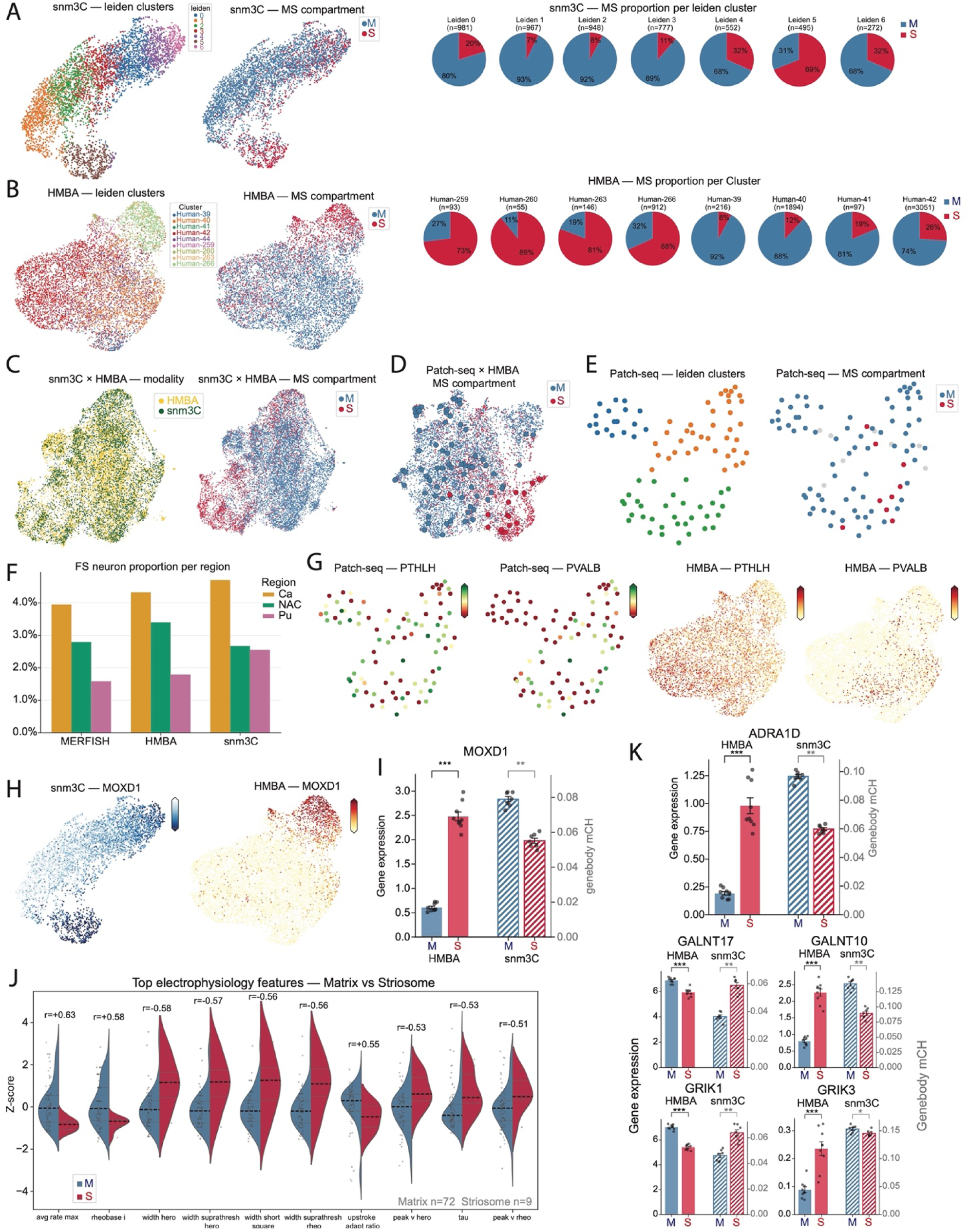
Matrix-Striosome compartment enriched genes in canonical SPN types and novel STR FS PTHLH-PVALB GABA gene enrichment. (A) Clustering of snm3C-seq STR FS PTHLH-PVALB GABA neurons shows striosome (S - red) and matrix (M - blue) enriched subtypes. Left: leiden clusters on umap embedding, Middle: imputed spatial compartment location from MERFISH data on umap embedding. Right: Proportions of each leiden cluster that belong to matrix and striosome. (B) Same as (A) but for the STR FS PTHLH-PVALB GABA neurons in the HMBA dataset. (C) Integration of the STR FS PTHLH-PVALB GABA neurons from snm3C and HMBA together show that the matrix-striosome localization is concordant in both datasets. Left: All cells in shared umap space colored by dataset identity; Right: All cells in shared umap space colored by predicted matrix (M - blue) or striosome (S - red) assignment. (D) Integration of STR FS PTHLH-PVALB GABA neurons from HMBA dataset with the same population from the macaque Patch-seq data. Smaller cells are from HMBA, bigger outlined cells are from Patch-seq. HMBA cells are colored by predicted localization from panel (B). Matrix (M - blue). Striosome (S - red). Patch-seq cells are colored by predicted localization via label transfer from the HMBA cells. (E) Clustering of the STR FS PTHLH-PVALB GABA neurons in the Patch-seq data. Left: Colored by leiden assignment. Right: colored by concordant localization from snm3C and HMBA label transfer. Matrix (M - blue). Striosome (S - red). (F) Regional enrichment of STR FS PTHLH-PVALB GABA neurons across 3 datasets with such information shows expected enrichment in the caudate (Ca) as opposed to the putamen (PU). Bar height is the percent of STR FS PTHLH-PVALB GABA cells within that brain region. (G) Visualizing *PTHLH* and *PVALB* gene expression in STR FS PTHLH-PVALB GABA neurons on the Patch-seq and HMBA cells shows a *PVALB*+ and *PVALB*- population. From left to right: *PTHLH* expression in Patch-seq embedding space, *PVALB* expression in Patch-seq embedding space, *PTHLH* expression in HMBA embedding space, *PVALB* expression in HMBA embedding space. The embedding space for Patch-seq is the same as in S11E. The embedding space for HMBA cells is the same as in S11B. (H) *MOXD1* hypomethylation and increased expression marked the striosome-localized STR FS PTHLH-PVALB GABA neurons in both snm3C and HMBA datasets. Left: gene body mCH of *MOXD1* in the snm3C embedding space. Right: log norm CPM of *MOXD1* in the HMBA embedding space. Embedding spaces are the same as in S11A-B. (I) Barplots showing the increased expression (HMBA - left) and mCH hypomethylation (snm3C - right) of MOXD1 in striosome (S - red) vs. matrix (M - blue). (J) Split violin plots of electrophysiological differences between the predicted matrix (M - blue) and striosome (S - red) localizing STR FS PTHLH-PVALB GABA neurons in the Patch-seq data. (K) Same as (I) but for different genes. From top to bottom and left to right showing: *ADRA1D*, *GALNT17*, *GALNT10*, *GRIK1*, *GRIK3*. Stars indicate significant: *<0.05, **<0.01, ***<0.001.

